# WDR76 promotes MLL-rearranged leukemia via selective recognition of 5-hydroxymethylcytosine in DNA

**DOI:** 10.1101/511394

**Authors:** Kathryn E. Malecek, Hengyou Weng, Matthew A. Sullivan, Claire Y. Kokontis, Michael S. Werner, Jianjun Chen, Alexander J. Ruthenburg

## Abstract

Although rare, the distribution of the 5-hydroxymethylcytosine (hmC) modification in mammalian DNA is tissue- and gene-specific, yet distinct from its transcriptionally-repressive methylcytosine (mC) precursor, suggesting unique signaling potential. To examine this possibility, we fractionated mammalian brain extracts to discover binding partners specific for oxidized states of mC. We demonstrate that one such factor, WDR76, is a highly hmC-specific binding protein that modulates gene expression within chromosomal regions enriched in hmC where it binds. We demonstrate direct transcriptional activation of several target genes in mouse embryonic stem cells as a function of hmC levels and contingent upon WDR76. In human cell lines and mouse models, WDR76 recruitment by hmC is critical for the initiation and maintenance of MLL-rearranged leukemias. Beyond its canonical role as an intermediate in mC remediation, we show that hmC can be an epigenetic mark whose recognition drives leukemogenesis, portending analogous signaling pathways for other rare DNA modifications.

## INTRODUCTION

Methylation of DNA at the 5-position of the cytosine nucleobase epigenetically regulates gene expression in vertebrates, impacting development, cell identity, disease processes and cognition (Bergman and Cedar, 2013; Day and Sweatt, 2010; Smith and Meissner, 2013). Without altering the DNA’s coding potential, the 5-methylcytosine (mC) mark can recruit protein complexes that stably repress local gene expression (Klose and Bird, 2006). The discovery that the TET family of enzymes can catalyze sequential oxidation of mC to 5-hydroxymethylcytosine (hmC), 5-formylcytosine (5fC), and 5-carboxylcytosine (carC) adducts in the genome (Tahiliani et al., 2009; Ito et al., 2011), raises the possibility that the epigenetic language of covalent modifications to the 5-position of cytosine may be far more complex (Figure 1A). Coupled to consequent DNA repair (He et al., 2011; Shen et al., 2013), or passive dilution in cycling cells (Inoue and Zhang, 2011; Wu and Zhang, 2014), this pathway represents the major enzymatic mC removal mechanism in human cells.

**Figure 1.**
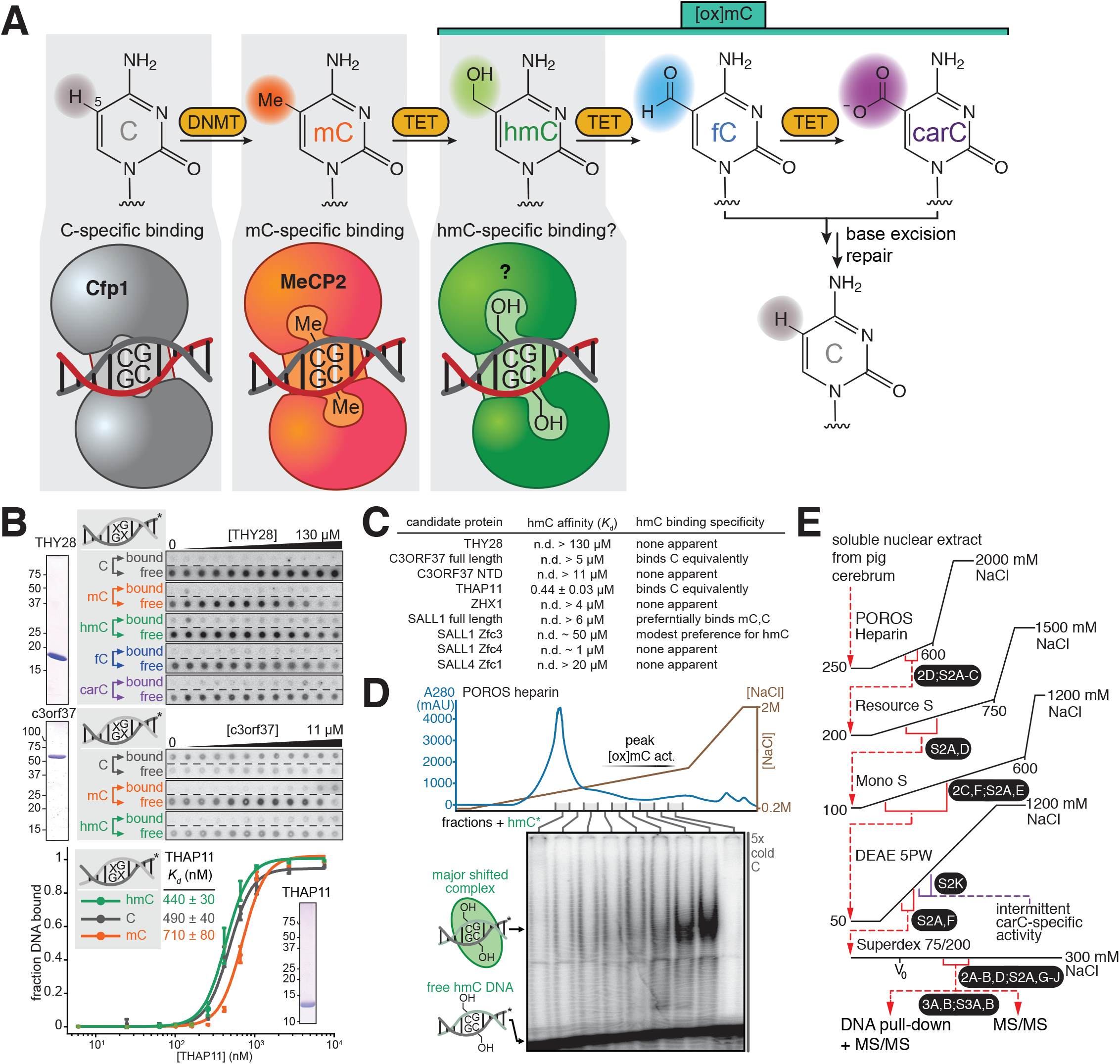
Biochemical purification of the major oxidized-5-methylcytosine-specific binding activities from pig brain. (A) 5-position modification pathways for the deoxycytidine nucleobase (C) within the genome including modification-sensitive binding partners that alter local transcriptional status (Klose and Bird, 2006; Thomson et al., 2010; Wu and Zhang, 2014). DNA methyltransferases install 5-methylcytosine (mC) (Goll and Bestor, 2005)and the TET1/2/3 proteins can sequentially oxidize mC to produce 5-hydroxymethylcytosine (hmC), 5-formylcytosine (fC), and 5-carboxylcytosine (carC) (He et al., 2011; Ito et al., 2011; Shen et al., 2013; Tahiliani et al., 2009). (B) Representative quantitative double filter binding experiments with reported hmC-specific binding factors (Spruijt et al., 2013) reveal lack of specific recognition. Recombinant candidate proteins expressed in expressed in *E. coli* and purified to homogeneity as well as replicates are presented in Figure S1B-H. (C) Summary of affinity and specificity measured in Figure S1 for putative hmC-specific binding factors reported in the literature (Spruijt et al., 2013; Xiong et al., 2016). (D) [ox]mC-specific activities are apparent after POROs heparin fractionation of soluble nuclear extract from porcine cerebrum. Individual fractions indicated are screened in a competitive electrophoretic mobility shift assay (EMSA) to query the binding activity for radiolabeled hmC-modified duplex in the presence of 5-fold molar excess unlabeled competitor duplex of the same sequence but bearing C (“5x cold C”). (E) Optimized chromatography scheme for the isolation of the major [ox]mC-specific DNA-binding activity from nuclear extract of porcine cerebrum; black ovals indicate the figure panels that display representative data at each step.

While modest in abundance in most tissues, hmC accumulates to ~0.7-1.5% of all C in embryonic stem cells and cortical neurons (whereas mC represents ~4% of C in these tissues) (Tahiliani et al., 2009; Kriaucionis and Heintz, 2009;Booth et al., 2012; Globisch et al., 2010; Wen et al., 2014). Individual hmC nucleobases can persist in the genome through several cell divisions in cultured cells and developing mice, with a halflife comparable to that of mC (Bachman et al., 2014). Intriguingly, hmC resides within highly-expressed and lineage-specific genes (Mellén et al., 2012; Song et al., 2012; Tsagaratou et al., 2014) and is enriched in DNA encoding alternative splice junctions and enhancer elements where its mC precursor is not abundant (Pastor et al., 2011; Wen et al., 2014; Yu et al., 2012). The far more rare fC and carC species also exist at somewhat distinct regulatory sites (Wu et al., 2014). Collectively, these data suggest that [ox]mC species, while rare, may not be merely intermediates in the reversion of mC to C. Rather, they may also function as site-specific signaling molecules in the genome distinct from mC. As the TET enzymes play important roles in development (Dawlaty et al., 2014), fertility (Yamaguchi et al., 2012), synaptic plasticity (Rudenko et al., 2013) and cancer (Huang et al., 2013; Huang and Rao, 2014; Rasmussen and Helin, 2016), discerning whether these effects are exerted by loss of mC-based repression, or by orthogonal signaling pathways initiated by specific-[ox]mC binding factors remains a crucial mechanistic question.

We posit a specific chromatin signaling role for [ox]mC modifications by analogy to the well-established epigenetic paradigm of mC (Klose and Bird, 2006) – an information carrying moiety embedded in DNA coupled to cognate binding partners that translate the mark into local signaling output (Figure 1A). Although this hypothesis is not new (Frauer et al., 2011; Iurlaro et al., 2013; Mellén et al., 2012; Shen and Zhang, 2013; Spruijt et al., 2013; Takai et al., 2014; Xiong et al., 2016; Yildirim et al., 2011), evidence for such gene regulatory signaling pathways, in the form of validated binding factors with unambiguous discrimination for particular [ox]mC species, is lacking. Given the scarcity of [ox]mC modifications compared to their far more abundant precursors, binding selectivity sufficient to overcome this disparity is a minimal requirement for distinct genomic localization to be imparted by such a binding event. Searches for binding partners based on similarity to established methyl binding domains (Frauer et al., 2011; Mellén et al., 2012; Yildirim et al., 2011) have proven problematic when subjected to rigorous biochemistry (Hashimoto et al., 2012; Valinluck et al., 2004). More unbiased quantitative proteomics from crude nuclear extracts has provided large lists of potential binding partners with little agreement between three very similar experiments (Iurlaro et al., 2013; Spruijt et al., 2013; Xiong et al., 2016). Moreover, none of these candidates discriminated between chemically distinct [ox]mC species in these experiments (Spruijt et al., 2013), nor have any been demonstrated to bind directly. We attempted biochemical validation of the four strongest candidates from the initial set (Spruijt et al., 2013) that we were able to overexpress in *E. coli* and purify (THY28, HMCEs/C3ORF37, RONIN/THAP11 and ZHX1), but none displayed appreciable specificity for [ox]mC-modified DNA (Figure 1B,C and S1). More recent examples, though coupled with greater functional characterization of the respective proteins, lack convincing biochemical evidence of specificity in the context of the full protein (Takai et al., 2014; Xiong et al., 2016) (Figure S1I-J). Hence, we sought a more rigorous approach using fractionation biochemistry to establish the existence of specific [ox]mC-binding partners as a means to probing the potential signaling roles of these marks.

### RESULTS

#### Discovery and purification of [ox]mC-specific binding factors from pig brain

As [ox]mC species are particularly enriched in adult brain (Globisch et al., 2010; Kriaucionis and Heintz, 2009), we chose porcine cerebrum as a source of soluble nuclear extract for biochemical fractionation to discover, characterize, and purify [ox]mC-specific DNA binding proteins. First, we subjected this extract to heparin chromatography and screened fractions by electrophoretic mobility shift assays (EMSA) using radiolabeled DNA duplexes with a central CpG dinucleotide step bearing modifications at both cytosine nucleobases (Figure 1D and S1A). Through much optimization, we found a sequence of chromatographic steps that can isolate the major apparent [ox]mC-binding activity from others with distinct mobility and specificity properties (Figure 1E and S2A).

The final chromatographic step resolves an activity that specifically binds duplex DNA with hmC and fC but not unmodified C, mC or carC (Figure 2A and B). In a several preparations we also observed a potent carC binding activity that co-fractionated with the major hmC- and fC-specific activity with similar electrophoretic mobility (Figure S2B). As this activity was intermittent from preparation to preparation, we focused on isolating the more reproducible fC- and hmC-specific activity. As another metric of specificity in this native preparation, we examined the sensitivity of the [ox]mC-specific shifts of radiolabeled DNA to different types and concentrations of unlabeled “cold competitor.” We found that all shifted complexes were substantially disrupted by excess fC cold competitor, but not mC or C cold competitors, even midway through the preparation (Figure 2C and S2F). Similarly, cold hmC is a specific competitor of hmC and fC binding (Figure 2C), particularly at later stages of the purification (Figure S2I). Despite resilient binding at high salt concentrations, the shifted complex is non-covalent and is insensitive to supplemented or depleted metals, redox agents, and RNase treatment (Figure S2J and K). Collectively, these data demonstrate that fC and hmC are preferentially bound over C, mC and carC by protein factors in these highly purified fractions.

**Figure 2.**
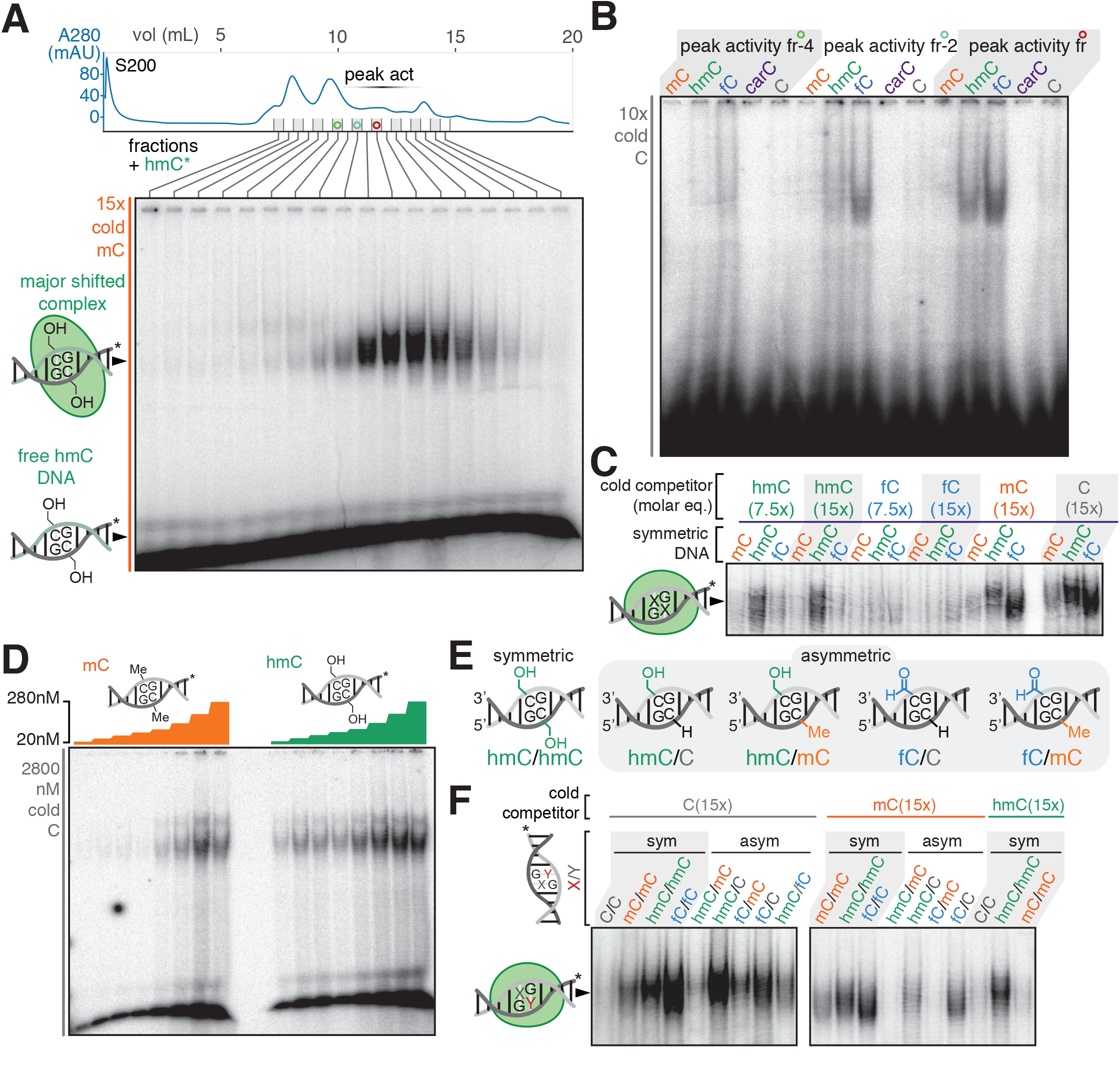
Specificity properties of the major oxidized-5-methylcytosine binding activity. (A) Fractions from the final Superdex 200 column (S200) by EMSA to query binding to a radiolabeled hmC-modified duplex in presence of 15-fold molar excess of unlabeled competitor mC of the same sequence (“15x cold mC”). (B) Three fractions of S200 peak activity screened against the indicated radiolabeled duplexes (symmetric) in the presence of 10 molar equivalents of cold C competitor. (C) After Mono S purification, the shifted complexes with equimolar mC, fC, and hmC DNA exhibit differential sensitivity to excess cold competitors indicated, suggesting fC and hmC are selectively bound, with greater affinity for the former. Only the major shifted complex is presented, the full gel is displayed in Figure S2F. (D) Titration of radiolabeled mC- or hmC- containing duplex from 20-280 nM against late stage protein fraction (Superdex 200) in the presence of 2800 nM unmodified C throughout (10 molar equivalents with respect to the highest concentration point). (E) Schematic of relevant symmetric and asymmetric configurations of 5-position modifications within duplex DNAs. (F) The major activity displays symmetry preferences for [ox]mC/C over [ox]mC/mC that is maintained in subsequent purification steps. Only the major shifted complex is presented, the full gels are in Figure S2D and S2F.

To more quantitatively interrogate specificity in the major hmC and fC-specific activity, a late stage fraction was titrated with increasing concentrations of either symmetric hmC or mC radiolabeled DNA in the presence of unmodified C cold competitor (Figure 2D). Binding is visible at 20 nM hmC DNA (with a 140-fold molar excess of competitor unmodified C), whereas the lowest concentration of mC for which comparable binding is observed is 100 nM mC. This suggests that the factor(s) present in this fraction possess ~five-fold greater affinity for hmC than for mC, and display very little affinity for unmodified C.

Asymmetric presentations of [ox]mC within CpG dinucleotide steps (Figure 2E) may also be relevant genomic targets for binding partners (Yu et al., 2012). We observe asymmetric duplexes with hmC or fC placed opposite mC display markedly attenuated binding relative to their symmetric counterparts, whereas unmodified C is well tolerated in this position (Figure 2F and S2D,F,I).

#### Identification and validation of WDR76 as an hmC-specific binding-factor

To identify the protein(s) responsible for the major observed [ox]mC specific DNA binding activity, we used two parallel mass spectrometry strategies on fractions from the final chromatographic separation. First, we compared the protein composition of the peak of [ox]mC-binding activity versus flanking inactive fractions from two independent preparations that culminated in different size exclusion columns (Figure S3A). Second, we affinity-captured proteins from a pool of the most active size exclusion fractions using immobilized duplex bearing 5-position modifications. We observed protein bands unique to symmetric [ox]mC and asymmetric [ox]mC/C modified DNA (Figure 3A), and excised them and corresponding regions of negative control pull-downs for comparative mass spectrometry. From the collective analysis of 20 solution fractions, 28 pull downs, and 48 gel slices from two independent preparations, we identified several candidate proteins. With one important exception, none of our candidates overlapped with previously reported [ox]mC specific binding proteins (Iurlaro et al., 2013; Mellén et al., 2012; Spruijt et al., 2013; Takai et al., 2014; Xiong et al., 2016; Yildirim et al., 2011). However, we did observe WD repeat protein 76 (WDR76) predominantly within [ox]mC active solution fractions as well as pull-downs using fC and hmC, either symmetric or asymmetrically combined with C (Figure 3B and S3B). WDR76 is among the many candidates reported by Spruijt and colleagues (Spruijt et al., 2013) which prompted us to examine it further.

**Figure 3.**
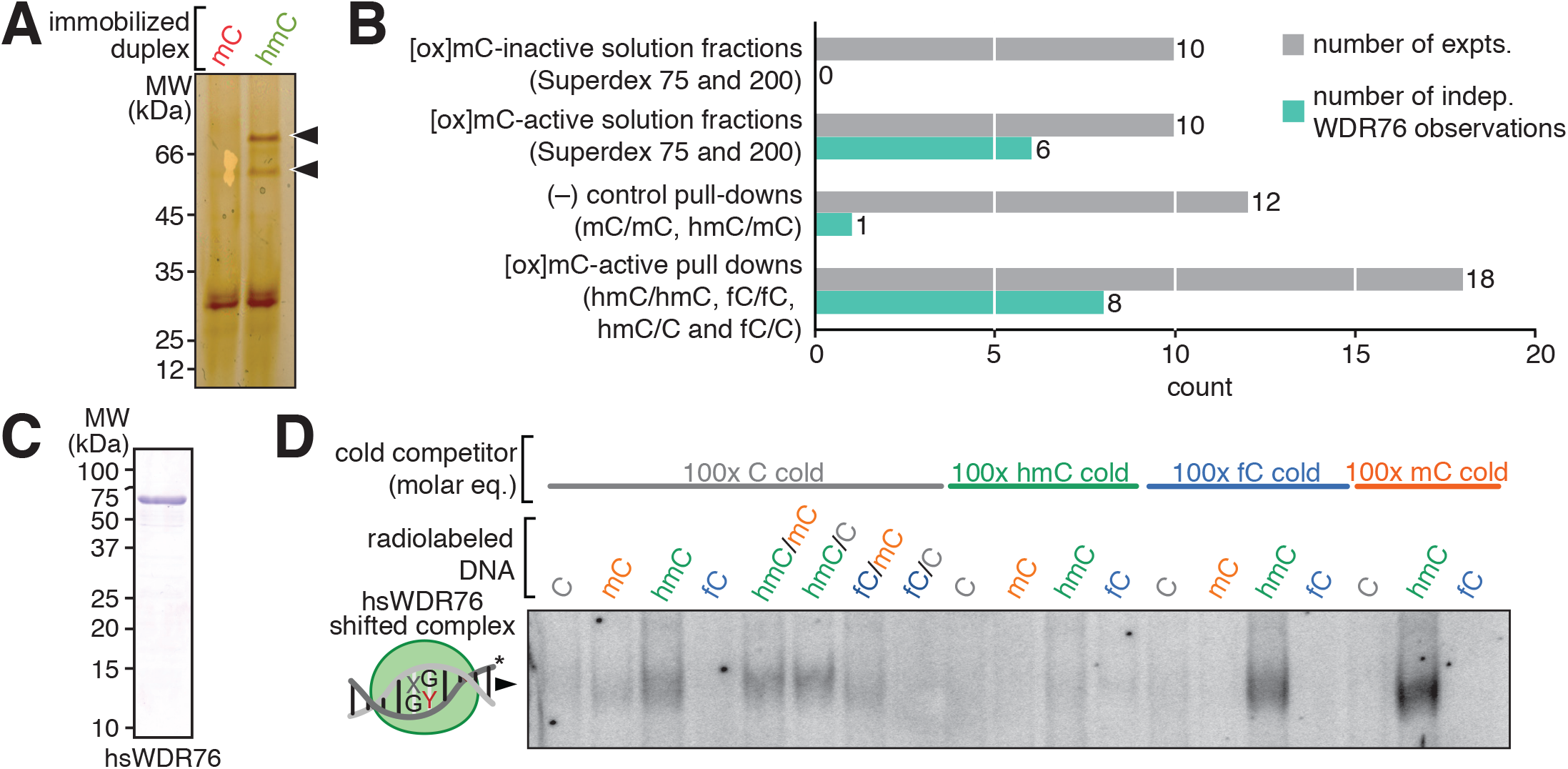
Identification of WDR76 as an hmC-specific binding protein reveals separation of this binding activity from fC-specific binding factors present in the native preparation. (A) A representative silver-stained SDS-PAGE gel comparing indicated biotinylated duplex DNA pull-downs from Superdex 200 fractions. Unique bands excised for identification by mass spectrometry are indicated with black wedges. (B) Candidate protein WDR76 was frequently observed in the active size exclusion fractions and the [ox]mC-specific pull-downs relative to negative controls. (C) Human full-length FLAG-WDR76 expressed by baculoviral transduction of insect cells and purified to apparent homogeneity as assessed by Coomassie-stained SDS-PAGE. (D) EMSA with purified recombinant WRD76 and the indicated radiolabeled oligonucleotides and cold competitors indicates robust hmC-specificity. Only the shifted band with equivalent mobility to the native preparation is presented, see Figure S3C-E for the full gel and related experiments.

We expressed and purified epitope-tagged human WDR76 from baculovirally transduced insect cells (Figure 3C), and subjected it to EMSA (Figure 3D and S3C-E). As with the native preparation, we observed specific binding of WDR76 to hmC, but the purified protein did not discriminate between asymmetric presentations of hmC. More strikingly, we observed no specific binding of WDR76 to fC in any context, nor was 100fold excess cold fC duplex able to perturb hmC-specific binding (Figure 3D and S3C,D). This biochemical separation of function was unexpected given that the hmC- and fC- binding activities co-fractionate over five columns in our native preparation, WDR76 was occasionally detected in fC duplex pull downs, and previous work indicated that WDR76 bound all three [ox]mC states equally (Spruijt et al., 2013). While WDR76 appears to be an hmC-specific binding partner, late stage fractions from our native purification clearly demonstrate the existence of additional factors specific for fC (Figure 2B), carC (Figure S2B), and sensitive to [ox]mC/C asymmetry (Figure 2F) that remain to be characterized.

Our recombinant human WDR76 preparations display low specific activity that further chromatography, alternate construct designs and expression hosts do not effectively remedy (Figure S3F-J), perhaps due to partial misfolding or lack of a critical accessory factor. Although this behavior precludes accurate dissociation constant measurements, semi-quantitative EMSA reveals that WDR76 binds duplex hmC at concentrations as low as 20 nM in the presence of 2000-fold molar excess cold C competitor, whereas comparable apparent mC-complex formation occurs at 100 nM under these conditions, notwithstanding the greater extent of mC radiolabeling (Figure S3E). Given the ratio of genomic hmC to mC in the most enriched tissues is ~1/3 (Booth et al., 2012; Globisch et al., 2010), this degree of preferential recognition of hmC-containing DNA relative to mC and unmodified C could support a distinct epigenetic pathway initiated by specific WDR76 binding.

#### Direct hmC binding by WDR76 potentiates chromatin tethering

As an orthogonal measure of hmC-binding specificity with protein from a different source, we performed duplex oligonucleotide pull downs with extract from HEK293 cells stably transfected with FLAG-HA-tagged human WDR76 (Figure 4A). We observe strong specific binding of WDR76 to hmC, and weak binding of C, mC, hmC/C and fC/C, essentially recapitulating the binding preferences of the isolated recombinant protein. To discern whether WDR76 alone is responsible for these properties or whether another protein in the extract contributes to the specificity, we mutated WDR76 to disrupt its DNA-binding capacity. The C-terminal WD40 repeat region of WDR76 bears sequence and structural homology to the β-propeller domain of DNA damage-sensing protein DDB2 (Figure S4A-C). We selected three basic residues that contact the phosphate backbone of the DDB2 bound DNA (Scrima et al., 2008), and mutated the corresponding conserved residues in WDR76 to serine (Figure S4A-C). This “3S” mutant disrupts binding to hmC-containing DNA (Figure 4B) and modestly reduces the capacity to bind chromatin (Figure 4C and D), indicating that WDR76 is directly responsible for the observed hmC-specific binding, and that this interface contributes to the protein’s chromatin association. Further biochemical dissection of the protein reveals both the WD-repeat domain and a predominantly a-helical domain near the N-terminus both play important roles in DNA binding, although neither module alone accounts for the apparent hmC-biding specificity of the full-length protein (Figure S3H and I).

**Figure 4.**
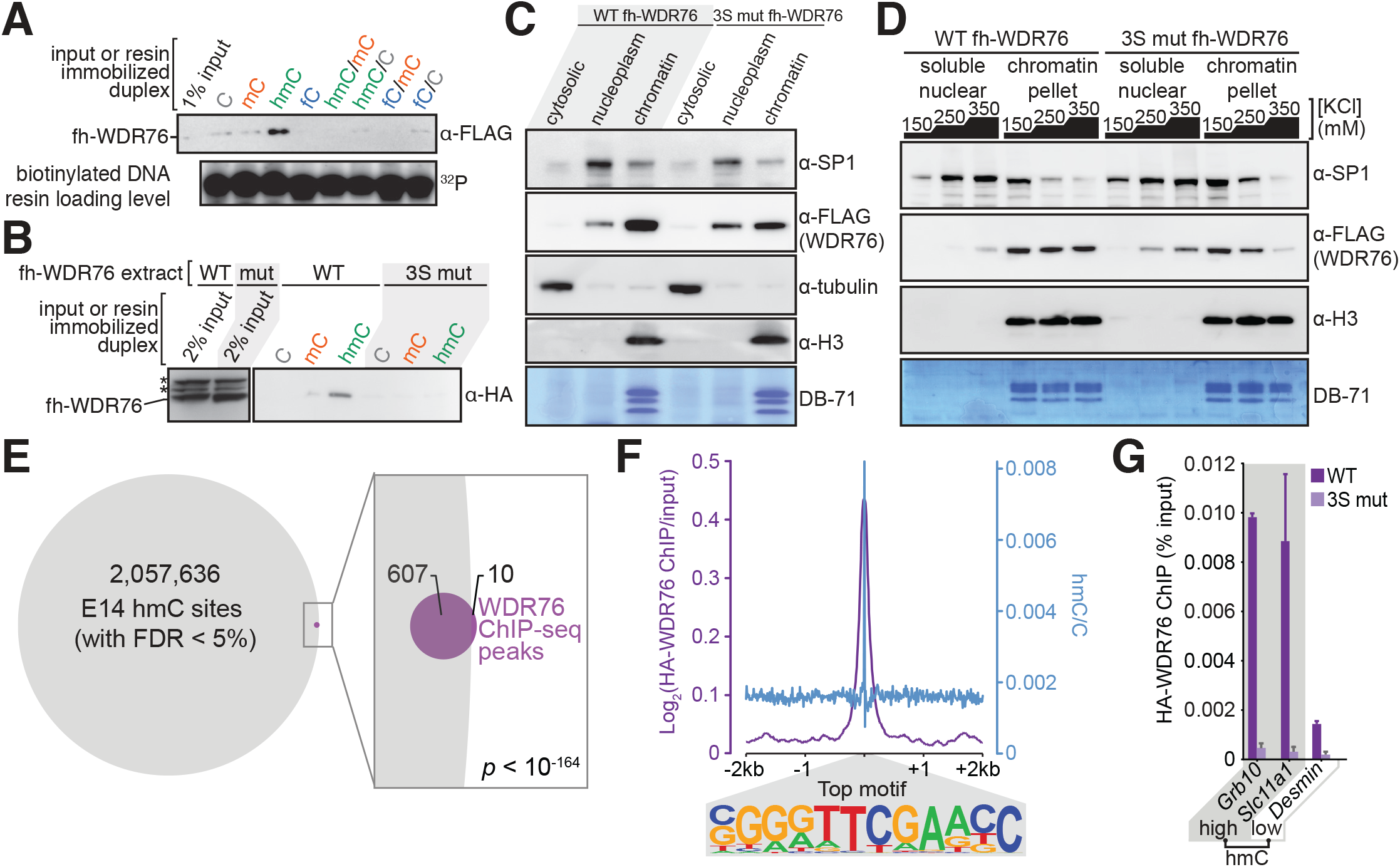
Chromatin engagement by WDR76 in mammalian extracts and cells at a small number of hmC sites depends on its DNA-binding capacity. (A) Pull-down from HEK293 extract expressing human FLAG-HA-WDR76 with the indicated oligonucleotide duplexes imaged by western blot; loading levels of immobilized DNA assessed by post hoc radiolabeling. (B) Mutation of three positively charged residues on the predicted DNA binding interface of WDR76 to serine (3S mut) disrupts DNA binding without altering expression level, solubility, or degradation pattern relative to WT protein in the same experimental format as panel A. Two nonspecific bands in the HA immunoblot input are indicated (*). (C) Subcellular fractionation of HEK293 cell lines stably expressing WT or 3S mutant WDR76 with a nuclear extraction using a 250 mM KCl buffer presented with loading controls for each compartment. (D) Protein extraction of nuclei from WT and 3S mutant WDR76-expressing HEK293 cell lines by parallel extractions with 150-350 mM KCl into soluble nuclear versus chromatin pellet extracts detected by western blot. Transcription factor SP1, histone 3 (H3), and direct-blue 71 (DB-71) staining serve as controls for solubilization of chromatin-bound factors as a function of salt. Tighter chromatin association of the WT protein is reflected in its greater abundance in the nuclear pellet versus the cytosolic extract, even at higher salt concentrations that tend to solubilize weakly bound factors from chromatin adhesion. (E) Venn diagram depicting the overlap greater than expectation (*p* < 10^−164^, two-tailed Fisher exact test) between called HA-WDR76 ChIP-seq peaks (with 600bp extension to reflect library fragment size) and existing high confidence hmC sites in the E14 genome (Yu et al., 2012). (F) ChIP coverage, computed as log2-fold change of depth-normalized HA-WDR76 IP relative to input, contoured over the top sequence motif found in WDR76 ChIP peaks (*p* < 10^−185^, Homer motif enrichment binomial test versus background set, exact motif found in 33.8% of all peaks, 38.2% and 24.8% of all peaks in TADS with genes down- and up-regulated, respectively; see Figure S4I). (G) ChIP-qPCR of WT versus 3S mutant 3xFLAG-HA-hsWDR76 mESC lines (Figure S4E,F) using primers that target hmC-rich sites at the *Grb10* and *Slc11a1* loci, versus an hmC-poor site near *Desmin.*

As specific binding is a crucial element of chromatin modification-based signaling pathways, WDR76 may represent a “reader” for a yet unmapped epigenetic pathway emanating from hmC recognition. While little is known about the function of WDR76, it interacts with chromatin remodeling complexes (Spruijt et al., 2013), and is associated with nucleosomes that bear the chromatin marks of transcriptional activity (Ji et al., 2015). Because hmC is enriched in mouse ES cells (Booth et al., 2012; Stroud et al., 2011; Tahiliani et al., 2009; Yu et al., 2012), we created a stable mESC line with human WDR76 ectopically expressed as an HA epitope-tagged fusion and performed ChIP-seq (Figure S4E-F). From this analysis, we found the vast majority of called peaks overlapped with previously identified high-confidence hmC sites in this cell line (Yu et al., 2012) (Figure 4E and S4G,H), although they are limited to a small subset of largely non-genic sites. It is unclear what accounts for this restriction, although a unique DNA sequence motif centered on a CpG dinucleotide occurs frequently within WDR76 ChIP peaks, and this element is enriched in hmC genome wide (Figure 4F and S4I). The hmC-DNA binding deficient mutant displays reduced ChIP-signal relative to commensurately expressed WT protein at several high hmC-density sites, indicating that an intact DNA-binding interface is important for chromatin engagement (Figure 4G). Although these data indicate that WDR76 engages hmC in mESCs, particularly at sites of an hmC-enriched motif sequence, its function was not immediately obvious from the regions bound.

#### WDR76 regulates transcription within topologically associating domains

To examine potential local gene regulatory roles of the apparent hmC-recruitment of WDR76, we queried the impact of *Wdr76* CRISPR knockout on gene expression by RNA-seq in the same E14 mESC background as our ChIP experiments (Figure S5A-D). Although there were no apparent defects in stemness (Figure S5E), we found both significant loss and gain in expression for several hundred genes in the *Wdr76^-/-^* mESCs relative to the parental E14 cell line (Figure 5A and B), even when normalized to doped internal standards (Figure S5F and G). Intriguingly, the set of altered genes substantially overlaps with a previously defined imprinted gene network that is characteristic of somatic stem cells and important for embryonic growth (Berg et al., 2011; Varrault et al., 2006) (Figure S5H). Genes that display significantly lower expression in the knockout possess greater hmC densities within their gene bodies towards their 3’ ends, relative to up-regulated genes in the knockout, all coding genes, the most highly expressed genes, or a randomly selected genic set of spans of equal length (Figure 5C,D and S5I). Although genic hmC-density generally scales as a function of gene expression (Mellén et al., 2012; Song et al., 2011; Tsagaratou et al., 2014), hmC levels within the bodies of genes that are downregulated in the knockout exceed hmC levels of the most highly expressed genes despite lower expression levels (Figure S5I and J). Normalized WDR76 ChIP signal follows a similar profile across genes (Figure 5E) and is enriched in genes that are down-regulated in the knockout (Figure S5K). Curiously, ChIP peaks were not frequently in the promoters of genes up- or down-regulated in the knockout line. Moreover, annotated enhancers predicted to be connected to genes (Shen et al., 2012) whose expression was significantly altered in the knock-out displayed no clear enrichment of hmC density nor WDR76 ChIP signal (Figure S5L). Consistent with Wdr76 being able to act directly downstream of Tet hydroxylation, there is overlap above chance expectation for genes down-regulated in both Tet1/2/3 triple knockout (Lu et al., 2014) and Wdr76 null mESCs (Figure S5M).

**Figure 5.**
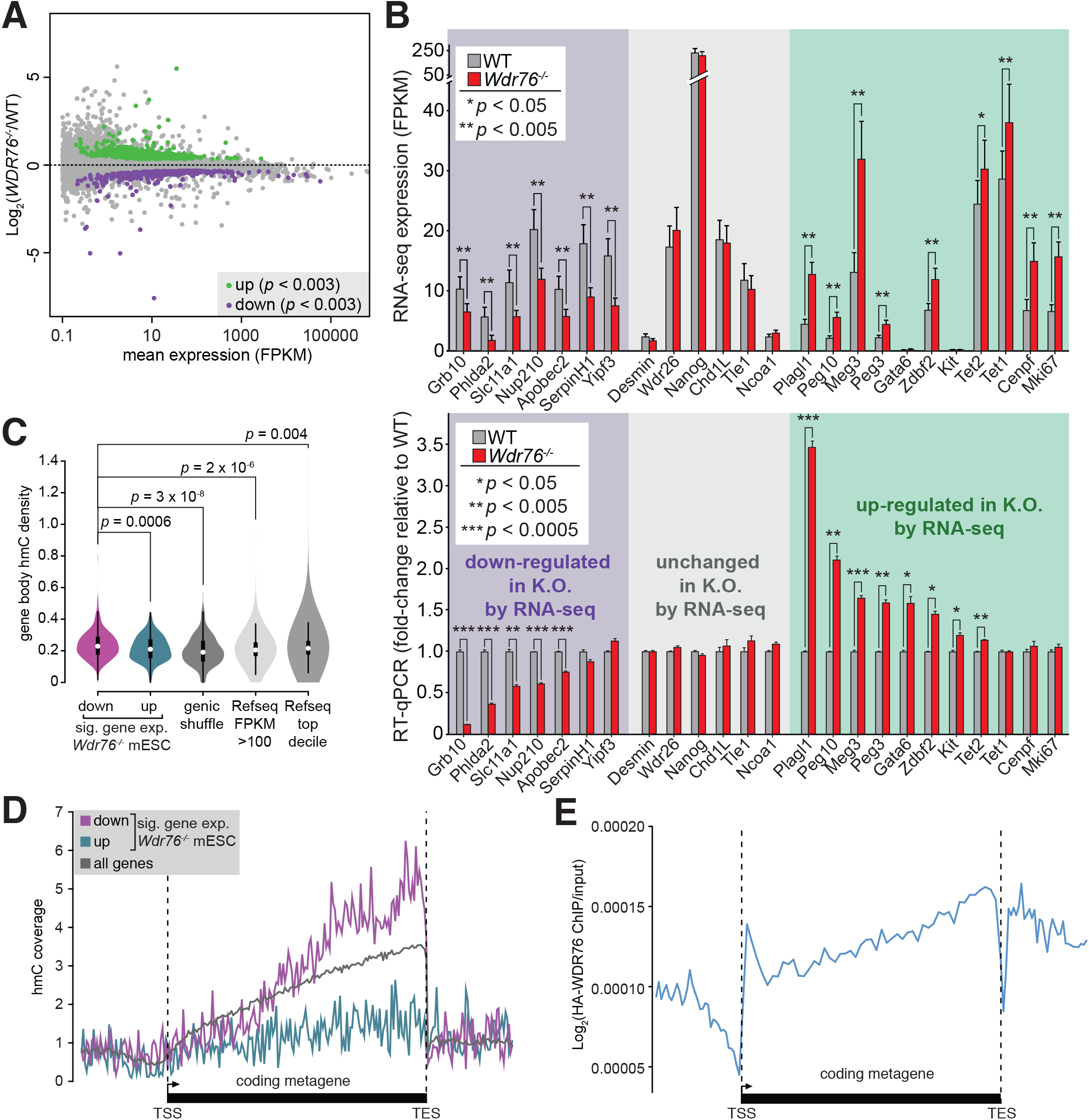
Wdr76 is a transcriptional regulator in mouse ES cells, acting at a set of imprinted genes associated with early mammalian growth. (A) Plot comparing the log2-fold change in Refseq gene expression in *Wdr76^-/-^* versus wt mESCs from RNA-seq. Genes significantly up- or down- regulated in the knockout are depicted in green (488) and purple (359), respectively (n = 3 independent cultures; *p* < 0.003, computed by CuffDiff with Benjamini-Hochberg correction for multiple hypothesis testing). (B) Representatives of three categories of transcripts from RNA-seq are validated by RT-qPCR comparing *Wdr76^−^’’^−^* to wt expression: down- and up-regulated genes, as well as unchanged genes spanning a wide range of basal expression levels. For RNA-seq (upper panel), CuffDiff computed average, uncertainty and p-values are presented (with the Benjamini-Hochberg correction for multiple hypothesis testing). For RT-qPCR (lower panel), error bars are S.E.M. from independent experiment variance, p-values computed by Welch’s two tailed t-test (n>3 independent experiments). (C) Gene body hmC density (% hmC/bp) from all reported E14 mESC sites in TAB-seq (Yu et al., 2012) for significantly down- or up-regulated genes in *Wdr76^-/-^* mESCs versus representative random shuffled gene set representing the same number and spans as the “down” set restricted to other coding genes, all Refseq genes with expression levels greater than 100 FPKM (n = 1038) or the top decile of highly expressed genes in WT E14 mESCs (n = 2459). Central rectangles delimit the upper and lower quartiles with the median indicated by a white dot; p-values computed by Mann-Whitney-Wilcoxon two-tailed test for the overrepresentation of hmC density in the down-regulated genes as compared to the other sets. (D) The abundance of mESC hmC (Yu et al., 2012) contoured over metagenes for the indicated gene sets. (E) WDR76 ChIP-seq coverage over same metagenes.

In recent years, chromosomal regions termed topologically associating domains (TADs) have gained attention as insulated neighborhoods for compartmentalizing genes with the distal regulatory elements that impinge upon them (Dixon et al., 2012; Lieberman-Aiden et al., 2009). We wondered whether there was any connection between TADs that had ChIP peaks for WDR76 and the genes regulated by Wdr76 as ascertained by significant alteration to their expression in the *Wdr76^-/-^* mESCs. Contouring WDR76 ChIP signal over topologically associating domains (TADs) reveals enrichment near TAD boundaries that is especially pronounced in TADS that contain genes that are differentially expressed in the *Wdr76^-/-^* (Figure 6A), and these TADs are more hmC-rich than expectation (Figure 6B and S6A). Remarkably, virtually all TADs that contain genes that are down-regulated in the knockout have significant WDR76 ChIP peaks, as do the majority of TADs bearing up-regulated genes, far above chance expectation (Figure 6C). This phenomenon is recapitulated in a variety of similar analyses with different TAD and ChIP datasets (Figure S6B-D) and is visually apparent at representative loci (Figure S6E).

**Figure 6.**
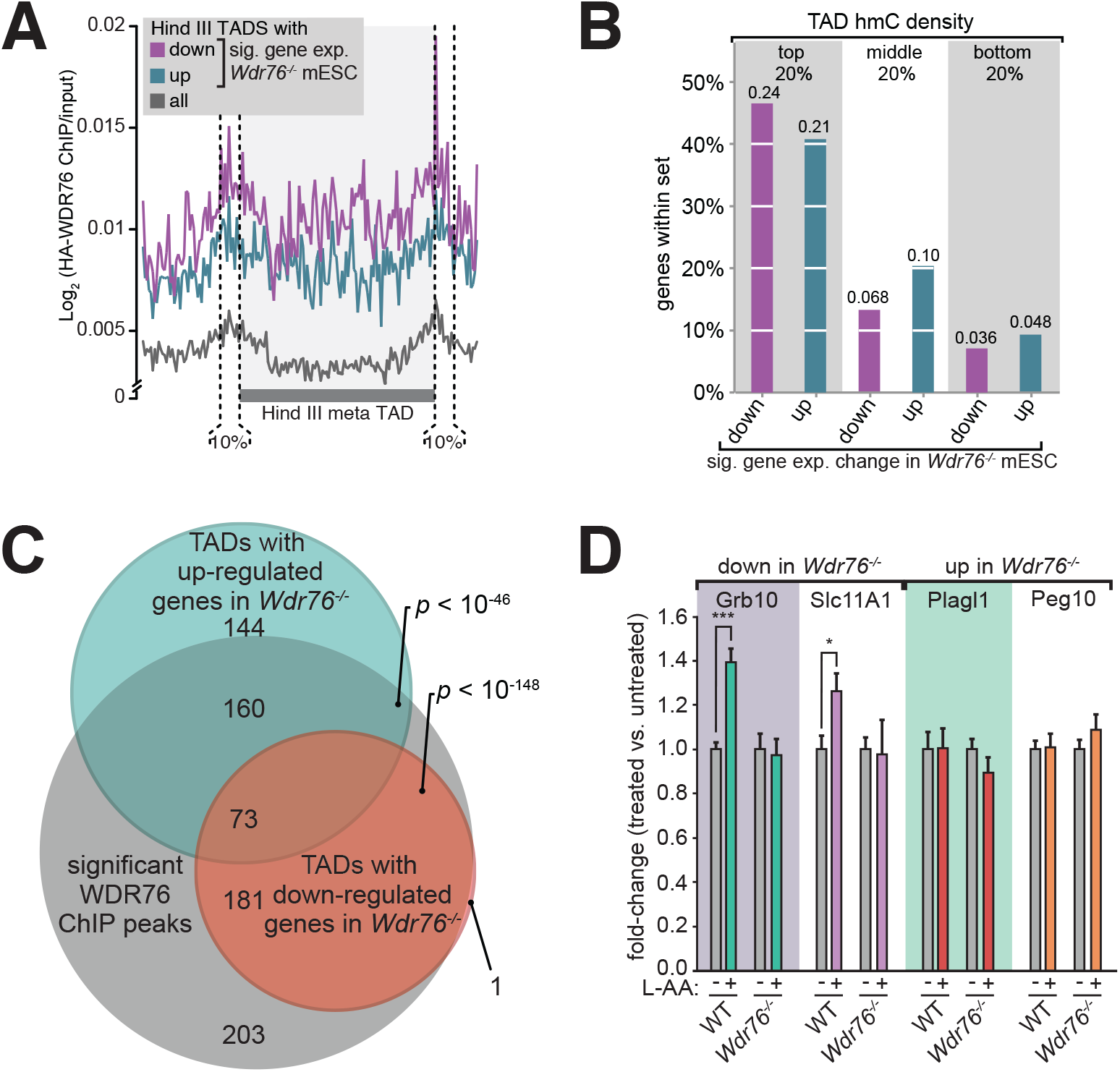
Wdr76 acts within topologically associating domains in mESCs. (A) Log2-fold change of depth-normalized HA-WDR76 IP relative to input, contoured over meta-TADs identified from the consensus of two HindIII Hi-C experiments in mESCs (Dixon et al., 2012), or the subsets of these TADs that bear genes that are up- or down-regulated in the *Wdr76* knockout. As the ChIP signal displays peaks centered on the boundaries, using TAD spans with added 10% cushion more effectively captures the bulk of these peaks, and this cushion was therefore used for follow-on intersection analysis. (B) The percentage of significantly up- or down- regulated genes in the *Wdr76*^-/-^ mESCs within Hind III TADs (Dixon et al., 2012), with the top, bottom, and middle quintiles of hmC-density. The average TAD hmC density for each set is indicated atop each histogram bar. (C) Venn diagram depicting the overlap of mESC topologically associating domains (TADs) harboring genes that are significantly down- or up-regulated in the *Wdr76^-/-^* mESCs with significant WDR76 ChIP-seq peaks. E14 mESC TAD datasets from both mappings (Dixon et al., 2012) are congruent in this comparison, the consensus Hind III dataset is used for this panel and the Nco I is presented in Figure S6D; p-values computed by two-tailed Fisher exact test. (D) Following precedent (Blaschke et al., 2013), stimulation of TETs (and other jumonji-family oxidases) with 100 μg/ml L-ascorbate (L-AA) in WT and Wdr76^-/-^ mESC lines for 12 hours, potentiates the transcription of two genes that are down-regulated in *Wdr76^-/-^*, only in the WT case. RT-qPCR used intron-specific probes to query the pool of pre-mRNA that is actively being transcribed to best expose transcription-level effects on this short-timescale of induction. Error bars represent S.E.M., *p* values are computed by Welch’s two tailed t-test for n =4 independent experiments, \**p* < 0.05, *\*\*\*p* < 0.001. Stimulation of global hmC levels upon L-AA treatment is presented in Figure S6F.

Next, we sought to functionally perturb the enzymes that install hmC on a timescale that minimizes secondary effects on gene expression as a consequence of perturbing 5mC levels (Dawlaty et al., 2014; Wen et al., 2014) to examine the effect on Wdr76-mediated gene expression. Brief L-ascorbate (L-AA) treatments have been shown to markedly stimulate the TET activities and potentiate the global hmC levels in mESCs (Blaschke et al., 2013). While this pleiotropic stimulation of all jumonji-family oxidases is a crude perturbation, it has the advantage over genetic perturbations of all three TET enzymes of being instantaneous. With a 12 hour treatment to elevate hmC levels (Figure S6F), L-AA modestly potentiated Wdr76-depedent expression of two target genes that are down-regulated in both the *Wdr76* and *Tet1/2/3* triple knockouts (Wen et al., 2014), with no discernable change in the expression of two genes that are up- regulated in *Wdr76-/-* (Figure 6D). Yet this potentiation was absent in L-AA treatment of the isogenic *Wdr76-/-* line, suggesting that these transcriptional activation effects are Wdr76-mediated at these two loci, whereas the apparent up-regulated gene set may be indirect Wdr76 targets.

Collectively, our mESC data suggest that WDR76 binds hmC within CpG elements in mESCs, yet some additional binding determinant, perhaps the flanking DNA sequence, restricts this engagement to an extremely small subset of hmC sites. Upon binding, WDR76 plays a role in hmC-driven gene expression in mESCs: within a given TAD with WDR76 ChIP peaks, it may be either a transcriptional activator or repressor, although the latter effect may be indirect. As we observed similar gene regulatory effects within high hmC TADs in the K562 leukemia line upon WDR76 depletion (Figure 7A-C), we sought to further explore the role of WDR76 in leukemogenesis.

**Figure 7.**
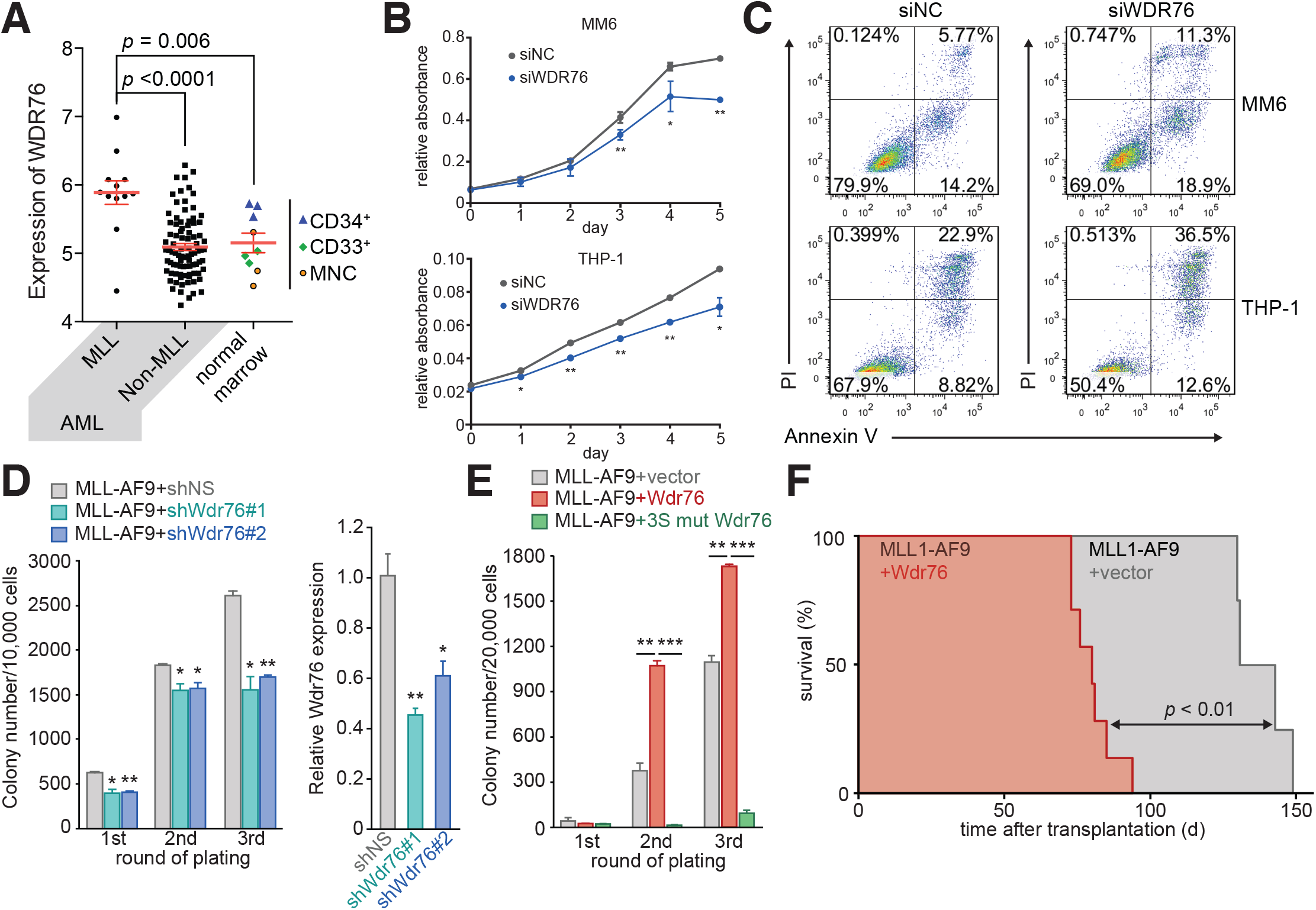
Specific engagement of hmC by WDR76 acts downstream of TET1 in MLL-rearranged leukemogenesis. (A) Microarray measurement of WDR76 transcript levels in human AML patients: 12 MLL1-rearranged, 88 non-MLL1-rearranged, and nine normal bone marrow samples (hematopoietic stem/progenitor CD34+, myeloid CD33+, and mononuclear cell MNC) (Huang et al., 2013). (B) Growth of MLL1-rearranged human leukemia cell lines MONOMAC6 (MM6) and ThP-1 when transfected with siRNA targeting WDR76 relative to negative control (NC). Degree of knockdown is presented in Figure S7B. (C) Apoptosis assays by flow cytometry of Annexin V on the same cell populations as in panel B. (D) Effects of depletion of endogenous Wdr76 by shRNAs (left panel, NS is nonspecific scrambled shRNA as negative control) on MLL-AF9-mediated transformation by colony-forming/replating assays; degree of knockdown by RT-qPCR presented in right panel. (E) Effects of overexpression of WT and 3S (DNA-binding deficient) Wdr76 on the immortalization of MLL-AF9 transduced bone marrow progenitor cells as shown in (D), and expression levels of Wdr76 are available in Figure S7C. (F) Kaplan-Meier survival analysis of the bone marrow transplantation recipient mice. *p* < 0.01; log-rank test.

#### WDR76 promotes MLL-rearranged leukemia by binding hmC

The TET 1 enzyme that installs hmC is required for the initiation and maintenance of MLL-rearranged leukemia, by regulating transcription of critical oncogenic targets of MLL fusions (Huang et al., 2013). This prompted us to examine whether WDR76 acts downstream of TET 1 in this context to promote acute myeloid leukemia. In support, the

MLL-rearranged cohort of 100 human AML patients significantly over-expresses WDR76 (Figure 7A), and the murine homolog, *Wdr76*, is also overexpressed in mouse progenitor cells transduced with several MLL fusions (Figure 7D). Furthermore, depletion of WDR76 by RNAi in two human leukemia cell lines that bear one such rearrangement, the MLL-AF9 translocation, reduces their apparent growth rate with concomitant increases in apoptosis, suggesting WDR76 plays a role in their maintenance (Figure 7B and C; Figure S7E). Modest knock-down of Wdr76 suppresses immortalization of mouse hematopoietic progenitors induced by virally transduced *MLL-AF9* fusion gene in colony-forming/replating assays (Figure 7D). Conversely, co-transduction of a Wdr76 overexpression construct with *MLL-AF9* promotes cell immortalization (Figure 7E and S7F) and results in earlier leukemia onset when engrafted in mice (Figure 7F), with increased lethality traceable to leukemic load (Figure 7G-I). Importantly, this growth acceleration by Wdr76 overexpression is blunted by the 3S mutation that specifically disrupts Wdr76’s DNA-binding capacity. These experiments suggest that hmC is a bona fide epigenetic mark that acts, at least in part, through specific WDR76 engagement to promote the development and maintenance of MLL-rearranged malignancies downstream of TET1 (Huang et al., 2013).

### DISCUSSION

Here we report the discovery WDR76 as a specific 5-hydroxymethylcytosine binding factor by biochemical fractionation, validate its hmC-recognition capacity, establish that WDR76 regulates gene expression by binding this DNA modification in mESCs and demonstrate that these binding events are critical for MLL-rearranged leukemogenesis. Prior reports of putative hmC-binding partners present ambiguous specificity assessments (Iurlaro et al., 2013; Mellén et al., 2012; Spruijt et al., 2013; Takai et al., 2014; Xiong et al., 2016; Yildirim et al., 2011) many of which have been called into question by direct measurements (Hashimoto et al., 2012; Valinluck et al., 2004) (Extended Data Fig. 1), nor have any of their functional effects clearly been ascribed to hmC-specific recognition (Mellén et al., 2012; Takai et al., 2014; Xiong et al., 2016; Yildirim et al., 2011). In contrast, the biochemical specificity we measure is sufficient for WDR76 to selectively localize to hmC in tissues where this modification is abundant. Moreover, this specificity is also manifest in our cell-based functional data that demonstrates hmC-engagement in the regulation of gene expression. Collectively, our results argue that hmC-specific recognition by WDR76 constitutes a bona fide epigenetic pathway for this mark with implications in oncogenesis.

Despite apparent mixed-effects on gene expression in the *Wdr76* knockout in mESCs, all but one of the TADs that contain significantly down regulated genes have at least one WDR76 ChIP peak within them, arguing for the primacy of a transcriptional activation mechanism emanating from hmC enriched sites bound by WDR76. Furthermore, stimulation of TET activity by short exposure to vitamin C (Blaschke et al., 2013), potentiates the transcription of two Wdr76 targets, contingent on the presence of Wdr76. Yet this immediate transcriptional modulation downstream of TET activation only occurs for genes that were down-regulated in *Wdr76^-/-^* (and notably also down-regulated in

TET1/2/3 knockout mESCs (Lu et al., 2014)). Critically, no comparable repression occurs for genes significantly up-regulated in the knockout. This is consistent with the apparent activation in the Wdr76 knockout perhaps being a consequence of indirect effects, while the Wdr76-dependent transcriptional activation observed in this experiment is apparent after a short 12 hours of vitamin C exposure co-incident with the peak of TET stimulation and hmC levels. Consequently, we infer the action of Wdr76 on transcription upon hmC binding is likely to be direct.

Although the events that ensue from specific WDR76-hmC binding have yet to be determined, this work establishes a new paradigm for hmC-directed regulation of proximal genes via specific cognate binding partner to drive the MLL-rearranged class of leukemias. Further research is needed to examine whether this or similar epigenetic pathways based on other specific-[ox]mC-binding factors may be operating during development, learning, and disease states where TET proteins are important (Dawlaty et al., 2014; Huang et al., 2013; Rudenko et al., 2013).

## ACKNOWLEDGMENTS

The authors would like to thank Chuan He and Miao Yu for providing the modified oligonucleotides used in our initial studies. We thank Roderick Davis and Jerry White at the RRC at UIC for technical assistance with mass spectrometry experiments. We would like to thank the following University of Chicago Core Facility personnel: Pieter Faber and Hannah Whitehurst in Functional Genomics for Illumina sequencing, and Don Wolfgeher for assistance with MS data processing. We are grateful to Joseph Piccirilli and Lucia Rothman-Denes for input on the manuscript. A.J.R. is supported by the American Cancer Society (130230-RSG-16-248-01-DMC), Chicago Biotechnology Consortium with support from The Searle Funds at The Chicago Community Trust and Ellison Medical Foundation (AG-NS-1118-13); K.E.M. was supported by NIH (T32 GM007183); M.A.S. and C.Y.K. were supported by the Biological Sciences Collegiate Division Research Endowments at the University of Chicago. J.C. was supported by NIH R01 grants CA178454, CA211614 and CA214965, as well as Leukemia & Lymphoma Society (LLS) Scholar Award.

## AUTHOR CONTRIBUTIONS

H.W. and J.C. designed and performed leukemia studies in human cell lines and mouse model systems. M.A.S. expressed and purified THY28, C3ORF37/HMCES, ZHX1 constructs, contributed to the synthesis, deprotection and purification of modified oligonucleotides, and performed all filter binding assays as well as the ZHX1 EMSA, as well as performed most of the RT-qPCR and associated experiments presented for mESCs including LAA induction of WDR76 target genes. C.Y.K. and M.A.S. performed SALL1/4 expression, purification and binding studies. M.S.W. designed and performed the K562 WDR76 CRISPRi experiment. A.J.R. performed sub-cellular fractionation experiments in HEK293s. All other experiments including the biochemistry and genome-scale experiments in mESCs were performed by K.E.M. A.J.R. and K.E.M. performed analyses and wrote the manuscript.

## ACCESSION NUMBERS

The ChIP- and RNA-seq reported in this paper have been deposited to the Gene Expression Omnibus (GEO) with the accession GSE108832.

## DECLARATION OF INTERESTS

The authors declare no competing financial interests.

