## Supplementary Figures and Methods for "WDR76 promotes MLL-rearranged leukemia via selective recognition of 5-hydroxymethylcytosine in DNA"

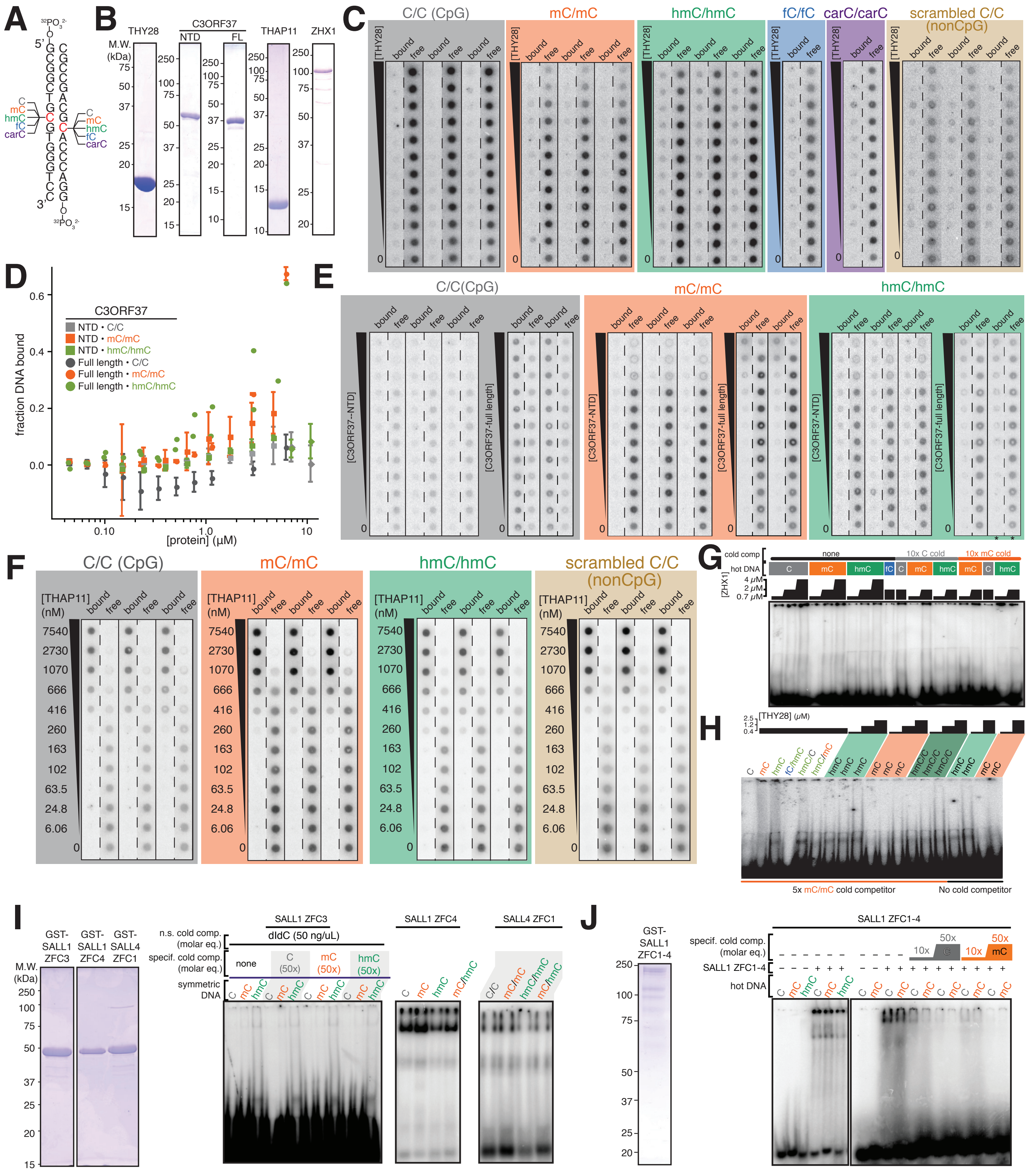

### Figure S1

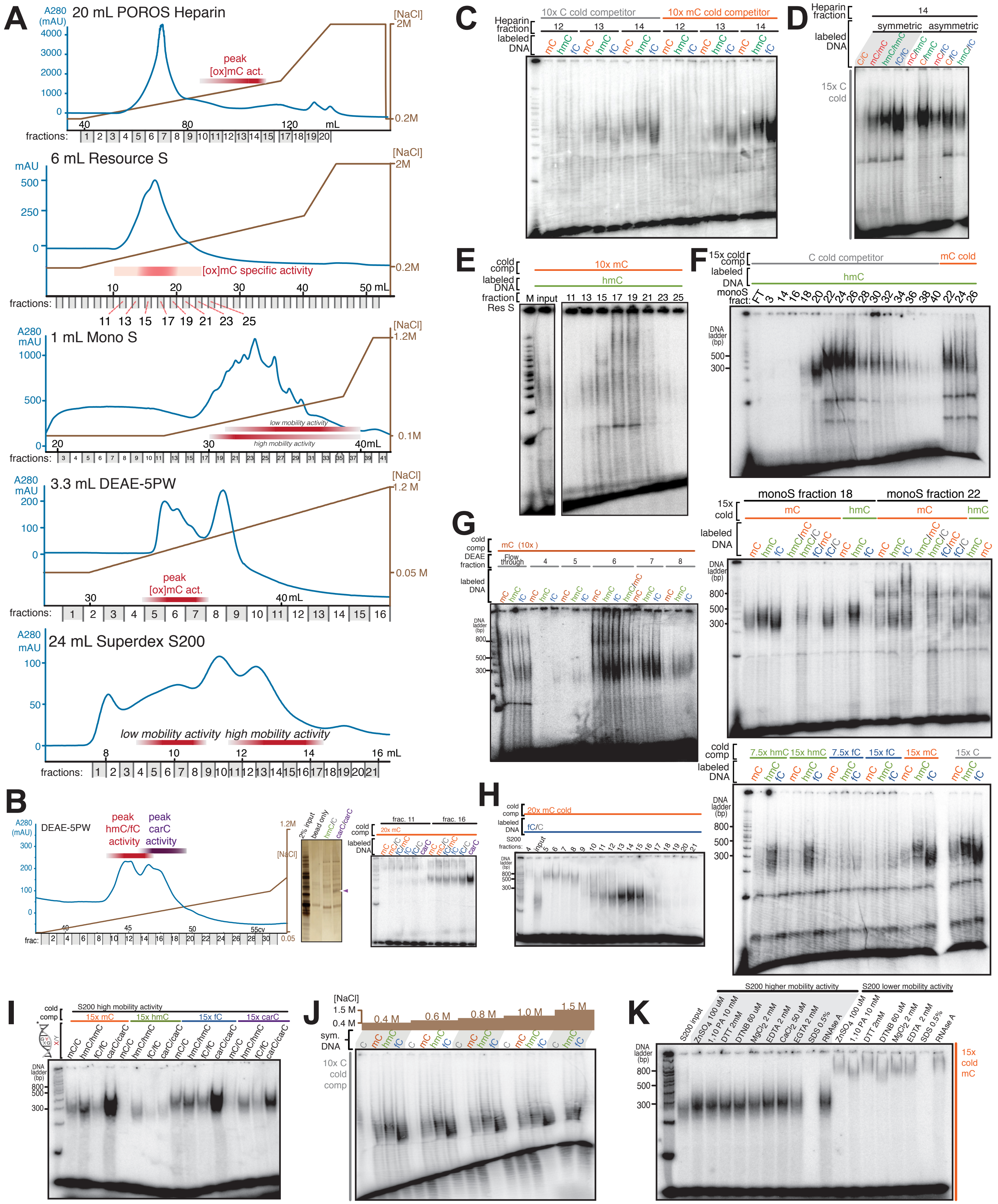
A
20 mL POROS Heparin
peak [ox]mC act.
6 mL Resource S
[ox]mC specific activity
1 mL Mono S
low mobility activity
high mobility activity
3.3 mL DEAE-5PW
peak [ox]mC act.
24 mL Superdex S200
low mobility activity
high mobility activity
B
DEAE-5PW
peak hmC/fC activity
peak carC activity
C
10x C cold competitor
10x mC cold competitor
Heparin fraction
labeled DNA
D
symmetric
asymmetric
15x C cold
E
10x mC
hmC
F
C cold competitor
mC cold
monoS fract:
monoS fraction 18
monoS fraction 22
G
mC (10x)
DEAE fraction
H
20x mC cold
fC/C
S200 fractions:
I
S200 high mobility activity
15x mC
15x hmC
15x fC
15x carC
J
[NaCl]
0.4 M
0.6 M
0.8 M
1.0 M
1.5 M
sym. DNA
10x C cold comp
K
S200 higher mobility activity
S200 lower mobility activity
15x cold mC

### Figure S2

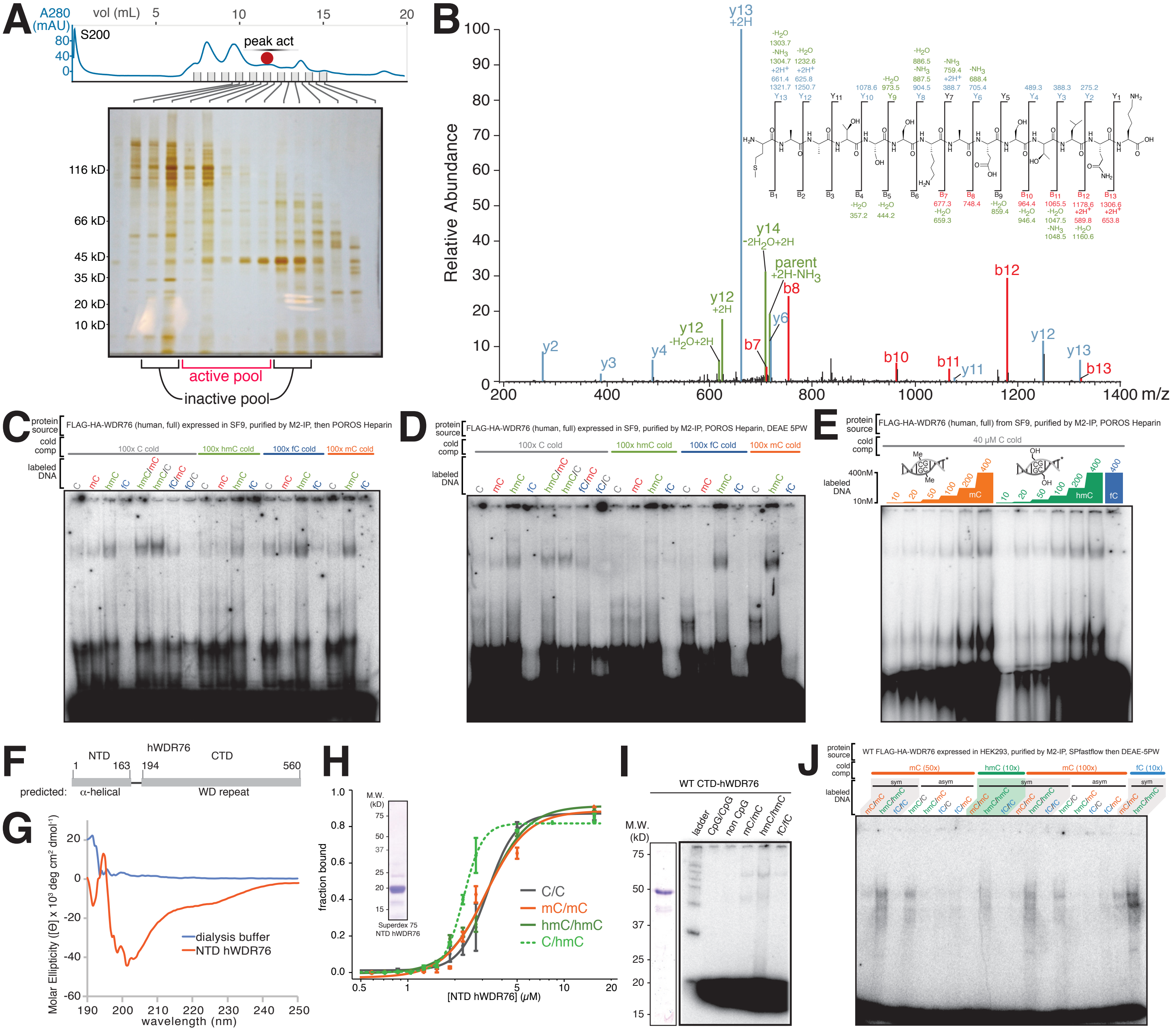

A
B
C
D
E
F
G
H
I
J
active pool
inactive pool
peak act
Relative Abundance
FLAG-HA-WDR76 (human, full) expressed in SF9, purified by M2-IP, then POROS Heparin
FLAG-HA-WDR76 (human, full) expressed in SF9, purified by M2-IP, POROS Heparin, DEAE 5PW
FLAG-HA-WDR76 (human, full) from SF9, purified by M2-IP, POROS Heparin
hWDR76
dialysis buffer
NTD hWDR76
[NTD hWDR76] (μM)
fraction bound
WT CTD-hWDR76
WT FLAG-HA-WDR76 expressed in HEK293, purified by M2-IP, SPfastflow then DEAE-5PW


### Figure S3

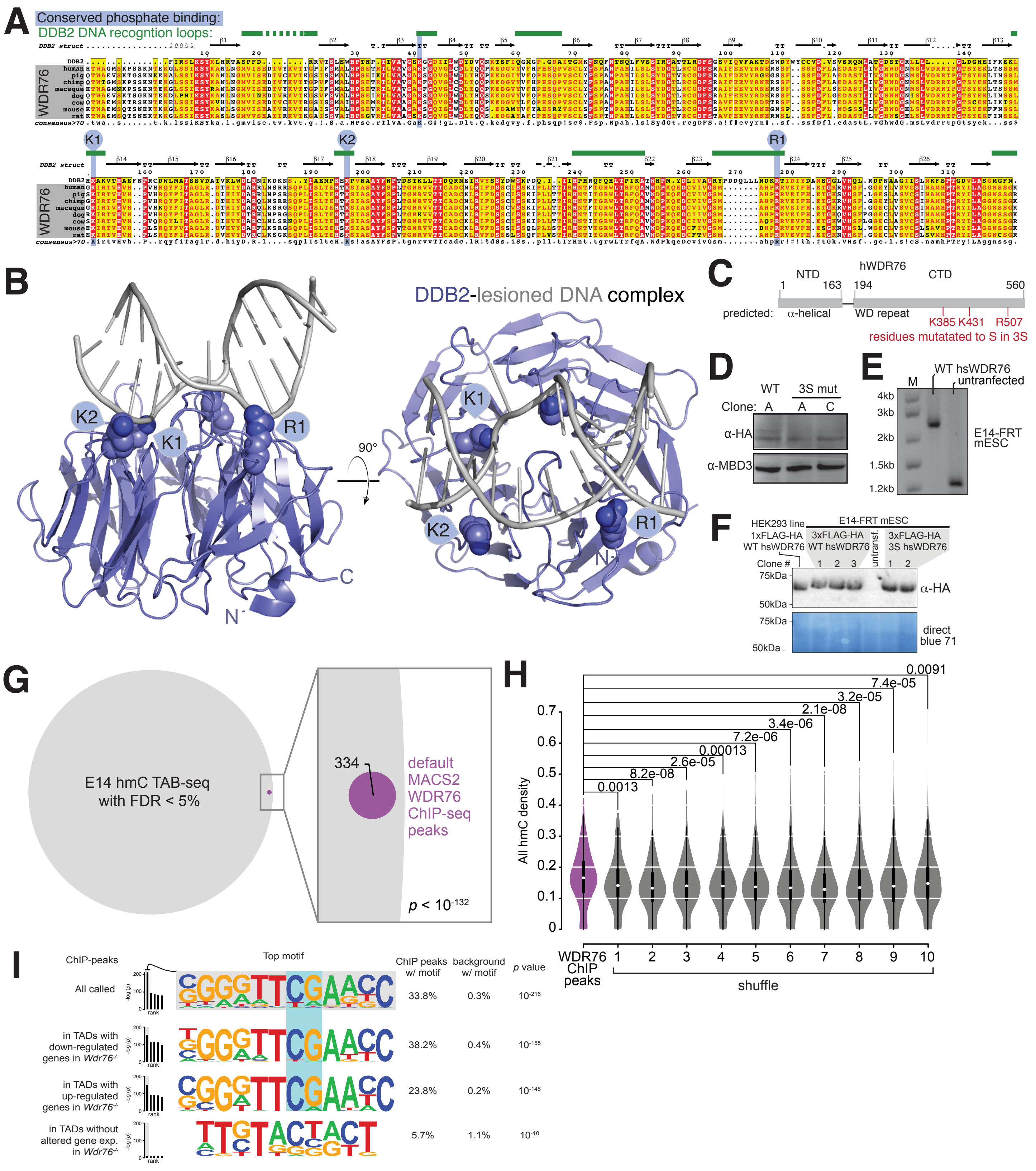
A
Conserved phosphate binding:
DDB2 DNA recogntion loops:
DDB2 struct
WDR76
human
pig
chimp
macaque
dog
cow
mouse
rat
consensus>70
K1
K2
R1
B
DDB2-lesioned DNA complex
K2
K1
R1
90°
C
N
C
hWDR76
NTD
CTD
1
163
194
560
predicted:
α-helical
WD repeat
K385 K431
R507
residues mutatated to S in 3S
D
WT
3S mut
Clone: A
A
C
α-HA
α-MBD3
E
M
WT hsWDR76
untranfected
4kb
3kb
2kb
1.5kb
1.2kb
E14-FRT mESC
F
HEK293 line
1xFLAG-HA WT hsWDR76
E14-FRT mESC
3xFLAG-HA WT hsWDR76
untransf.
3xFLAG-HA 3S hsWDR76
Clone #
1
2
3
1
2
75kDa
50kDa
α-HA
direct blue 71
G
E14 hmC TAB-seq with FDR < 5%
334
default MACS2 WDR76 ChIP-seq peaks
p < 10-132
H
All hmC density
0.0091
7.4e-05
3.2e-05
2.1e-08
3.4e-06
7.2e-06
0.00013
2.6e-05
8.2e-08
0.0013
WDR76 ChIP peaks
1
2
3
4
5
6
7
8
9
10
shuffle
I
ChIP-peaks
Top motif
ChIP peaks w/ motif
background w/ motif
p value
All called
33.8%
0.3%
10-216
in TADs with down-regulated genes in Wdr76-/-
38.2%
0.4%
10-155
in TADs with up-regulated genes in Wdr76-/-
23.8%
0.2%
10-148
in TADs without altered gene exp. in Wdr76-/-
5.7%
1.1%
10-10
-log (p)
rank

### Figure S4

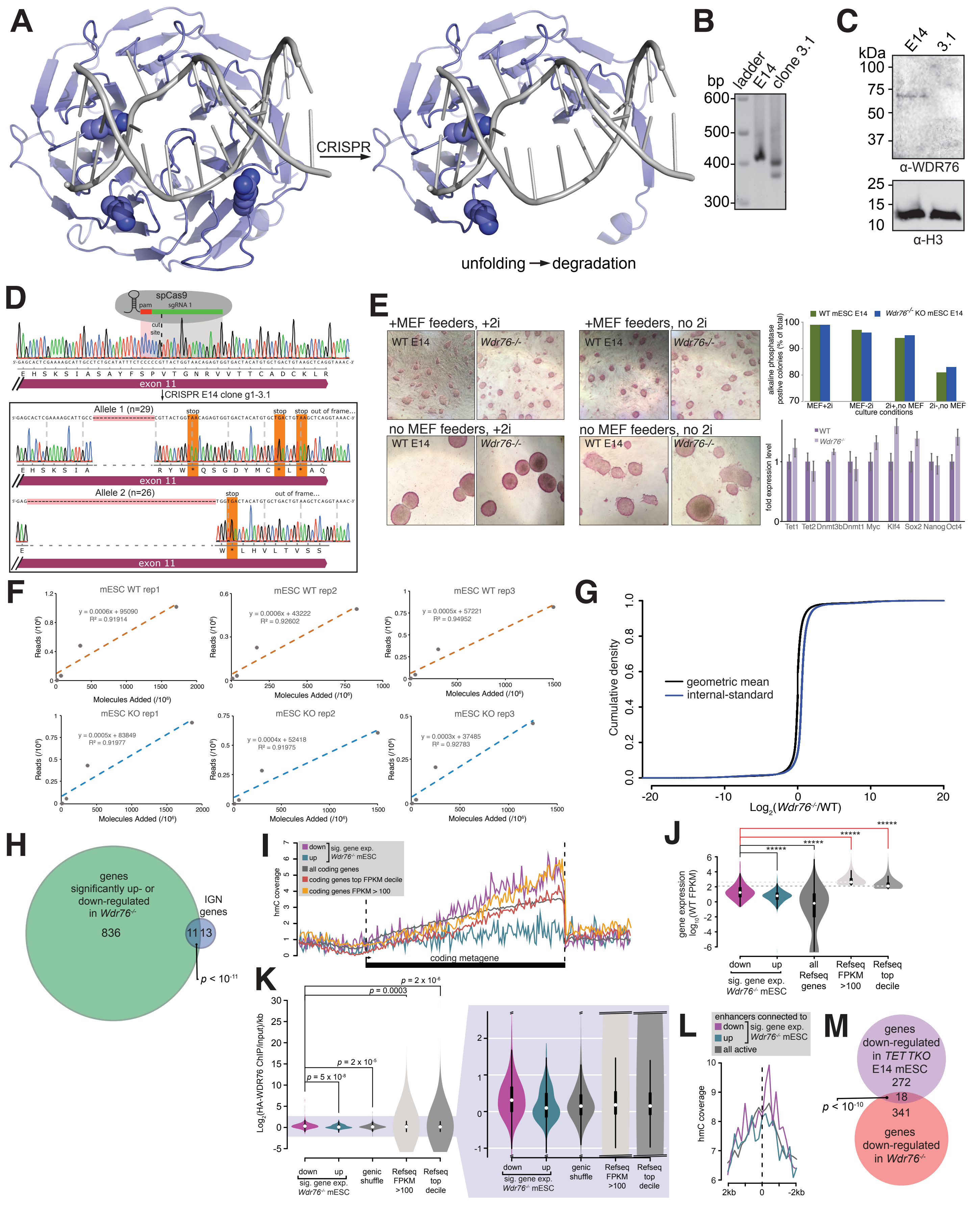
A
CRISPR
unfolding → degradation
B
bp ladder E14 clone 3.1
600
500
400
300
C
kDa E14 3.1
α-WDR76
α-H3
D
spCas9
pam
sgRNA 1
cut site
exon 11
CRISPR E14 clone g1-3.1
Allele 1 (n=29)
Allele 2 (n=26)
E
+MEF feeders, +2i
+MEF feeders, no 2i
no MEF feeders, +2i
no MEF feeders, no 2i
WT E14
Wdr76-/-
WT mESC E14
Wdr76-/- KO mESC E14
alkaline phosphatase postive colonies (% of total)
culture conditions
fold expression level
Tet1 Tet2 Dnmt3b Dnmt1 Myc Klf4 Sox2 Nanog Oct4
F
mESC WT rep1
mESC WT rep2
mESC WT rep3
mESC KO rep1
mESC KO rep2
mESC KO rep3
Reads (/10⁶)
Molecules Added (/10⁶)
G
geometric mean
internal-standard
Cumulative density
Log2(Wdr76-/-/WT)
H
genes significantly up- or down-regulated in Wdr76-/-
836
IGN genes
11 13
p < 10-11
I
hmC coverage
coding metagene
all coding genes
coding genes top FPKM decile
coding genes FPKM > 100
J
gene expression log10(WT FPKM)
down up
sig. gene exp. Wdr76-/- mESC
all Refseq genes
Refseq FPKM >100
Refseq top decile
K
Log2(HA-WDR76 ChIP/input)/kb
p = 2 x 10-6
p = 0.0003
p = 2 x 10-5
p = 5 x 10-8
genic shuffle
L
enhancers connected to
all active
hmC coverage
M
genes down-regulated in TET TKO E14 mESC
272
18
341
p < 10-10
genes down-regulated in Wdr76-/-

### Figure S5

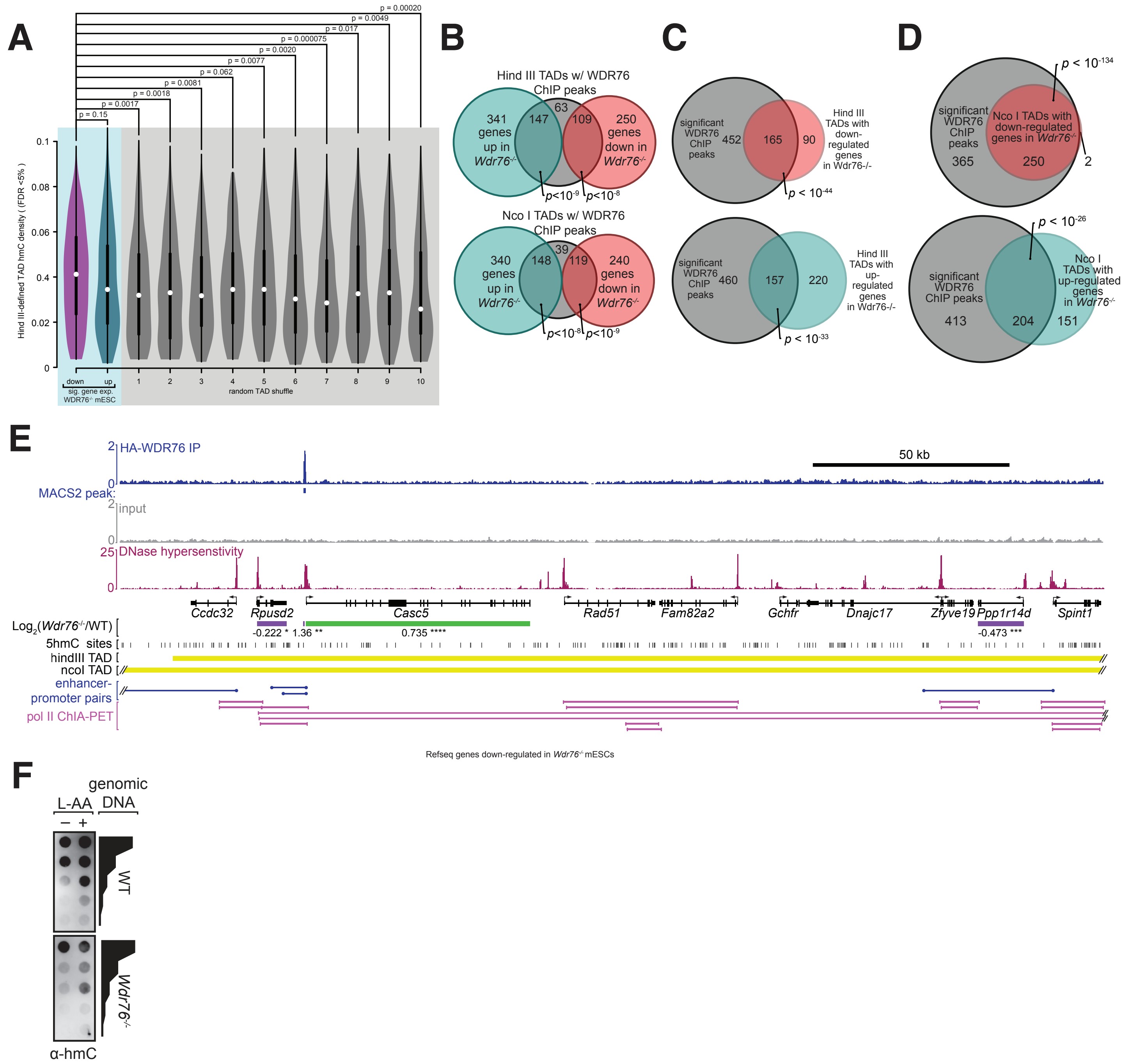
A
p = 0.15
p = 0.0017
p = 0.0018
p = 0.0081
p = 0.062
p = 0.0077
p = 0.0020
p = 0.000075
p = 0.017
p = 0.0049
p = 0.00020
Hind III-defined TAD hmC density ( FDR <5% )
down
up
sig. gene exp. WDR76-/- mESC
random TAD shuffle
B
Hind III TADs w/ WDR76 ChIP peaks
341 genes up in Wdr76-/-
147
63
109
250 genes down in Wdr76-/-
p<10-9
p<10-8
Nco I TADs w/ WDR76 ChIP peaks
340 genes up in Wdr76-/-
148
39
119
240 genes down in Wdr76-/-
p<10-8
p<10-9
C
significant WDR76 ChIP peaks
452
165
90
Hind III TADs with down-regulated genes in Wdr76-/-
p < 10-44
significant WDR76 ChIP peaks
460
157
220
Hind III TADs with up-regulated genes in Wdr76-/-
p < 10-33
D
p < 10-134
significant WDR76 ChIP peaks
Nco I TADs with down-regulated genes in Wdr76-/-
365
250
2
p < 10-26
significant WDR76 ChIP peaks
Nco I TADs with up-regulated genes in Wdr76-/-
413
204
151
E
HA-WDR76 IP
50 kb
MACS2 peak:
input
DNase hypersenstivity
Ccdc32
Rpusd2
Casc5
Rad51
Fam82a2
Gchfr
Dnajc17
Zfyve19
Ppp1r14d
Spint1
Log2(Wdr76-/-/WT)
-0.222 *
1.36 **
0.735 ****
-0.473 ***
5hmC sites
hindIII TAD
ncoI TAD
enhancer-promoter pairs
pol II ChIA-PET
Refseq genes down-regulated in Wdr76-/- mESCs
F
L-AA
genomic DNA
WT
Wdr76-/-
α-hmC

### Figure S6

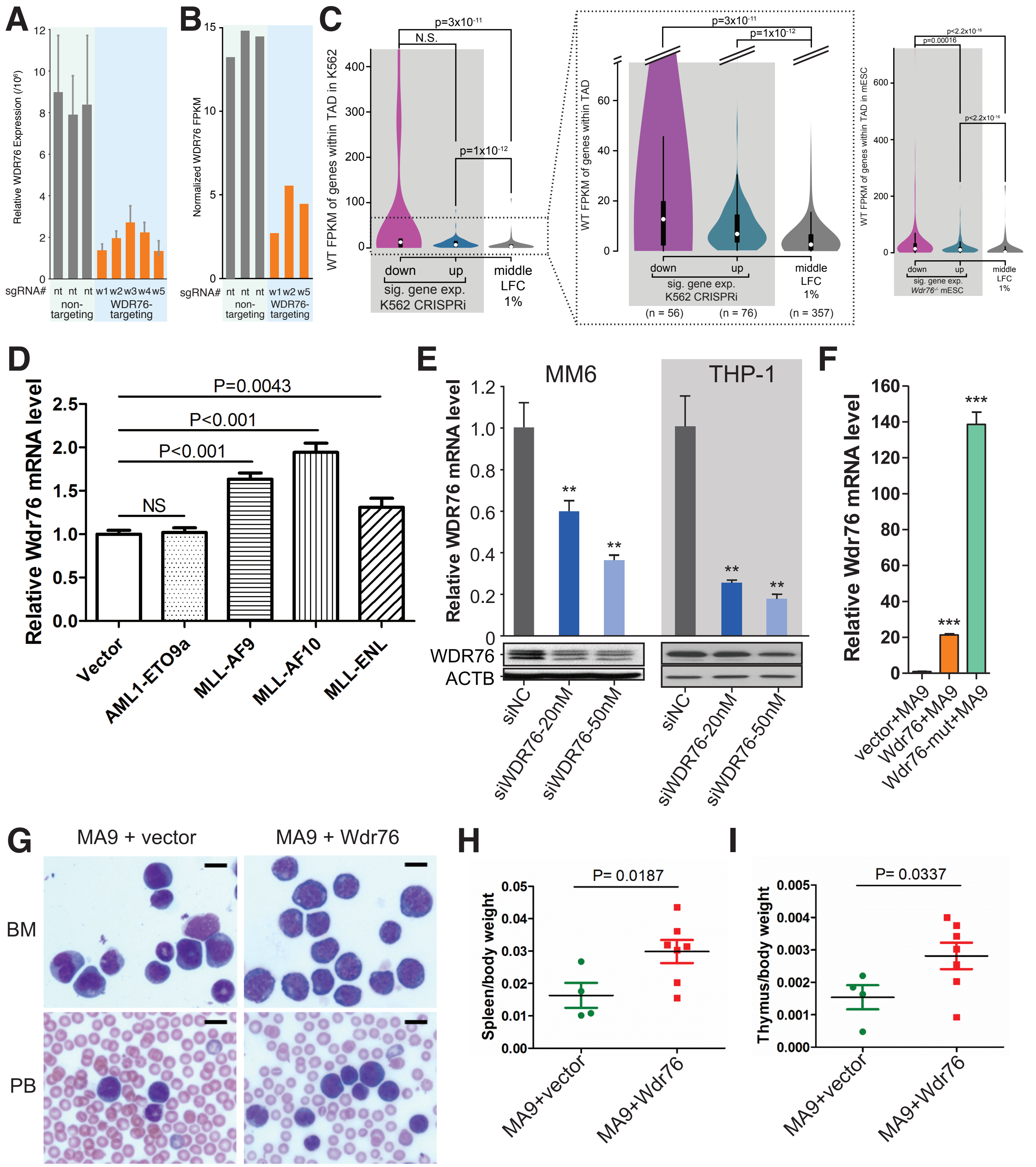
A
B
C
D
E
F
G
H
I
MM6
THP-1
WDR76
ACTB
MA9 + vector
MA9 + Wdr76
BM
PB
P=0.0043
P<0.001
P<0.001
NS
P= 0.0187
P= 0.0337
Relative Wdr76 mRNA level
Relative WDR76 mRNA level
Spleen/body weight
Thymus/body weight
Vector
AML1-ETO9a
MLL-AF9
MLL-AF10
MLL-ENL
siNC
siWDR76-20nM
siWDR76-50nM
vector+MA9
Wdr76+MA9
Wdr76-mut+MA9
MA9+vector
MA9+Wdr76
sig. gene exp.
K562 CRISPRi
down
up
middle
LFC
1%

### Figure S7

#### Extended Data Figure Captions

**Figure S1. Quantitative binding measurements with purified proteins expressed in *E. coli* do not display preferential [ox]mC binding reported from mass spectrometry studies, Related to Figure 1**

(A) Sequence of 16-mer oligonucleotides used for *in vitro* DNA binding studies composed of the strongest E14 mESC hmC-bearing motif (Yu et al., 2012). Complementary strands with the desired modifications were 5' [$^{32}$P] end-labeled with PNK, column purified, and annealed. A scrambled sequence duplex ("nonCpG") with the same base composition without CpG dinucleotides was also employed.

(B) The final stage of homogeneity assessed by SDS-PAGE for four of the strongest hmC-binding candidates (Spruijt et al., 2013) that can be expressed in *E. coli.* after extensive chromatographic purification: THY28, C3ORF37/HMCES (both the "NTD", N-terminal SRAP domain retaining the GST tag, and full-length protein), THAP11/Ronin, and ZHX1 were subjected to quantitative binding studies.

(C) Filter binding experiments with THY28 concentration range that spanned into mid-micromolar regime but did not detect appreciable binding to any of the labeled duplexes examined. "bound" and "free" column panels correspond to signal observed for a given technical replicate on the nitrocellulose membrane and underlying Zeta-Probe membrane, respectively. Dilution series of THY28 without tag, Superdex 75-purified for C/C: 140μM, 56μM, 1.7x fold dilution down to 0.56 μM; for mC/mC, hmC/hmC, carC/carC, fC/fC, nonCpG: 140μM, 55μM, 1.7x fold dilution down to 0.55μM.

(D) Plot of average fraction C, mC, or hmC duplexes bound by C3ORF37-NTD or C3ORF37 full-length, obtained from n=4 and n=2 technical replicates, respectively. Rather than the suggested specific hmC recognition (Spruijt et al., 2013), no appreciable modification-specific DNA binding is observed for the two different NTD experiments, whereas the full-length protein displayed weak binding to symmetric mC and hmC – but not C – at concentrations at least approximately 3 μM and higher. As no clear specificity difference between mC and hmC is revealed by these data, the "HMCES" designation (5-*h*ydroxy*m*ethyl*c*ytosine-binding, *e*mbryonic *s*tem cell-specific) that has recently appeared in HGNC and NCBI databases does not appear to be warranted.

(E) Raw filter binding assay data from C3ORF37 constructs used to generate the plot in panel D. Dilution series for C3ORF37-NTD: 13μM, 10μM, 7.7μM, 1.6x fold dilution down to 0.15 μM; for C3ORF37 full length: 6.3μM, 3.0μM, 1.2μM, 1.5x fold dilution down to 0.045 μM.

(F) Raw data from filter binding assays with THAP11 used to generate the plot in Figure 1B. Dilution series for THAP11: 7.5μM, 2.7μM, 1.1μM, 1.6x fold dilution down to 6.1 nM. THAP11 exhibits equivalent affinity for hmC, C, and nonCpG within experimental error. Importantly, these observations indicate no preference for hmC over C without modification in two distinct sequence contexts, which is far more abundant in the genome.

(G) Purified ZHX1 was subjected to an EMSA using symmetric unmodified C, mC, hmC and fC DNA. ZHX1 protein was titrated in an effort to observe concentration-dependent shift with minimal aggregation in the wells. No clear DNA binding activity of this sequence is observed in the absence of cold competitor, or in the presence of 10-fold molar excess C or mC cold competitor.

(H) EMSA of 6x-His-THY28 with the indicated labeled duplexes in the presence and absence of 5-fold molar excess mC/mC cold competitor. No clear binding is detected by this assay, although weak smears are observed for symmetric hmC and hmC/CpG as a function of protein concentration.

(I) Left panel, Final purity of constructs used by SDS-PAGE. Left middle panel EMSA with 13 μM SALL1 ZFC3 and 1 nM radiolabeled duplex indicated with 50 nM of the indicated specific cold competitor and 50 ng/μL poly (dIdC). Right middle panel, EMSA with 3 μM SALL1 ZFC4 reveals nearly complete shift of all duplexes under identical conditions. Right panel, 3 μM SALL4 ZFC1 does not display much apparent selectivity under these conditions. Although the individual ZFCs display similar properties to those reported (Xiong et al., 2016), any hmC-specificity of SALL1 ZFC3 is countermanded by higher affinity and 5-position non-specific binding of the other modules in the context of the nearly full-length protein.

(J) A construct spanning SALL1 ZFC1-4, was expressed, purified and subjected to EMSA analysis (right panel). Absent cold competitor (middle), most of this protein (5.5 μM) remains in the wells, but that which enters the gel does not display much selectivity. With 6.2 μM and cold mC and C competitors (10-fold and 50-fold molar equivalents with respect to the labeled duplex), even well-retention is ablated.

**Figure S2. Fractionation biochemistry: chromatographic separation and EMSA fractions to isolate that major apparent [ox]mC-binding DNA-binding activity from pig brain, Related to Figure 2**

(A) The chromatography sequence used in the preparation of major [ox]mC activities from clarified porcine cerebrum soluble nuclear extract. Throughout, the blue indicates $UV_{280}$ absorbance chromatogram, the brown trace indicates salt gradient at the mixer, and gray lines with alternating gray shading indicate relevant fractions collected, and red gradient bar indicates activity peak; in several cases flow through and wash portions of the chromatogram are cropped so that the relevant individual fractions can be seen more clearly. Pooled active [ox]mC-binding fractions from the POROS Heparin column indicated (panels C and D) were subjected to 6 mL Resource S chromatography (panel E), then 1 mL Mono S. Fractions from Mono S chromatography containing predominantly the high mobility activity (also asymmetric substrate discrimination remains apparent, panel F) were pooled together for further purification. The fourth column purification step, a chromatography with a 3.3mL DEAE-5PW, concentrates the activity; the portion of the chromatogram with the active fractions (panel G) is shown. The final bulk chromatographic step, either a Superdex 200 or 75 (S200 or S75), affords size-based isolation of the [ox]-mC specific activity, and can separate residual lower EMSA mobility activity from the more [ox]mC-specific higher mobility activity (panels H, I).

(B) A strong carC-specific activity can be separated from the major mobility activity, but occurs inconsistently from prep-to-prep. A unique protein band specific to carC is observed in the oligonucleotide pull down assay. Together, these observations suggest that multiple specific activities for the different [ox]mC states, including carC, exist within brains and partially co-purify through part of the preparation.

(C) EMSA analysis of the peak of [ox]mC-binding activity POROS Heparin fractions indicated with radiolabeled symmetric mC, hmC and fC duplexes in the presence of 10-molar equivalents of cold C or mC.

(D) More detailed specificity screen of POROS Heparin fraction 14, indicating interesting symmetry/asymmetry preferences emerging at this early stage in the preparation.

(E) Alternating fractions spanning the hmC-specific activity peak from Resource S column screened by EMSA with hmC radiolabeled and 10 equivalents of cold mC DNA.

(F) EMSA of the indicated Mono S fractions and flow-through from column loading (FT), examining symmetric hmC-binding in the presence of indicated cold competitor (top panel). In a lower percentage polyacrylamide gel (middle panel), the higher mobility complex in both fractions 18 and 22 exhibits strong binding for symmetric hmC, fC, and to a lesser extent, mC, in the presence of 15-fold molar excess of mC cold competitor DNA. When symmetric hmC is provided as a cold competitor in 15-fold excess, it acts as a specific competitor to both the mC and fC shifts, yet the radiolabeled hmC-binding persists. We note that at later purification stages this concentration of cold hmC is an effective competitor (panel I). Fraction 22 displays an additional lower mobility activity that is less discriminating between the symmetric mC, hmC, and fC DNA, but shows similar properties with asymmetric presentations as the higher mobility activity exemplified by fraction 18. In the bottom panel, the full gel of the portion displayed in Figure 2C is presented, showing more effective competition of fC, as compared to mC and C, which do not afford substantial competition for hmC and fC binding. This is consistent with the idea that the dominant activity present preferentially binds fC over hmC, although we note that there is residual binding that seems insensitive to the concentration of cold fC in this range, perhaps indicating a separate hmC-specific binding activity present.

(G) EMSA shows that both the high and low mobility activities co-elute in DEAE fraction 6, with the latter tailing into later fractions (and pooled separately for sizing).

(H) In this preparation, the lower and higher mobility activities in EMSA were largely separated by S200 size exclusion.

(I) Screen of [ox]mC cold competitors when the higher mobility activity is predominantly isolated from the lower mobility activity and cold hmC acts as an effective competitor. This preparation is one of the two that exhibited a clear carC-specific activity that is stronger than the activity for hmC and fC, whereas the fC binding is unusually weak. This carC-specific property was highly variable from preparation-to-preparation, rendering it challenging to study, but the sensitivity to [ox]mC cold competitor still suggests a ranked preference for these modification states over mC.

(J) Active dialyzed POROS Heparin fractions were subjected to EMSA with a titration of salt provided during binding equilibration prior to electrophoresis. Despite severe gel running artifacts due to the varying salt, the shifted complex is remarkably insensitive to high concentrations (up to 1.5M).

(K) The [ox]mC activity (for fC/C) of the final purified lower and higher mobility activities from S200 purification is insensitive to various chemical additives, including metals, metal chelators (Zn-chelator 1,10-phenanthroline is labeled 1,10 PA), reducing agents, oxidizing agents, and RNAse A treatment, yet is susceptible to SDS.

**Figure S3. Identification of WDR76 as an hmC-specific binding protein and biochemical and cell-based characterization of these properties, Related to Figure 3**

(A) Silver stained SDS-PAGE showing the protein composition of fractions spanning the peak [ox]mC activity from final Superdex 200.

(B) A representative MS2 peptide spectrum for a WDR76 peptide found predominantly in [ox]mC-specific pull downs and active size exclusion fractions (Mascot ion = 44.1 ($p < 4x10^{-5}$); Mascot identity score = 51.5; n =15, independent observations in 15 samples at 90% peptide confidence in Peptide Prophet (Keller et al., 2002). The fragmentation of the peptide is shown in the inset with the observed b and y ions noted in blue and red, respectively. Ions observed with additional protons, loss of ammonia (-17) from R, K, or Q, or loss of water (-18) from S, T, or E, are labeled green.

(C) M2-agararose followed by POROS heparin purification of recombinant WDR76 produced in insect cells.

(D) Full gel of cropped Figure 3D displaying the slight specificity improvement afforded by additional DEAE-5PW chromatography beyond the material depicted in panel C.

(E) Titration of radiolabeled C, mC and hmC (10, 20, 50, 100, 200, 400 nM) against insect cell purified human WDR76 EMSA in the presence of 40 $\mu$M of cold competitor indicated, affords its relative affinity for each modification state. Despite radiolabeling equimolar DNA in side-by-side reactions, the hmC duplex incorporated markedly less radioactivity per mole than the mC and C duplexes, and thus the slightly greater apparent shift observed here at a given protein concentration point for hmC, is an underestimate of the hmC-specificity. The major shift has an electrophoretic mobility of approximately ~300 bp, similar to that of the natively purified activity. A higher mobility shift is also observed that is both protein and oligonucleotide identity dependent.

(F) Putative domain structure of human WDR76 based on secondary structure prediction, homology to other WD repeat proteins and fragment analysis (jpred TNG).

(G) Circular dichroism measurements of the N-terminal domain (NTD) shown demonstrating its alpha helical nature (bi-modal negative ellipticity with local minima at 220 and 210 nm).

(H) Filter binding of the WDR76 NTD indicates that it does bind DNA, but it displays only modest discrimination for hmC opposite unmodified C. Inset: hWDR76 NTD (1-151) expressed in *E. coli* as an N-terminal His-tag fusion protein and purified by IMAC, heparin, and size exclusion chromatography, Coomassie-stained SDS-PAGE of the peak fraction presented.

(I) A C-terminal domain (CTD) comprising 7-8 bladed beta-propeller WD repeats (201-568), was expressed in insect cells as a N-terminal FLAG fusion protein and purified by FLAG M2, POROS Heparin, then S200 chromatography (inset). This preparation is very weakly active in a gel shift assay-- in the absence of cold competitor this is some specificity for hmC/hmC, weaker binding for symmetric mC and fC, and no apparent binding for unmodified duplexes. The presence of cold competitor smears the shift, obscuring assessment of specificity (data not shown).

(J) EMSA analysis of human FLAG-HA WT WDR76 expressed ectopically in HEK293 cells, and purified by M2-agarose and SP-fast flow, then step eluted from a 3.3 mL DEAE-5PW column.  With 50 nM radiolabeled duplexes indicated, and molar-fold excess cold competitors specified, this material displays similar properties to the insect-cell purified protein (hmC-specific, with some affinity for mC, and no detectable binding to fC), with one distinction: the hmC/mC is not appreciably bound here.

**Figure S4. Design of a WDR76 mutant deficient in hmC-DNA binding and production of corresponding ectopically tagged cell lines, Related to Figure 4**

(A) Comparison of the structurally characterized zebrafish DDB2 sequence by alignment with WDR76 sequences from several mammals. DDB2 is a close sequence homolog of WDR76 within their WD repeat domains (55% homology, 25% identity). Secondary structure elements and loops on the top surface involved in DDB2-DNA interactions are indicated with green bars (Scrima et al., 2008). We targeted conserved lysine and arginine residues within these loops involved in phosphate backbone to erode WDR76's DNA-binding capacity.

(B) The positions of all conserved basic residue-phosphate contacts depicted in van Der Waals spheres on the structure of DDB2 with abasic-site DNA (PDB:3EI2) (Scrima et al., 2008), sites indicated K1, K2, and R1 were chosen for mutation to serine. All available data suggest comparable folding of the mutant relative to WT protein: indistinguishable protein expression and solubility in HEK293-FRT cell lines, absence of abundant or distinct protein degradation bands by western blotting, and some residual DNA binding activity under conditions with less aggressive washing.

(C) Putative domain structure of human WDR76 with sites targeted for mutation in the 3S mutant indicated.

(D) HA-blot of WT and 3S WDR76 stable HEK293 cell lines with equivalent expression as compared to MBD3 loading control.

(E) representative genomic PCR confirmation of integration of FLAG-HA-hWDR76 constructs in mESC E14 FRT cell lines; the insertion of the tagged expression construct into the FRT site (EF1-FLAG-HA-hWDR76 = 2670 bp vs. intact untransfected line = 1290 bp).

(F), Western blotting of ectopically expressed 3xFLAG-HA hsWDR76 (WT or 3S mutant) mESC clones following line selection with 12CA5 ($\alpha$-HA) antibody, as compared to the untransfected parental line and an HEK293 1xFLAG-HA line, direct blue staining of the blot is shown for loading levels.

(G) Venn diagram as in Figure 4A, but using the set of peaks called by MAC2 with default settings rather than broad peaks recapitulates massive overrepresentation of WDR76 peaks at high confidence hmC sites (Yu et al., 2012).

(H) Violin plot analysis of hmC density (%hmC/bp) (Yu et al., 2012), for default MACS2 peaks with 500 bp cushion added to each side, compared to 10 random shuffled chromosomal position sets with the same spans using Bedtools shuffle (Quinlan and Hall, 2010), *p* values are computed using unpaired Wilcoxon-Mann-Whitney test.

(I) Independent motif enrichment analysis for different subsets of ChIP called peak set (All called peaks, peaks in TADs with up or down-regulated genes in the *Wdr76* knockout, versus peaks in all other TADs). The top motif in each analysis (depicted) as compared to other the top 5 ranked motifs is presented as a bar graph of the -log10 (*p*-value), and the top motif within the indicated set and background sets is depicted; *p*-values computed via Homer motif enrichment binomial test versus background set. Very similar top motifs are highly enriched in functionally important sites, but not detected in all other ChIP peaks outside if TADs with down- and up-regulated genes in the WDR76 knockout.

**Figure S5. Generation of *Wdr76* knockout mESCs, and functional genomic analysis of its role in gene expression, Related to Figure 5**

(A) Schematic of the anticipated protein-level results of exon 11 nonsense mutation introduced by CRISPR. Given the $\beta$-propeller structure requires 6-8 intact blades composed of WD40 repeats to stably fold (Fülöp and Jones, 1999), these mutations should preclude the folding of a module for which three point mutations in this domain are sufficient to disrupt the DNA-binding capacity of WDR76.

(B) Generation of *Wdr76*$^{-/-}$ mESC cell line with CRISPR/Cas9 editing. PCR screen of an amplicon centered on the CRISPR target site demonstrates distinct mobilities of each allele of clone g1-3.1 relative to the parental E14 line.

(C) Western blot demonstrating loss of Wdr76 protein in the g1-3.1 clone with an H3 loading control.

(D) Location in *Wdr76* exon 11 of the target site for the sgRNA1 overlaid on representative sequencing of TA-cloned amplicon from the g1-3.1 clonal cell line demonstrating clonal purity of bi-allelic deletions (n = 29, 26 TA clones of each allele with no other alleles detected) causing frame-shift mutations that result in missense mutations that delete the final three predicted WD40 repeats.

(E) WDR76 knockout does not result in changed colony morphology or percent of AP positive colonies under the same growth conditions (+2i or MEF feeders), nor altered expression of key pluripotency genes and DNA methylation enzymes (n=3 independent replicates for each condition).

(F) Linear regression fits of spiked-in RNA standard reads (Werner et al., 2017) as a function of molecules of each standard added

(G) Cumulative density plot of the $\log_2$-fold change of mRNA in the knockout versus the WT E14 line, comparing internal standard to geometric mean normalization. As there is little difference, we used geometric mean normalization for all comparative analyses.

(H) Overlap of significantly altered genes (4 down, 7 up) with the 24 imprinted gene network of somatic stem cell/early development genes subject to mono-allelic methylation mediated gene regulation (Berg et al., 2011). *p*-value for overrepresentation over the null hypothesis of chance overlap was calculated with two-tailed Fisher Exact test phyper in R against the background of all Refseq genes.

(I) The E14 hmC coverage (Yu et al., 2012) contoured over metagenes for the indicated gene sets.

(J) The distribution of WT expression levels expressed as $\log_{10}$ (FPKM) for genes in the sets described in panel A. ******$p < 2\text{x}10^{-16}$*, computed by unpaired Mann-Whitney-Wilcoxon two-tailed test, for the left two comparisons (black line), the down-regulated gene set FPKM distribution is greater than chance expectation, and for the right two comparisons (red line), the *p* value represents probability that down-regulated gene set distribution is less than the indicated comparison group by chance.

(K) WDR76 ChIP density per gene expressed as $\log_2$-fold change of ChIP density [(WDR76 IP/input)/kb] for indicated gene sets as in Figure 5C. All *p*-values are computed by Mann-Whitney-Wilcoxon two-tailed test for the overrepresentation of WDR76 ChIP signal over chance in the down-regulated genes in the knockout relative to indicated partner.

(L) Contouring high confidence hmC coverage over all active center-point aligned enhancers and those that are computationally connected to target genes (Shen et al., 2012) whose expression is significantly up- or down-regulated in the *Wdr76* knockout.

(M) Venn diagram comparison of E14 genes down-regulated in the TET1/2/3 triple knockout (Lu et al., 2014) versus Wdr76 knockout; *p*-value for overrepresentation over the null hypothesis of chance overlap was calculated with two-tailed Fisher Exact test phyper in R.

**Figure S6. Combined analysis of WDR76 ChIP, RNA-seq from *Wdr76$^{-/-}$* and hmC density at the level of TADs, Related to Figure 6**

(A) Violin plot of hmC density within TADs that contain the indicated the significantly altered gene sets in the *Wdr76$^{-/-}$* cell line, as compared to 10 representative random genic shuffles of the “down” set’s coordinates using the FDR < 0.05 set of sites (Yu et al., 2012); *p*-values computed via unpaired Mann-Whitney-Wilcoxon test for hmC overrepresentation in the down-regulated gene group compared to each other indicated.

(B) The overlap of genes that are significantly down or up-regulated in the *Wdr76$^{-/-}$* mESCs with topologically associating domains (TADs) bearing significant WDR76 ChIP-seq peaks from two different E14 mESC TAD datasets (Hind III and Nco I) (Dixon et al., 2012).

(C) Same analysis presented in Figure 6D, but without the with 10% length cushion.

(D) Same analysis presented in Figure 6D, but using the Nco I digest dataset defined TADs rather than the consensus Hind III TADs (Dixon et al., 2012). WDR76 broad peaks are intersected with TADs that bear up- or down- regulated genes in the *Wdr76$^{-/-}$* mESCs and *p*-values computed by two-tailed Fisher Exact test in Bedtools with 10% length cushion.

(E) Representative locus view of data used in the genome-wide analyses: WDR76 ChIP and input coverage presented at normalized reads per million, significantly altered genes from RNA-seq are indicated in (purple and green), presented alongside the high-confidence E14 hmC site data (Yu et al., 2012), with DNase I hypersensitivity and RNA pol II ChIA-PET from mESCs (Kieffer-Kwon et al., 2013), predicted promoter-enhancer pairs in mESCs (Shen et al., 2012) and E14 mESC TADs (Dixon et al., 2012).

(F) Inset is an $\alpha$-hmC dot blot of 2-fold serial dilution starting with 50 ng genomic DNA after 24 hours of L-AA treatment.

**Figure S7. The function of WDR76 in mouse and human leukemia models, Related to Figure 7**

(A) RT-qPCR of WDR76 expression relative to 18*S* RNA in K562 dCAS9-KRAB cell lines transfected with five different guide RNAs that target the promoter of WDR76 as compared to control non-targeting guide RNAs

(B) Geometric mean-normalized FPKM of WDR76 from RNA-seq of non-targeting sgRNA (nt, n=3) versus WDR76 directed depletions (w1, w2, w5) demonstrates appreciable knock-down.

(C) CRISPRi depletion of WDR76 in human myeloid leukemia cells displays mixed expression effects traceable to alterations in expression of highly transcriptionally active topologically associated domains (TADs). The average mRNA expression of TADs that contain genes significantly down- or up-regulated, versus unchanged expression (middle percentile) when WDR76 is depleted by CRISPRi in K562 (Gilbert et al., 2014) cells using three different guide RNAs. Violin plots with zoomed inset shows the bulk of the distributions distributions more clearly absent the outliers (*p*-values computed via Mann-Whitney-Wilcoxon test). Although no hmC dataset exists for this line, this expression analysis indicates genes within highly expressed TADs are preferentially modulated by WDR76, akin to what we observe in the mESC system (right panel).

(D) Comparison of Wdr76 mRNA levels in mouse bone marrow progenitor cells transduced with the indicated viruses. Analogous to the human patient samples in Figure 7A, cells transduced with three individual MLL-rearranged fusions, MLL-AF9, MLL-AF10, and MLL-ENL, display significantly elevated *Wdr76* expression as compared to those transduced with empty vector or the AML1-ETO9a oncogene.

(E) Knock-down efficiency at the mRNA and protein levels for the two human MLL-AF9 fusion leukemic cell lines presented in Figure 7B-C.

(F) Relative expression of *Wdr76* in the various MLL-AF9 (MA9) virally co-transduced bone marrow progenitor cells presented in Figure 7E; its expression level in the vector+ MLL-AF9 group was set as 1.

(G) Histopathology of bone marrow (BM) and peripheral blood (PB) of bone marrow transplant recipient mice (MA9 is the MLL-AF9 fusion).

(H,I) Weight of spleen and thymus relative to body weight at the time of sacrifice of recipient mice in Figure 7F.

### METHODS

#### CONTACT FOR REAGENT AND RESOURCE SHARING

#### METHOD DETAILS

##### Preparation of oligonucleotides

The duplex sequence composition is designed to present a central **CpG** dinucleotide based on the highest scoring motif in base-resolution TAB-seq from mESCs (Yu et al., 2012) step ('top': 5'-GCGGCTG**CG**TGGGTCC-3'; 'bottom': 5'-GGACCCA**CG**CAGCCGC-3', underlined C positions are subject to 5-position modification). Biotinylated oligonucleotides were synthesized with the same 'bottom' sequence with a 3' $T_8$-TEG extension, yielding a 24 nucleotide 3' biotinylated oligonucleotide. These were synthesized on a 3' biotin CPG (Glen Research) using the synthesis and deprotection conditions dictated by the modified cytosine incorporated. This oligonucleotide was annealed with the 'top' sequence to form 3' biotinylated duplexes. Finally, a scramble oligonucleotide duplex lacking CpG content was designed to test the contribution of the CpG dinucleotide step to specific binding ('nonCpG top': 5'- GGCCTGGCCTGGGTCC -3'; 'nonCpG bottom': 5'- GGACCCAGGCCAGGCC).

All ox-mC modified oligonucleotides were synthesized on an Expedite 8909 DNA synthesizer (Applied Biosystems) using conventional solid phase 2-cyanoethyl (CE) phosphoramidite synthesis reagents (Glen Research). The synthesis, deprotection, and purification were according to the manufacturer instructions, with some modifications. Nucleoside phosphoramidites with Ultramild protecting groups (Pac-dA-CE, Ac-dC-CE and iPr-Pac-dG-CE) were used to synthesize the oligonucleotides containing C, mC, hmC and fC. Regular phosphoramidites (N-benzoyl-dA-CE, N-benzoyl-dC-CE and N-isobutyryl-dG-CE) were used to synthesize oligonucleotides containing carC, in order to accommodate its deprotection conditions without side reactions.

Modified cytosines were included at the underlined positions in the sequences below using 5-methyl-dC-CE, 5-hydroxymethyl-dC-CE (5-hydroxymethyl-dC II CE (carbamoyl protecting group spanning exocyclic amine and 5-hydroxyl position), 5-formyl-dC-CE (1,2-diacetyloxy-ethyl and N-acetyl protecting group), and 5-carboxy-dC-CE (5-ethylcarboxyl, N-benzoyl protecting groups). Oligonucleotides bearing C, mC, hmC and fC were synthesized on Ac-dC 1000 Å controlled pore glass (CPG) support on a 1 $\mu$mol scale. Oligonucleotides bearing carC were synthesized on Bz-dC-CPG 1000 angstrom support in micromole scale syntheses.

Oligonucleotides bearing C, mC, and in the central CpG dinucleotide were deprotected with fresh aqueous 30% ammonium hydroxide, shaking at 55°C overnight in benchtop ThermoMixer (Eppendorf). The recovered material was then lyophilized and purified from truncated, degraded, or partially deprotected material by preparative 20% denaturing polyacrylamide gel with after heating to 95°C for 5 minutes in denaturing nucleic acid loading buffer (95% formamide, 5mM EDTA pH 8.0, 0.025% bromophenol blue, 0.025% xylene cyanol). The product band was visualized by UV shadowing on a fluorescent TLC plate, excised, and extracted with 100 mM triethyl ammonium acetate pH 7 (TEAA) shaking at 30°C overnight in 2 x 25mL extractions. This material was then loaded onto a Waters Sep-Pak C18 cartridge, washed in 100 mM TEAA, then MilliQ water and eluted in 50% aqueous acetonitrile. Oligonucleotides containing hmC were deprotected using 20 mM $K_2CO_3$ in methanol for 4 hours shaking at 800 rpm at 30°C in a ThermoMixer, then purified as described above. Oligonucleotides containing fC were deprotected initially in a 1:1 solution of 40% $CH_3NH_2$ and 30% $NH_4OH$ at 65°C shaking at 800 rpm for 2 hours in a ThermoMixer to reveal the 1,2 diol. Following gel purification, Sep-Pak purified material was then lyophilized and resuspended in cold dd$H_2O$ at 4°C. To oxidize the 1,2 diol to generate the aldehyde of fC, aqueous $NaIO_4$ at 4°C was added to a final concentration of 50 mM, and the mixture incubated for 30 minutes at 4°C rotating end over end. This reaction was then quenched with 10 volume equivalents of 100 mM TEAA pH 7 and loaded onto a second C18 Sep Pak for purification as above. Oligonucleotides containing carC were deprotected and cleaved from the resin using 0.4 M NaOH in a 4:1 methanol:water solvent overnight at room 800 rpm at 30°C in a ThermoMixer. The deprotected material was then purified as described above.

Purified oligonucleotides were then analyzed by MALDI-TOF mass spectrometry to confirm identity and complete deprotection/conversion to the desired final molecule. Oligonucleotide purity was assessed by gel to be > 90% in all cases. The oligonucleotides were lyophilized, and resuspended in 20 mM Tris HCl pH 7.5, 1 mM EDTA at 400 μM final concentration, and stored at -80°C.

Duplex oligonucleotide stocks were annealed by heating equimolar top and bottom strands at 80°C in 200 μM in TE300 (20 mM Tris, pH 7.5, 1 mM EDTA, 300 mM NaCl) for 5 minutes in a heading block, followed by passive slow cooling of the block to room temperature over ~1 hour. Small aliquots were then stored at -80°C. Annealed oligonucleotides were analyzed by Sybr Green melts to assess stability of the duplex in a real time PCR instrument (BioRad CFX96). The calculated melting temperature of these 16 base pair oligonucleotides to be 63.5°C, (IDT Oligo Analyzer web tool) however, the observed melting temperature using Sybr Green is closer to ~75°C at the concentrations of NaCl used in labeling, annealing and EMSA procedures (typically ≥300 mM NaCl). Freshly thawed, annealed oligonucleotides were radiolabeled with T4 PNK (NEB) overnight using 50 pmol of [$\gamma$-$^{32}$P]ATP per 50 pmol of 5' ends (6000 Ci/mmol, Perkin Elmer) in 1x PNK buffer. After addition of 20 mM EDTA, the PNK was heat-denatured for five minutes at 95°C, supplemented with 300 mM NaCl, and the mixture slow cooled to reanneal. The concentrated labeled oligonucletotide stock was then diluted in TE300 for use in gel shift assays. For filter binding assays, additional purification of the annealed duplex from residual [$\gamma$-$^{32}$P] ATP was accomplished with Illustra ProbeQuant G-50 Micro Columns (GE Healthcare) according to manufacturer instructions.

#### Preparation of candidate hmC-specific readers from *E. Coli*

*Expression and purification of THY28*

Human THY28 cDNA (residues 54-221), identified by SILAC as a pan-[ox]mC reader (Spruijt et al., 2013), was cloned into expression vector pMCSG7 from MGC IMAGE 30717693 (NCBI Accession BC093074) to form an N-terminally 6x-His-tagged construct with a TEV protease cleavage site and SNG linker. This construct was designed by reference to prior THY28 expression and structural biology (Song et al., 2005). The resulting plasmid was transformed into *E. coli* BL21(DE3) + pRARE2. Expression cultures were grown at 37°C and induced at $OD_{600}$ = 0.5 with 0.4 mM IPTG for 3 hours at 37°C. Cell pellets were collected via centrifugation (Sorvall RC-3B with H-6000A rotor, 4000 rpm for 20 minutes at 4°C), resuspended in Ni-NTA lysis buffer (600 mM NaCl, 50 mM $Na_xH_{2-x}PO_4$ pH 8.0, 10% glycerol (v/v), 5 mM imidazole, 0.5 mM PMSF, 5 mM β-mercaptoethanol), and stored at –80°C. Freshly thawed cell pellets were lysed with an Avestin EmulsiFlex-C3 homogenizer and lysate was clarified via two sequential centrifugation steps (30,000xg in Sorvall RC-5B with SS-34 rotor, 4°C for 25 minutes). Clarified lysate was incubated with pre-equilibrated $Ni^{2+}$-NTA resin (Qiagen; 0.25 mL final bed volume per 1 L of culture) for 30 minutes in a 2.5x10 cm Econo-column (Bio-Rad), rotating at 4°C. After elimination of flow-through, the resin was washed sequentially with 4 column volumes (CV) each of 1 M NaCl, 600 mM NaCl, and 400 mM NaCl wash buffers (otherwise identical in composition to Ni-NTA lysis buffer). Bound proteins were eluted over multiple fractions in Ni-NTA elution buffer containing 300 mM imidazole pH 7.5 and 400 mM NaCl (otherwise identical in composition to Ni-NTA lysis buffer). Peak fractions enriched for THY28 were pooled, combined with 1/100 mass equivalents of TEV protease, and dialyzed overnight at 4°C against 100 mM NaCl, 50 mM Tris·HCl pH 7.0, 5% glycerol (v/v), 5 mM β-mercaptoethanol. The dialysis output was clarified via centrifugation and diluted approximately 1.5-fold with Buffer TG100 (100 mM NaCl, 20 mM Tris·HCl pH 7.0, 5% glycerol (v/v), 5 mM β-mercaptoethanol). The pH and conductivity of the clarified output was adjusted with 1 M Tris·HCl pH 6.5 and salt free dilution buffer (identical in composition to Buffer TG100 except for the absence of NaCl) to match that of Buffer TG50 (50 mM NaCl, otherwise identical in composition to Buffer TG100). This sample was loaded onto a 6 mL RESOURCE S column (GE Healthcare) pre-equilibrated in Buffer TG50; the column was washed with 2 CV of Buffer TG50 and eluted with a linear gradient over 5 CV to 1 M NaCl. Desired peak eluate fractions were pooled and concentrated with an Amicon Ultra-4 10 kDa NMWL Centrifugal Filter Unit (EMD Millipore) according to manufacturer instructions. This concentrate was filtered with a 0.45 μm Ultrafree-MC HV Centrifugal Filter (EMD Millipore) and loaded onto a Superdex 75 10/300 GL column (GE Healthcare) equilibrated and eluted in 20 mM Na·HEPES pH 7.8, 150 mM NaCl, 5% glycerol (v/v). Peak eluate fractions were pooled, quantified by NanoDrop approximately 320 μM (A260:A280 ratio of 0.58), and subjected to filter binding assays. All column chromatography steps were performed at 4°C.

*Expression and purification of C3ORF37/HMCES*
C3ORF37 is an uncharacterized protein that has been renamed "HMCES" (5-*h*ydroxy*m*ethyl*c*ytosine-binding, *e*mbryonic *s*tem cell-specific) based on mass spectrometry enrichment for [ox]mC (Spruijt et al., 2013). Human C3ORF37 cDNA (full length protein designated "FL", or residues 1-275 designated "NTD" that comprise the predicted SRAP domain (Aravind et al., 2013) was amplified from MGC IMAGE 30398247 (NCBI Accession BC088363) and cloned into the pMCSG20 expression vector to form N-terminally S- and GST-tagged constructs with a TEV protease cleavage site and SNG linker. C3ORF37 FL and NTD constructs were expressed in *E. coli* BL21(DE3) + pRARE2 by induction at $OD_{600}$ = 0.5 with 1 mM IPTG for 12 hours at 16°C. Cell pellets were harvested as above via centrifugation, resuspended in GST lysis buffer (500 mM NaCl, 50 mM Tris·HCl pH 7.5, 10% glycerol (v/v), 0.5 mM PMSF, 5 mM β-mercaptoethanol), and stored at –80°C. Freshly thawed cell pellets were lysed and clarified as described above for THY28. Clarified lysate was loaded onto a 5 mL GSTrap HP column (GE Healthcare) pre-equilibrated in GST lysis buffer; the column was washed with 2 CV of GST lysis buffer and eluted over 3 CV (linear gradient) to GST lysis buffer + 20 mM reduced glutathione. This and all subsequent column chromatography steps were performed at 4°C.

For C3ORF37 FL, the desired GST column peak eluate fractions were pooled and combined with 1/50 mass equivalents of TEV protease, and dialyzed overnight at 4°C against 250 mM NaCl, 50 mM Tris·HCl pH 8.45, 10% glycerol (v/v), 5 mM β-mercaptoethanol. The dialysis output was clarified via centrifugation and adjusted to match the pH and conductivity of Buffer TG100 by sequential addition of the appropriate volumes of 1 M Tris·HCl pH 6.0 and TG0 (identical in composition to Buffer TG100 except for the absence of NaCl). This sample was loaded onto a 5 mL HiTrap Heparin HP column (GE Healthcare) pre-equilibrated in Buffer TG100; the column was washed with 2 CV of Buffer TG100 and eluted over a linear 6 CV gradient to 550 mM NaCl, then to 1 M NaCl over 1 CV. Desired peak eluate fractions were pooled, supplemented with NaCl to achieve a final approximate concentration of 800 mM, and concentrated with an Amicon Ultra-4 10 kDa NMWL Centrifugal Filter Unit (EMD Millipore) according to manufacturer instructions. This concentrate was filtered as described above for THY28 and loaded onto a Superdex 75 10/300 GL column (GE Healthcare) equilibrated and eluted in 20 mM Na·HEPES pH 7.8, 150 mM NaCl. Desired peak eluate fractions were pooled and concentrated via centrifugation, and this concentrate was stored at 4°C for use in filter binding assays (designated “C3ORF37 FL FBAI”). The FL concentration of this sample as measured by NanoDrop was approximately 8.7 μM, with an A260:A280 ratio of 0.62.

The NTD was purified using a similar strategy as that described above for FL, but, as was also the case for the FL preps, despite large expression scales, relatively inefficient and incomplete GST tag cleavage combined with various solubility issues throughout the preps (of greater severity for FL than for NTD) and loss to the membrane of the centrifugal concentrators resulted in significant yield losses between the GST, Heparin, and size exclusion chromatography steps that did not permit the recovery of a sufficiently large amount of protein following size exclusion to perform the desired number of filter binding replicates at final concentrations higher than approximately 4–6 μM. Accordingly, we elected to also purify NTD without cleaving the GST tag . For this prep, the desired GST column peak eluate fractions were pooled and adjusted to match the pH and conductivity of Buffer BTG100 (100 mM NaCl, 20 mM Bis-Tris pH 6.0, 10% glycerol (v/v), 5 mM β-mercaptoethanol) by sequential addition of the appropriate volumes of 770 mM Bis-Tris pH 6.0 and salt free dilution buffer (identical in composition to Buffer BTG100 except for the absence of NaCl). This sample was loaded onto a 5 mL HiTrap Heparin HP column (GE Healthcare) pre-equilibrated in Buffer BTG100; the column was washed with 3 CV of Buffer BTG100 and eluted over a linear 2 CV gradient to 1 M NaCl. Desired peak eluate fractions were pooled and split into two portions: the first (approximately 1/3 of the prep) was directly dialyzed overnight at 4°C against 20 mM Na·HEPES pH 7.8, 150 mM NaCl with one buffer change. The GST-C3ORF37 NTD concentration of as measured by NanoDrop was approximately 14.4 μM A260:A280 ratios of 0.65.

*Expression and purification of THAP11*

THAP11 or Ronin is a protein reported to bind hmC specifically by LFQ MS of material captured from adult mouse brain nuclear extracts (Spruijt et al., 2013) and is also required for pluripotency during early embryogenesis in mouse embryonic stem cells (Dejosez et al., 2008). Residues 2-89 of full length human THAP11 encode a THAP domain responsible for DNA binding activity, which was cloned from MGC IMAGE 4554554 (NCBI Accession BC012182) into the expression vector pMSCG20 to form an N-terminally S- and GST-tagged construct with a TEV protease cleavage site and a SNG linker. The fusion protein was expressed in *E. coli* BL21(DE3) + pRARE2 by induction at $OD_{600}$ = 0.5 with 1 mM IPTG for 16 hours at 16°C. Cell pellets were harvested, lysed and purified over a 5mL GSTrap HP column as described for C3ORF37 above.

The GST column peak eluate fractions were visualized by SDS-PAGE, pooled and combined with 1/50 mass equivalents of TEV protease to remove the GST fusion tag over 16 hours of incubation at 4°C. Complete removal of the tag was confirmed by SDS-PAGE, and as the purified protein showed no nucleic acid contamination at this point (A260:A280 = 0.63), an ion exchange chromatography was omitted. The TEV-digested material was concentrated in a 3kDa MWCO centrifugal concentrator (Vivaspin 2 Polyethersulfone concentrator, Sartorius). The concentrated material was loaded onto a HiLoad 16/600 Superdex 75 size exclusion column equilibrated and developed in 20 mM Na·HEPES pH 7.8, 150 mM NaCl. The peak eluate fractions were inspected by SDS-PAGE, pooled and quantified as described above with a final concentration of 21.1 μM and an A260:A280 ratio of 0.63.

*Expression and purification of ZHX1*

ZHX1 is a protein reported by Spruijt and colleagues (Spruijt et al., 2013) to bind hmC in nuclear extracts from murine neuronal precursor cells. We expressed the largest previously described fragment (55-830) spanning two zinc-fingers and five homeodomains (Bird et al., 2010) cloned from MGC IMAGE 5271344 (NCBI Accession BC040481.1) into pET30a to form an N-terminally 6x-His-tagged construct with a TEV protease cleavage site and SNG linker. The fusion protein was expressed in *E. coli* BL21(DE3) + pRARE2 by induction at $OD_{600}$ = 0.5 with 1 mM IPTG for 4 hours at 37°C. At $OD_{600}$ = 0.25, 50 µM $ZnSO_4$ was added to the cultures to support folding of the zinc fingers during protein expression. Cell pellets were collected via centrifugation (Sorvall RC-3B with H-6000A rotor, 4000 rpm for 20 minutes at 4°C), resuspended in Ni-NTA lysis buffer and stored at –80°C. Freshly thawed cell pellets were lysed with an Avestin EmulsiFlex-C3 homogenizer and lysate was clarified via ultracentrifugation (Beckman Ti70.1 rotor at 37,000 rpm) at 4°C for 40 minutes. Clarified lysate was incubated with pre-equilibrated $Ni^{2+}$-NTA resin (Qiagen; 0.25 mL final bed volume per 1 L of culture) for 60 minutes, rotating at 4°C. After collection of flow-through, the resin was washed sequentially with 4 column volumes (CV) each of 1 M NaCl, 600 mM NaCl, and 200 mM NaCl wash buffers (otherwise identical in composition to Ni-NTA lysis buffer). Bound proteins were eluted over multiple fractions in Ni-NTA elution buffer containing 300 mM imidazole pH 7.5 and 200 mM NaCl (otherwise identical in composition to Ni-NTA lysis buffer). Elution fractions enriched in the desired full length protein were combined and diluted approximately 1.5-fold with Buffer TG100 (100 mM NaCl, 20 mM BisTris pH 6.5, 10% glycerol (v/v), 5 mM β-mercaptoethanol and 50 uM $ZnSO_4$). The pH and conductivity of the fraction pool was adjusted with salt free dilution buffer (identical in composition to Buffer TG100 except for the absence of NaCl) to match that of Buffer TG50 (50 mM NaCl, otherwise identical in composition to Buffer TG100). This sample was loaded onto a 1 mL POROS Heparain S column (GE Healthcare) pre-equilibrated in Buffer TG50; the column was washed with 2 CV of Buffer TG50 and eluted with a linear gradient over 12 CV to 1 M NaCl. Desired peak eluate fractions were pooled and concentrated with an Amicon Ultra-4 30 kDa NMWL Centrifugal Filter Unit (EMD Millipore) according to manufacturer instructions. This concentrate was filtered with a 0.45 µm Ultrafree-MC HV Centrifugal Filter (EMD Millipore) and loaded onto a Superdex 200 10/300 GL column (GE Healthcare) equilibrated and eluted in 20 mM Na·HEPES pH 7.8, 150 mM NaCl, 5% glycerol (v/v). Peak eluate fractions were pooled and stored at 4°C for use in gel shift assays, as described above.

*Expression and purification of SALL1 and SALL4 fragments*
Recently, the SALL1 and SALL4 proteins have been implicated as hmC-binding proteins (Xiong et al., 2016), as one out of several DNA-binding zinc finger clusters (ZFCs) in each protein has been demonstrated to preferentially recognize hmC-containing DNA *in vitro*. Yet the impact of the remaining 2-3 ZFCs that lack hmC specificity on the overall binding properties of these proteins was not explored, although duplex pulldowns did not indicate significant discrimination between hmC and the more abundant mC modification (Xiong et al., 2016). Moreover, as these experiments were performed by a qualitative EMSA readout (blotting and ECL) with a biotinylated DNA fragment bearing 15 uniformly substituted 5-position modified CpG's, such that the specificity for a single hmC CpG dinucleotide is unclear. Yet given the low abundance of these modifications, this is a more realistic substrate. We expressed and purified GST-fusions of the implicated hmC-specific SALL1-ZFC3 (982-1063), and the much more modestly selective SALL4-ZFC1 (376-461), as well as the completely 5-postion non-specific SALL1-ZFC4 (1115-1202) as previous authors (Xiong et al., 2016). We also expressed and purified a construct that spans all known DNA binding modules of the SALL1 protein ZFC1-ZFC4 (438-1202) as a GST fusion, to examine the properties of these domains with different selectivities on the overall binding preferences of the protein. This construct was proteolytically sensitive, but could be purified by the rapid sequence of GST-trap, Heparin Hi-Trap, then Superdex S200 chromatography to afford useable homogeneity for EMSA assays. We recapitulate a modest preference for hmC with the SALL1-ZFC3, yet do not observe any specificity for the other two SALL1/4 ZFCs (Figure S1I). Importantly, the non-specific binding of SALL1-ZFC4 is at least 50-fold higher than that observed for SALL1-ZFC3 engaging hmC-DNA. This is consistent with our

finding of no apparent specificity in the context of the larger SALL1 ZFC1-ZFC4 construct (Figure S1J).

#### Double filter binding assays with recombinant proteins

For THY28, 70 μL binding reactions were setup in a 96-well microplate format as follows: 30 μL of THY28 FBAI was added to 20 μL of binding buffer (20 mM Na·HEPES pH 7.8, 150 mM NaCl) and mixed by repeated pipetting ("row 2"); the resulting solution was serially diluted into 9 additional rows via the same 30 μL removal volume / 20 μL initial buffer volume scheme; following dilution, 10 μL of binding buffer was added to rows 2–11; rows 1 and 12 were prepared by addition of solely 30 μL of THY28 FBAI or binding buffer, respectively; the microplate was then briefly spun down at 1500 rpm (Sorvall RC-3B H-6000A rotor); lastly, 40 μL of radiolabeled DNA, diluted from frozen stock to achieve a final concentration of 1 nM in the binding reaction, was added to each row, and the resulting solution was mixed by pipetting. After a 30-minute incubation at room temperature, reaction mixtures were transferred to the assembled FBA apparatus, as described below. For C3ORF37 FL, 50 μL binding reactions were setup as described above for THY28 with some differences, as follows: rows 1 and 2 were prepared by addition of 36 μL of C3ORF37 FL FBAI to 4 μL of binding buffer, or 17 μL of FBAI to 23 μL of binding buffer, respectively; rows 3–11 were prepared via a 20 μL removal volume / 10 μL initial buffer volume serial dilution scheme; radiolabeled DNA was delivered in 10 μL to achieve a final concentration of 1 nM. For GST-C3ORF37 NTD, 80 μL binding reactions were setup as described above for THY28 with some differences, as follows: rows 1 and 2 were prepared by addition of solely 70 μL of GST-C3ORF37 NTD FBAI-1, or 55 μL of FBAI-1 to 15 μL of binding buffer, respectively; rows 3–11 were prepared via a 110 μL removal volume / 70 μL initial buffer volume serial dilution scheme; radiolabeled DNA was delivered in 10 μL to achieve a final concentration of 1 nM. For THAP11, 100 μL binding reactions were setup as described above for THY28 with some differences, as follows: rows 1–11 were initially prepared via a 55 μL removal volume / 33 μL initial buffer volume serial dilution scheme, after which an additional 15 μL of THAP11 FBAI diluted with 15 μL of binding buffer was added to row 1 and 30 μL of binding buffer was added to rows 2–11; radiolabeled DNA was delivered in 37 μL to achieve a final concentration of 1 nM.

Filter binding assays (FBA) were performed using the 96-well Bio-Dot Microfiltration Apparatus (Bio-Rad) with Amersham Protran 0.1 μm NC nitrocellulose membrane (GE Healthcare) on top of a Zeta-Probe membrane (Bio-Rad). Prior to apparatus assembly, membranes were well-equilibrated in binding buffer (20 mM Na·HEPES pH 7.8, 150 mM NaCl). The apparatus was assembled by sandwiching, in the order encountered by applied sample, the nitrocellulose membrane, Zeta-Probe membrane, and 2 dry Whatman blotting papers (Grade 3MM Chr). Immediately prior to sample application, 100 μL of binding buffer was applied to each position, and a vacuum of approximately –20 inHg was temporarily applied to evacuate liquid from each well. Reaction mixtures were then similarly applied and drained, after which a single 100 μL wash with binding buffer was performed. Following complete drainage of the wash, vacuum was applied for an additional 2–3 minutes before disassembly of the apparatus to facilitate drying of the membranes. Disassembled membranes and Whatman filter papers were exposed to a Fujifilm "CR" phosphorimaging screen, which was subsequently scanned at 100 nm resolution on a Typhoon 9200 (GE Healthcare).

Signal quantitation of nitrocellulose and Zeta-Probe membranes was performed using TotalLab Quant Array Analysis of .GEL files generated by the Typhoon 9200 system. For each spot on both membranes, automatic background subtraction was applied to the observed signal using the spot edge average method to generate a raw counts value. Fraction DNA bound ($\theta$) was calculated for each spot in a given replicate as follows:

$$\theta = [S_N/X]/[(S_N/X) + (S_Z/Y)] - [S_{DNA\ only,N}/X]/[(S_{DNA\ only,N}/X) + (S_{DNA\ only,Z}/Y)]$$

where $S$ is the raw counts value and the indices $N$ and $Z$ correspond to the nitrocellulose and Zeta-Probe membranes, respectively; $X$ and $Y$ are normalizing factors given by $X = S_{max,N} - S_{DNA\ only,N}$ and $Y = S_{DNA\ only,Z} - S_{min,Z}$ for the nitrocellulose and Zeta-Probe membranes, respectively, where $S_{max}$ and $S_{min}$ are the maximum and minimum raw counts value observed within a given replicate for the respective membrane, and $S_{DNA\ only}$ is the raw counts value of the DNA only spot (row 12) of that replicate for the respective membrane. Plots of $\theta$ vs. protein concentration shown in Figure 1B and S1D were generated using KaleidaGraph v4.5.2 (Synergy Software) from averaged $\theta$ values across all replicates of a single FBA experiment for a given DNA modification state, and vertical error bars correspond to a standard deviation measured values. For THAP11, unaveraged $\theta$ values for a given DNA modification state obtained from all replicates of a single FBA experiment were plotted versus final protein concentration and fit to the Hill function *m3*((m0^m2)/((m0^m2)+(m1^m2)))* using the General Curve Fit functionality of KaleidaGraph v4.5.2 (Synergy Software), where *m0* is the concentration of free protein (assumed to be equal to the concentration of total protein), *m1* is the dissociation constant $K_d$, *m2* is the Hill coefficient, and *m3* is the saturation point. Each $K_d$ value is displayed ± 2*S.E., where the S.E. reflects standard error in data fits.

#### Isolation of nuclei and nuclear extract from porcine cerebrum

Whole brains were taken from recently sacrificed adult pigs of indeterminate agricultural breeds. The whole brains were briefly washed in PBS and then coarsely dissected to remove the brain stem and cerebellum from the cerebrum for separate processing. The meninges and external vasculature were removed before finely mincing the cerebral cortex. This tissue in 200 g batches was suspended in 2 L of homogenization buffer (20 mM Na·HEPES pH 7.9, 30 mM KCl, 1 mM EDTA, 1 M sucrose (34% w/v), 10% glycerol (v/v)). This suspension was then processed twice through a Yamato LH-21 continuous flow homogenizer operating at 150-180 RPM at a flow rate of ~20 mL per minute. The cell homogenate was centrifuged in 500 mL bottles to pellet the intact nuclei (Sorvall RC5B with Fiberlite F10-6 x 500y rotor at 16,000xg for 20 minutes, 4°C). After decanting the supernatant, and sloughing off excess lipid residue with paper towels, recovered crude nuclei were then resuspended by pipette in 200 mL of buffer per 500 mL of lysate pellet in a hypotonic reduced sucrose buffer (20 mM Na·HEPES pH 7.9, 10 mM KCl, 1 mM EDTA, 340 mM sucrose (10% w/v), 10% glycerol (v/v)) and loaded onto a 30% (w/v) sucrose cushion (30% sucrose variation of the same hypotonic sucrose buffer, 100 mL of cushion in 750 mL Bio-Bottle (Thermo Scientific). Gentle centrifugation of the nuclei through the cushion (Sorvall Legend XTR with TX-750 rotor at 600 x g for 10 minutes, 4°C) serves to wash the nuclei and remove lipid carry-over from the cell lysis. The purified pelleted nuclei were resuspended in three packed nuclear pellet volumes of the hypotonic reduced sucrose buffer supplemented with 1 mM PMSF and 5 mM β-mercaptoethanol, and flash frozen for storage at -80°C.

Nuclear extract was prepared as previously described (Parker and Topol, 1984) with some modifications. Briefly, nuclear pellets in 50 mL conical tubes were thawed on ice, gently resuspended, and supplemented with 200x Protease Inhibitor Cocktail in DMSO (concentrations at 1x: 1mM AEBSF, 0.8 μM aprotinin, 20 μM leupeptin, 15 μM pepstatin A, 40 μM bestatin, 15 μM E-64). Saturated ammonium sulfate stock was added dropwise to 400 mM final concentration and the mixture was incubated with gentle rocking at 4°C for 30 minutes. The nuclear extract was then clarified by ultracentrifugation (Beckman Ti70.1 rotor at 37,000 rpm for 1.5 hours, 4°C). The soluble nuclear extract was filtered through a 0.45 μm mixed cellulose syringe filter (Millipore) prior to dilution with 50 mM Bis-Tris pH 6.2, 1 mM EDTA, 10% glycerol (v/v) (v/v), 5 mM β-mercaptoethanol (BTEG0) to lower the salt in preparation for ion exchange chromatography.

#### Column chromatography

The chromatography steps outlined in Figure 1E represent a highly refined path to isolating the major [ox]mC-binding activity observed and characterized here. The sequence of columns was optimized through much trial and error, by experimenting with the running conditions to afford the greatest amount of enrichment of [ox]mC specific activity in as few and distinct steps as possible. After each column, EMSAs were used to assess the DNA binding activity and interrogate its properties through variation of the labeled oligonucleotide, as well as cold competitor ratio and identity. Fractions that exhibit specific shift for [ox]mC over mC and C cold competitors were then pooled and subjected to further characterization by EMSA and chromatographic fractionation. The exact running conditions, were sometimes proportionally adjusted from prep-to-prep in order to account for changes in the scale of the prep. From point of nuclear lysis onward, freezing the partially purified activity significantly diminished the strength and specificity of the ox-mC specific activity. As such, extract and fractions were never frozen and purification to mass spectrometry inputs was performed as rapidly as possible, typically within ten to fourteen days after nuclear lysis.

Due to the overwhelming abundance of DNA-binding factors that are non-specific for 5-position modifications or preferentially bind mC-containing duplexes, these factors dominate the bulk binding activity of crude nuclear extract. Thus, affinity purification methods that proceed directly from nuclear extract are largely saturated with non-[ox]mC-specific binding proteins (Iurlaro et al., 2013; Spruijt et al., 2013). It takes several optimized chromatographic steps for our purification scheme to enrich the [ox]mC activities enough to separate the [ox]mC-specific activity from all other DNA binding activities. Prior to this enrichment, even within the partially purified extract, non-specific binding activities will be recovered by [ox]mC pulldowns because this pool of proteins lacks the usual polyanion sink of the genome to which it normally binds, and will bind any DNA available to a certain extent in its absence.

*20 mL POROS Heparin*

To load the 20 mL bed volume POROS Heparin (Applied Biosystems resin packed in XK-16 column, GE Healthcare), the salt concentration and pH of the soluble nuclear extract was first lowered by dilution in BTEG0 (50 mM Bis-Tris pH 6.2, 1 mM EDTA, 10% glycerol (v/v), 5 mM β-mercaptoethanol) until the extract matched the pH and conductivity of BTEG250 (BTEG0 supplemented with 250 mM NaCl). After two column volumes of wash in BTEG250 on an Akta Purifier (GE Healthcare), the column was developed first with a linear gradient to 600 mM NaCl over four column volumes, then by a linear gradient to 2M NaCl over one column volume, and held at this salt for another column volume. 5 mL fractions were collected during the shallow gradient and the [ox]mC-binding activity in each was examined by a gel shift. For this and other columns, the initial EMSA (described below) screened all column fractions, washes and input for preferential binding activity for radiolabeled hmC/hmC as compared to radiolabeled mC or C in the presence of mC or C cold competitors. hmC-specific fractions also shift mC to a lesser extent under these conditions, demonstrating that there are factors that preferentially bind hmC over mC in the mammalian brain. Whereas, in the presence of mC cold competitor, the shifted bands are more intense as compared to C cold competitor, suggesting that mC is an even less preferred binding partner than C at this stage in this preparation. The fractions that exhibit hmC-specific shift in this first EMSA are then subjected to further characterization by EMSA using other oligonucleotide presentations and a variety of cold competitor identities and concentrations. An hmC specific activity elutes later in the gradient (Figure 1D, S1A-C, fractions 12, 13, and 14) than the early large peak of protein which does not bind hmC-containing DNA (data not shown). From the first column forward, the fC-specific binding is typically somewhat “stronger” than hmC specific activity, in that a greater amount of oligonucleotide is shifted for the same amount on fraction input and labeled oligonucleotide. The fC-specific shift also has a slightly higher electrophoretic mobility (Figure 2C and F; Figure S2F).

*6 mL Resource S*

Active fractions from the POROS Heparin column were pooled and diluted with BTEG0 to equivalent conductivity with BTEG200 (BTEG0 supplemented with 200 mM NaCl), and loaded onto a 6 mL Resource S column (GE Healthcare), pre-equilibrated in BTEG200. After loading via superloop, and washing with two column volumes with BTEG200, the column was developed with a linear gradient from 200 to 750 mM NaCl over six column volumes then washed by linear gradient over 1 column volume to BTEG1500 (BTEG with 1.5M NaCl) and held in this buffer for a column volume thereafter (Figure S2A). The fractions eluted in the fifth column volume exhibit [ox]mC-specific activity (Figure S2E). This activity is observed as shifts of several mobilities for hmC, suggesting multiple activities or the decomposition of a complex activity into several components as a result of this chromatography. The major activity is the highest mobility activity, though lower mobility activities are also revealed by the Resource S column that are not enriched in the input. These activities were not found to be significantly different in their specificity for [ox]mC at this stage, and fractions containing both the high mobility activity and the low mobility activities were combined for further fractionation.

*1 mL Mono S*

Active fractions from the Resource S column were diluted with BTEG0 to iso-conductivity with BTEG200. The column was loaded by superloop and washed for two column volumes, then developed over sixteen column volumes to 600 mM NaCl, then over 2 column volumes to BTEG1200 (BTEG with 1.2M NaCl). Fractions (0.3 mL) were collected across the shallow gradient elution (Figure S2A). In the gel shift assays, the binding activity of every other fraction for hmC was screened in the presence of C and mC cold competitor. The fractions eluted in the fifth to seventh column volumes of the gradient exhibit the strongest [ox]mC specific activity in the gel shift assay, as shown in Figure 2C and S2F. In Figure S2F (top panel), fractions less than a column volume apart (0.3 mL) show differential enrichment in a lower mobility activity (compare fractions 20 and 22). The hmC-binding activity first appears in fraction 18 and the bulk of this mobility activity tails into the 20s, substantially co-fractionating with a lower electrophoretic mobility activity (the distinction is more clearly apparent in middle panel of Figure S3F). After this chromatography step, this lower mobility activity shows less discrimination between [ox]mC states and asymmetric presentations and also exhibits less sensitivity to hmC as a specific competitor as compared to the higher mobility activity. This higher mobility activity is remarkably sensitive to the symmetry of [ox]mC: [ox]mC juxtaposed with unmodified C is effectively bound (hmC/C and fC/C), whereas we detect little binding of [ox]mC/mC counterparts (hmC/mC and fC/mC) under these conditions (Figure 2F right panel; S2F middle panel). The higher mobility shift is more robust in the presence of C cold competitor as compared to mC, which while different from the activity previously observed at the POROS Heparin stage (Figure S2C), indicates a change in the aggregate preferences as the composition and relative abundances of individual factors within the protein pool change via purification. In pursuit of the major and stronger high mobility activity, fractions more clearly enriched in this higher mobility activity were pooled together for further fractionation, whereas fractions with the low mobility activity were pooled separated and processed secondarily. This observation by EMSA of separation of the two activities by this step is very sensitive to the loading level of the column (more material conflates the two) and the percentage of the gel used in the EMSA (compare top and middle panels of Extended Data Figure 2F).

*3.3 mL DEAE-5PW*

Fractions from the Mono S column were diluted in 100 mM Tris pH 8.0, 1 mM EDTA, 50 mM NaCl, 5% glycerol (v/v), and 5 mM β-mercaptoethanol (TEG0) to iso-conductivity with TEG50 (50 mM NaCl). A 3.3 mL TSKgel DEAE-5PW column (Tosoh Biosciences) was equilibrated in TEG50, loaded, and eluted with a linear gradient over 5 column volumes to 1.2 M NaCl with 0.2 mL fractions collected (Figure 2A). The fractions eluted in the first peak exhibit the strongest [ox]mC specific activity in the gel shift assay, as well as a marked concentration of the activity by this column. This column can partially chromatographically separate the high and low mobility activities, as shown in Figure S2G. These activities again show slightly different specificity for [ox]mC, with the high mobility activity being qualitatively stronger and more specific. Residual activity is still present in the flow-through, consistent with column saturation (Figure S2G). Fractions containing predominantly the high mobility activity were carried forward to the size exclusion column and ultimately to the duplex oligonucleotide pull-down and mass spectrometry experiments. However, the low mobility activity was also further characterized through size exclusion chromatography and EMSA, or by reloading the DEAE-5PW for a second round of capture and separation (data not shown).

*Superdex 75/200 (24 mL 10/300GL) size exclusion*
Both size exclusion columns were equilibrated and developed in 50 mM Na·HEPES pH 7.8, 1 mM EDTA, 300 mM NaCl, 5% glycerol (v/v), and 5 mM β-mercaptoethanol. Fractions from the previous column enriched in the high mobility activity were concentrated on a 10 kDa MWCO Ultrafree centrifugal concentrator (Millipore) prior to loading in 0.5-1 mL injection volume. Fractions (0.3 mL) were collected. The [ox]mC specific activity elutes at ~12.5 mL retention volume on the Superdex 200 (Figure 1A, B and D; S2A and H-K) and ~8 mL on the Superdex 75 (near void volume, data not shown). Within the major high mobility shifted complex bands, there are still two sub-bands visible at the beginning of the elution of [ox]mC specific activities, and these can be partially separated in some preparations that are more enriched for the lower mobility activity. Fractions from both of these columns were employed in solution MS/MS (Figure S3A), and the higher mobility activity pools were also subjected to duplex DNA pull-downs coupled to MS/MS (Figure 3A and B).

#### Electrophoretic mobility shift assays (EMSA)
Binding reactions (10µL) were set up as follows: 2 µL 5x binding buffer (where the final 1x concentration is 20 mM Na·HEPES pH 7.8, 10 mM $(NH_4)_2SO_4$, 30 mM KCl, 1 mM EDTA, 0.2 % Tween 20, supplemented with 0.1 mM PMSF, 5 mM β-mercaptoethanol, and including cold competitor as needed), 1-6 µL chromatographic fraction (remaining volume is current column input buffer), and 1 µL radiolabeled DNA (typically 50 nM), added last. The binding reactions were incubated at room temperature for 20 minutes, then 1 µL of loading buffer is added (20 mM Tris·HCl, pH 7.5, 20% glycerol (v/v), 0.02% bromophenol blue, 0.02% orange G). The binding reactions (4-10 µL) are then loaded into a 19:1 polyacrylamide gel pre-equilibrated and running at 4°C in 0.25x TBE for 30-60 minutes at 8 mA. The percentage of the gel varied between 4-7% depending on the mobility of the activity and the resolution desired. The volume of chromatographic fraction used was adjusted to allow for optimal gel loading (minimal hold-up in wells) and sensitivity (resolution of bands with discrete mobilities). Each gel was dried on Whatman blotting papers (Grade 3MM Chr) on a Bio-Rad gel drier before overnight exposure (12-18 hours) to a “CR” phosphorimaging plate (Fujifilm), and then scanned on a Typhoon 9200 (GE Healthcare) at 100 nm resolution.

#### Oligonucleotide pull-downs
As a final enrichment step, we performed comparative affinity capture with immobilized duplex oligonucleotides from highly purified size exclusion fractions. This affinity purification procedure

has two advantages over protein identification directly from the size exclusion fractions. Foremost, it allows for enrichment of low abundance proteins within the active fractions that may contribute greatly to EMSA activity, that might not otherwise be detected in solution mass spectrometry samples without affinity capture, washing, and enrichment. The enrichment of these proteins through several washing steps also represents a more stringent protein identification as compared to direct fractions, as the pull-down represents multiple captures through dilution and re-equilibration between the binding partners. Secondarily, by performing pull-downs with different substrates, we can potentially differentiate symmetric, asymmetric, hmC and fC binding activities.

Symmetric and asymmetric presentations of C, mC and [ox]mC were used as immobilized affinity reagents on streptavidin M280 Dynabeads (Life Technologies). Following pre-equilibration in 20mM Tris·HCl pH 7.8, 1M NaCl, 2 mM DTT, this paramagnetic resin was incubated with 600 pmol of biotin-labeled DNA per mg of beads (this amount represents at least 3-fold the calculated binding capacity of the resin for short oligonucleotides). These duplex biotinylated oligonucleotides were annealed as described previously but with a 10% molar excess of the non-biotinylated strand to ensure that all the immobilized DNA is double-stranded. This oligonucleotide-capture incubation proceeded for 1 hour at room temperature with gentle rotation in siliconized tubes. The beads were then washed six times with three tube changes for ten minutes in five times the original bead slurry volume with 20mM Tris·HCl pH 7.8, 1M NaCl, 2 mM DTT. These loading conditions yield equivalent oligonucleotide attachment across preparations as assessed by PNK radiolabeling of 5ug of beads and denaturing gel electrophoresis (Figure 4A inset). Prior to use, these immobilized duplex beads were resuspended at the original concentration of 10 mg/mL in 50 mM Na·HEPES pH 7.8, 300 mM NaCl, 5% glycerol (v/v), and 5 mM β- mercaptoethanol.

Size exclusion fractions with peak [ox]mC binding activity were pooled and incubated with 100 ug prepared bead slurry. All comparisons were performed with side-by-side handling controls, for example, a given volume of an active fraction pool compared by probing with equivalent amounts of immobilized hmC/hmC- or mC/mC-resin. The beads were incubated with the active fractions for 60 minutes in siliconized tubes and washed by separating the beads from the solution by setting the tubes in a neodymium magnetic rack (Life Technologies) and removing the supernatant. Beads were washed three times for ten minutes with two tube changes with 0.5 mL of 50 mM Na·HEPES pH 7.8, 400 mM NaCl, 5% glycerol (v/v), and 5 mM β-mercaptoethanol. Washing conditions were screened, varying salt concentration, number, and length of washes, however, this did not substantially change the results. Proteins retained by the beads were eluted with 12 µL of 4x SDS-PAGE loading buffer (1x = 50 mM Tris·HCl pH 6.8, 2% SDS (w/v), 10% glycerol (v/v), 0.2 % bromophenol blue and 5 mM β-mercaptoethanol) and boiled for five minutes before loading onto a Bis-Tris 4-20% NuPAGE (Life Technologies) gradient SDS-PAGE gel in 1x MOPS-SDS running buffer. The gels were then stained by standard procedures using the Pierce Silver Quest silver staining kit (Figure 3A). Corresponding gel slices in parallel bead only, C/C, mC/mC and [ox]mC/mC pull-down lanes were also excised to compare use as negative controls in the mass spectrometry analysis.

#### Mass spectrometry

*Sample preparation*

Individual gel bands were digested with the In-Gel Tryptic Digestion kit (Thermo Fisher) according to the manufacturer's instructions. Briefly, disulfide bonds were reduced by incubation with 3 mM TCEP for 20 minutes at 30°C, followed by alkylation with 12 mM iodoacetamide for 15 minutes at 30°C. Gel bands were then shrunken and dried with addition of 50% acetonitrile. Each gel band was digested with 0.1 ug trypsin for 4 hours at 37°C, then for 16 hours at 30°C. This material was filtered in 0.45 µm centrifugal spin filters (Millipore), and protonated by addition of 5% formic acid prior to LC-MS loading.

Twenty size exclusion fractions were prepared for mass spectrometry analysis: 10 fractions from each size exclusion column, composed of 5 [ox]mC-active fractions and 5 [ox]mC inactive Individual size exclusion fractions were digested using reagents from the In-Gel Tryptic Digestion kit from Thermo Fisher Scientific. Briefly, disulfide bonds were reduced by incubation with 3 mM TCEP for 20 minutes at 30°C, followed by alkylation with 12 mM iodoacetamide for 15 minutes at 30°C shielded from light. 20uL of size exclusion fraction (<1 µg total protein) was digested with 0.1 µg trypsin for 3 hours at 37°C, then for 16 hours at 30°C. This material was filtered in 0.45 µm Ultrafree-MC HV Centrifugal Filter (Millipore) and protonated by addition of 5% formic acid prior to LC-MS loading.

*Mass spectrometry measurements*
Liquid chromatography-mass spectrometry and data analyses of the digested samples were carried out as previously described(Keller et al., 2002) . Briefly, all experiments were performed on a Dionex Ultimate 3000 Nano-HPLC, coupled to a linear ion trap Orbitrap Velos Pro mass spectrometer (Thermo Fisher Scientific, Bremen, Germany) equipped with a nanoelectrospray ion source mounted with New Objective uncoated silica tip electrospray emitters, 260/75 µm OD/ID, 8 µm tip ID. For HPLC separations, buffer A consisted of 5% acetonitrile, 0.1% formic acid in water and buffer B consisted of 95% acetonitrile and 0.1% formic acid in water. For the LC-MS/MS experiments, digested peptides were directly loaded at a flow rate of 500 nL/minute onto an Acclaim PepMap 100 C18 LC trapping cartridge (Thermo Scientific) and then onto a Zorbax 300 SB C18 3.5 µm x 150 mm x 75 µm LC column (Agilent Technoligies). The column was enclosed in a column heater operating at 30°C. After 30 minutes of loading time, the peptides were separated with a 90-minute gradient at a flow rate of 350 nL/minute. The gradient was as follows: 0–5% buffer B (5 minutes), 5-45% buffer B (60 minutes), and 100% buffer B (5 minutes). The Orbitrap was operated in data-dependent acquisition mode to automatically alternate between a survey scan (m/z 5, 400–1,600) in the Orbitrap and 20 collision-induced dissociation (CID) MS/MS scans in the linear ion trap. CID was performed with helium as the collision gas at a normalized collision energy of 35% and 10 ms of activation time.

*Mass spectrometry data analysis*
Thermo RAW files were converted to MGF file for searches with Mascot (Matrix Science). Spectra were searched against all NCBI nr mammalia taxonomy database (version 20131113, 2037278 entries). Allowed variable modifications, as defined by Unimod (followed by monoisotopic mass), included oxidation of Met (115.9949), Carboxyamidomethylation (57.0215), deamination of Asn and Gln and formylation (0.984016), and formylation (27.994915).

The mass accuracy for ESI-FT-ICR by the Orbitrap Velos Pro in Mascot searches was set as 10 ppm/0.02 Da and the fragment mass tolerance was set at +/- 0.6 Da, with peptide charges of 2+, 3+ and 4+ allowed for the monoisotopic mass. Trypsin cleavage tolerance was set at +/- 1 missed cleavages. Individual mass spectrometry runs were inspected in XCalibur Qual Browser (Thermo Scientific). Mascot search output DAT files were analyzed using Scaffold (version 4, Proteome Software) to compare protein identifications between the different oligonucleotide pull-down and solution mass spectrometry experiments with peptide identification threshold set at 90% (3% FDR) in Scaffold (calculated according to Peptide Prophet algorithm (Keller et al., 2002) ). At 95% confidence (1% FDR), 11 observations of WDR76 are made in three pull-downs using hmC/hmC, fC/fC, and hmC/fC.

In total, all of the native brain purification gel shift assays, the oligonucleotide pull-down experiments, and the mass spectrometry analyses presented herein reflect biochemical isolation of three different batches of porcine brains, six distinct chromatographic preparations, 20 solution mass spectrometry experiments, and 48 oligonucleotide pull-down gel bands digests arising from two of the fractionation replicates through two different final size exclusion columns.

##### Expression and purification of recombinant WDR76

Human WDR76 was cloned by Gibson assembly of 3 codon-optimized synthetic fragments based on the sequence of the long isoform (Genbank Accession BC025247). We attempted to express this protein in *E. coli*, and observed little soluble full-length expression, and specific, but limited activity by EMSA. By sequence homology and secondary structure prediction, WDR76 exhibits striking similarity to DDB2 in its C-terminal WD repeat region (amino acids 194-560). The N-terminal region is distinct and is predicted to be alpha helical (approximately amino acids 1-236). In order to begin to dissect the molecular basis of hmC-specific binding by WDR76, the contribution of the N-terminal alpha-helical region and the C-terminal WD repeat region were assessed separately (Figure S3F). We also designed 4 C-terminal constructs encompassing the WD repeat region, none of which were soluble under multiple expression conditions, including various hosts and chaperone-assisted expression hosts. Three N-terminal constructs were also designed and found to be soluble, but did not exhibit full specificity for [ox]mC (Figure S3F-H). These three constructs were designed to span the N-terminal region from the first codon to a varying the C-terminal boundary, and cloned into pMSCG7 with an N-terminal polyhistidine tag. All three exhibited high soluble expression in *E. coli* and were purified by Ni-NTA, ion exchange, and gel filtration chromatography. The highest yield and purity was observed from a construct spanning amino acids 1-165. The secondary structure of this N-terminal construct was assessed by CD spectroscopy using a J-715 spectrometer (JASCO Corporation) scanning from 190 to 250 nm (Figure S3G). The binding properties of this N-terminal construct were assessed in gel shift assays filter binding assays as described above (Figure S3H).

In insect cell expression systems, we attempted to make both the full length and complete WD repeat region C-terminal construct from both human and mouse sequences, all with various affinity and solubility tags. The best-behaved soluble protein was the full-length human WDR76 sequence, although a C-terminal activity displayed weak modestly specific activity (Figure S3I). To produce recombinant protein for these studies, the human WDR76 construct was cloned into pFastBac with an N-terminal FLAG-$His_6$-$Arg_6$-tag cleavable with human rhinovirus R3C protease. This construct was transformed into DH10Bac cells and clones containing recombinant bacmid DNA were isolated and screened by PCR for proper incorporation of the construct into the bacmid. *S. frugiperda* (Sf9) cells were transfected with validated bacmid DNA using FuGENE HD (Promega), and baculovirus was amplified and isolated for large scale infections of *T. ni* Hi5 (Invitrogen) cells. Sf9 cells were grown in Sf-900 II SFM media (Gibco) supplemented with 10% FBS (v/v), 2 mM L-glutamine, 50 mg/L gentamicin, 10 units/mL penicillin, and 10 mg/L streptomycin. Hi5 cells were grown in Insect-XPRESS Protein-Free Insect Cell Medium with L-glutamine (Lonza) supplemented with an additional 2 mM L-glutamine, 50 mg/L gentamicin, 10 units/mL penicillin, and 10 mg/L streptomycin. Both cell types were grown in 2L fernbach flasks in a platform shaker (Kuhner) rotating at 100 rpm at 27°C.

Cells were collected by centrifugation 72 hours post-infection with baculovirus (Sorvall Legend XTR with TX-750 rotor at 500 x g for 10 minutes, 4°C) and each liter of culture was resuspended in 60 mL HEGN600 (50 mM Na·HEPES pH 7.8, pH 7.8, 1 mM EDTA, 10% glycerol (v/v), 0.02% NP-40, 600 mM NaCl) and supplemented with protease inhibitor cocktail. Cell slurry was flash-frozen in liquid nitrogen and stored at -80°C until purification.

Thawed cell slurries were lysed in a Dounce homogenizer with 30 tight pestle strokes. Lysis was confirmed under the microscope with Trypan Blue staining. The lysate was clarified by centrifugation (Sorvall RC5B with SS-34 rotor at 30,000xg for 25 minutes, 4°C), the salt was lowered to 200 mM by dilution with supplemented HEGN0 (no NaCl), and the centrifugation step was repeated. This clarified extract was incubated with 250 µL of FLAG M2 agarose affinity gel (Sigma-Aldrich) per liter of culture input for 1 hour, rotating at 4°C. The flow-through was collected by centrifugation at 500 x g for 5 minutes. The resin was washed in 10 resin volumes of HEGN600 for 10 minutes, and then in HEGN300 for 10 minutes, rotating at 4°C. The FLAG-fusion protein was eluted by incubating the resin in 0.5mL of HEGN300 supplemented with 150 µg/mL 3x FLAG peptide for 30 minutes, rotating at 4°C. Six resin volumes of elution were collected.

Pooled elutions were visualized by SDS-PAGE before pooling for further purification on a 1mL POROS Heparin column (Applied Biosystems resin packed in Tricorn 10/50 column, GE Healthcare). The pooled elutions were diluted in BTEG0 to iso-conductivity with BTEG150, loaded onto the column, the column washed with four column volumes BTEG150, then developed in a step gradient to 20% BTEG1000 with a four column volume hold, and then to 100% BTEG1000 over six column volumes. This purification step serves to remove the excess FLAG peptide and reduce the amount of co-purifying nucleic acid carried with the protein from the insect cells. The eluted material was visualized by SDS-PAGE and quantified by $A_{280}$ ($\varepsilon$=42,860 L $M^{-1}$ $cm^{-1}$) for use in gel shift assays.

In an effort to enhance the limited specific activity of the recombinant WDR76, protein purified by the 1 mL POROS Heparin was also purified over the 0.36 mL DEAE 5-PW. The input was diluted in TEG0 (100 mM Tris pH 8.0, 1 mM EDTA, 50 mM NaCl, 5% glycerol (v/v), and 5 mM $\beta$-mercaptoethanol) to iso-conductivity with TEG50 (Buffer A). The input was loaded by superloop, washed with 10 column volumes of 20% TEG1000 (Buffer B) and eluted over a 5 column volume linear gradient to 100% Buffer B. The eluted material is again visualized by SDS-PAGE and quantified by $A_{280}$ ($\varepsilon$=42,860 L $M^{-1}$ $cm^{-1}$) for use in gel shift assays (Figure S3D). The specific activity of WDR76 in EMSAs was not substantially enhanced by this or other types of chromatography tested (MonoQ and Superdex 75).

#### Generation of stable WDR76 HEK293 cell lines

The WDR76 construct described above was sub-cloned into a pCDNA5-derived mammalian expression vector to yield an N-terminal FLAG-HA tag fusion under control of a CMV promoter, followed by a hygromycin resistance gene, and flanked by FRT donor sites. The construct was co-transfected into HEK293 FRT cells with Flp recombinase (pOG44) using FuGENE HD (Promega). Single clones were isolated and amplified under hygromycin selection of 150 µg/mL. Mutants of the full-length WDR76 construct were generated by site-directed mutagenesis or through reassembly of the Gibson assembly construct using alternative gene blocks to encode the mutations. The mutant constructs were individually transfected into HEK293 FRT with FLP recombinase and screened with hygromycin as described above. All HEK293 cell lines were grown in DMEM supplemented with 10% FBS,10 units/mL penicillin, and 10 mg/L streptomycin at 37°C in 5% $CO_2$. All HEK293 lines were validated by blotting whole cell lysates for the FLAG tag epitope (monoclonal α-FLAG-HRP, Sigma-Aldrich) or with 12CA5 monoclonal antibody serum and observation of the FLAG-HA-fusion constructs at the appropriate molecular weight (blotting procedures described below) with comparable expression levels assessed via control loading levels as visualized by MBD3 or H3 antibody blotting in lower sections of the same blot (Figure S4B).

To investigate the binding properties of the C-terminal WD repeat region of WDR76, four constructs were made across this region varying the N-terminal boundary in order to sample different predicted WD repeat propeller blades. The full C-terminal region (amino acids 191-560) was pursued further in both mammalian tissue culture (as an N-terminally tagged FLAG-HA construct introduced at an FRT site in HEK293s as described above) and in insect cells (as a FLAG-HA-polyarginine construct expressed by baculoviral transduction as described above). This C-terminal construct can be solubly expressed and purified from insect cells using a FLAG affinity purification, ion exchange and gel filtration chromatography. However, the yield is typically one-third that of full length hWDR76 from insect cells. This construct was also subjected to the gel shift and oligonucleotide pulldown assay described above (Figure S3I).

#### Subcellular fractionation of HEK293 lines

To perform oligonucleotide pull-down assays, a modified Dignam-Roeder (Dignam et al., 1983) nuclear extract and whole cell extract were prepared in parallel from HEK293 cell lines stably transfected with FLAG-HA-WDR76 constructs with commensurate expression (either wild type or 3S mutant). Specifically, equal cell numbers of wild type and mutant FLAG-HA-WDR76 lines were counted, washed in PBS, and collected by centrifugation at 500xg (Sorvall Legend XTR with TX-750 rotor) for 10 minutes at 4°C. The cells were then resuspended 3x the packed cell volume (PCV) in hypotonic buffer HB (10 mM Na·HEPES pH 7.8, 10 mM KCl, 1.5 mM $MgCl_2$, 340 mM sucrose (33% w/v), 10% glycerol (v/v), 0.5 mM PMSF, 0.5 mM DTT, and 1x protease inhibitor cocktail), then lysed by addition of an equal volume of HB supplemented with 0.2% Triton-X100 (v/v) and incubation at 4°C with gentle end-over-end rotation for 12 minutes.

To prepare whole cell extract, the salt concentration of the cell lysate was increased to 0.6M NaCl and the mixture incubated for 30 minutes at 4°C with gentle end-over-end rotation. The whole cell lysate was then clarified by centrifugation at 15,000xg for 30 minutes at 4°C (Beckman Coulter 22R microfuge). The clarified lysate was collected and the salt concentration lowered to 200 mM NaCl by dilution with HEGN0 (no NaCl). The conductivity-adjusted extract was again clarified by centrifugation at 15,000xg for 30 minutes at 4°C (Beckman Coulter 22R microfuge). This whole cell extract was the input for oligonucleotide pull-down assays.

For subcellular fractionation experiments, lysis was accomplished as above, but the cytosolic extract was recovered and subjected to clarification (Beckman Microfuge 22R, 18000xg, for 30 minutes at 4°C). The recovered nuclei were subjected to additional purification through a sucrose cushion (30% w/v sucrose, HEPES, pH 7.8, 1.5 mM $MgSO_4$), then extracted in parallel with 2 PCV of HB supplemented to 150 mM, 250 mM, or 350 mM KCl with gentle end-over-end mixing at 4°C for 1 hour to isolate soluble nuclear extract at monotonically increasing ionic strength. After extraction, the nucleoplasmic supernatants were subjected to high-speed clarification (Beckman 22R, 18000xg, for 30 minutes at 4°C) then gel samples prepared by the addition of the appropriate volume of 6x SDS-PAGE loading buffer. Whereas the nuclei were recovered by brief centrifugation (Beckman Microfuge 22R, 1200xg, 5 minutes at 4°C), and resuspended in 2 PCV of buffer HB supplemented to 1 mM $CaSO_4$ and pre-equilibrated briefly at 37°C. The chromatin pellets were then digested with 0.15 U/µL micrococcal nuclease (Worthington) for 25 minutes and the mixture quenched by the addition of 6x SDS-PAGE loading buffer.

Salt titrations of the nuclear extract were loaded for SDS-PAGE with equivalent volume, as both the soluble nuclear and chromatin pellet extracts were made from the same nuclei for a given salt concentration, and equivalent numbers of nuclei were used for each salt concentration point, the differences observed reflect partitioning between chromatin and nucleoplasm as a function of KCl (Figure 4D). The subcellular fractionation depicting cytosolic, nucleoplasmic, and chromatin levels of the protein (Figure 4C) was loaded by equivalent total protein as assayed by Bradford Assay (Bio-Rad). Gels were transferred to PVDF (Millipore, Immobilon-$P^{SQ}$) in modified Towbin's buffer (25 mM Tris, 192 mM glycine, 20% Methanol, 0.04% SDS [w/v], pH 8.0), by semi-dry apparatus (Bio-Rad Trans-Blot SD) at 200 mA/gel for 30 minutes. Blots were stained by Direct Blue 71, then blocked for at least one hour in 2% (w/v) ECL Prime blocking agent (GE Healthcare) in TBST. Primary antibodies in the same blocking solution were used as follows: 1:2000 $\alpha$-FLAG (Sigma M2-HRP conjugate), 1:5000 $\alpha$-HA(12CA5), 1:5000 $\alpha$-SP1 (Bethyl A300-134A), 1:30,000-1:150,000 $\alpha$-H3 C-term (Epicypher 13-0001), 1:5000 $\alpha$-MBD3 (Bethyl A302-529A), and 1:25,000 $\alpha$-Tubulin alpha (DM1A monoclonal). Secondary antibody HRP conjugates and M2-HRP were detected using ECL Ultra (Lumigen) on a Fuji LAS-4000 imager at sub-saturation within the linear range.

#### Oligonucleotide duplex pull-down assays from HEK293 lines

To perform oligonucleotide pull-down assays with HEK293 cell lines, 800 µL of whole cell lysate was added to 100 µg M280 streptavidin beads (Life Technolgies) pre-loaded with biotinylated modified oligonucleotides as described previously for the duplex oligonucletotide pull-down assays from fractionated pig brain nuclear extract. The lysate was incubated with the loaded beads in siliconized tubes for rotating for 1 hour at 4°C and the flow through was collected by isolation of the beads on a neodymium magnetic rack. The beads were washed with 500 µL of HEGN400 for five minutes, three times, with two intervening tube changes. The proteins retained on the beads were eluted with 10 µL of 4x SDS-PAGE buffer, resolved by gel electrophoresis, and blotted as above.

To confirm fair loading of biotinylated DNA onto the streptavidin beads used in the pull-down assay, prepared beads were subjected to radiolabeling. For each oligonucleotide type, 5 µg of prepared bead slurry was incubated with T4 PNK (NEB) overnight at 37ºC with 50 pmoles of [γ-$^{32}$P] ATP. The labeling reaction was then heated to 90ºC for 5 minutes with 1:1 addition of 95% formamide denaturing loading buffer, the loaded onto a thermally equilibrated 22% polyacrylamide gel with 6M urea and 1x TBE. After electrophoresis, the gel was dried and subjected to phosphorimaging.

#### mESC culture

All cells were grown at 37°C in humidified 5% $CO_2$ atmosphere. Unless noted otherwise, mouse embryonic stem cell (mESC) E14 lines were cultured in feeder-free conditions on 0.1% gelatin (Sigma G9391)-coated tissue culture-treated plates. +FBS/+LIF/+2i mESC medium was high glucose DMEM (GE HyClone) supplemented with 15% (v/v) heat-inactivated fetal bovine serum (Gibco 10438), 2 mM L-glutamine (Gibco), 0.1 mM non-essential amino acids (Gibco), 100 units/mL penicillin + 100 µg/mL streptomycin (Gibco), 0.1 mM β-mercaptoethanol (Gibco), 1000 units/mL LIF (Millipore ESG1107), 1 µM PD0325901 (MEK inhibitor; LC Laboratories), and 3 µM CHIR99021 (GSK inhibitor; LC Laboratories). Supplemented medium was sterile filtered (0.1 µm PES membrane), stored at 4°C, and used within 7-10 days. mESCs were routinely fed 1-2 times per day as needed by removal of conditioned medium and addition of fresh medium such that the final medium was approx. 20% (v/v) conditioned. mESCs were routinely passaged at approx. 70-80% confluency every 1-2 days as appropriate, with feeding 3 hours prior, by washing in 1X DPBS (no $Mg^{2+}$/$Ca^{2+}$), dissociation with TrypLE Express (Gibco), trituration to single cells via pipetting, quenching with three-fold excess of +FBS medium, pelleting (250xg, 5 minues, room temperature), and resuspension into approx. 60% fresh / 40% conditioned +FBS/+LIF/+2i medium. mESCs were routinely plated at approx. 6.0-7.0 x $10^4$ cells/cm$^2$ growth area.

#### Generation and characterization of *Wdr76*$^{-/-}$ mESCs

In order to assess a potential effect of WDR76 on gene expression, knockout *Wdr76* mouse E14TG2a embryonic stem cell lines were generated by CRISPR/Cas9 targeting to introduce indels that result in premature stop codons in exon 11 of *Wdr76* (Figure S5D). This site was chosen because disrupting the protein open reading frame within exon 11 truncates the protein within the central predicted WD40 repeats, such that three of the blades critical to proper β-propeller folding will not be translated (Fülöp and Jones, 1999), most likely leading to degradation (Figure S5A). Even without this putative degradation, prevention of this domain from folding would be anticipated to have an even more severe impact on DNA-binding capacity than the triple point mutant (3S mutant) used in Figure 4B.

Several guide RNAs were designed using by maximizing scores from the Zhang lab CRISPR Design tool for target fidelity (http://crispr.mit.edu) and the Liu lab SSC tool for targeting efficiency (http://cistrome.org/~hanxu/SSC/), while minimizing predicted nucleosome occupancy (http://genie.weizmann.ac.il/software/nucleo_prediction.html) (Horlbeck et al., 2016). Guides 1-3 were cloned into pX458 at BbsI sites as described (Cong et al., 2013), sequence-verified, and then transfected into E14 cells using 12 µl Lipofectamine 2000 and 2µg of plasmid per million cells. Prior to transfection, E14 mESCs were cultured to maximal doubling rate. Transfected cells were plated at low density and single colonies picked, or sorted on the FITC channel by flow cytometry (BD FACS Aria II/III) and re-plated in gelatinized 96-well plates as single cells, or the $GFP^+$ bulk sorted material was plated at low density and individual colonies were picked manually after 2-3 days, and re-plated in gelatinized 96 -well plates.

When colonies in 96-well plates approached ~30% confluence, the CRISPR-targeted cells were characterized. The entire plate is passaged into a new 96-well plate and half of the cells were collected for analysis by PCR. This material is heated to 70°C, bath sonicated and vortexed to lyse the cells, and genomic DNA was isolated using SPRI (Sera-mag speed beads, Fisher) at a ratio of 1:1 (v/v). Individual clones were initially screened by PCR from this template: a short amplicon flanking exon 11 resolved on a 2% agarose gel is diagnostic of insertions and deletions (Figure S5B). Putative bi-allelic mutants were selected from this screen, displaying two PCR products that were distinct from the parental E14 line. Several clones passed this threshold and were subjected to genotyping by TOPO-cloning the gel-purified PCR products into a pCR vector (Life Technologies). Sequencing of insert-positive bacterial colonies from candidate mESC clones of interest revealed no wild type sequence, and approximately ~30 observations of each of the two deletion alleles for clone g1-3.1 (Figure S5D). Western blot analysis with 1:1000 anti-WDR76 antibody (PA5-24147, ThermoFisher) confirmed knockout of this gene at the protein for this clonal line (Figure 5C).

The relative pluripotency of the stem cell lines created in this study was investigated by alkaline phosphatase staining of colonies. Alkaline phosphatase staining was performed using Fast Red Violet and Napthol AS-BI phosphate reagents provided in the Millipore AP Detection kit (SCR004). Cells were seeded at low density in triplicate independent experiments and subjected to different growth conditions over six days. Cells were grown in the presence and absence of mouse embryonic fibroblast (MEF) feeder cells purchased from Applied Stem Cell (Menlo Park, CA) and seeded at a density of 20,000 cells per $cm^2$ on gelatinized plates the day prior to the addition of mESCs. Cells were also grown in the presence and absence of 2i cocktail in their media in order to assess their ability to maintain normal colony morphology in the absence of these pluripotency-promoting small molecules. mESCs were initially seeded at 2,500 cells per $cm^2$. After six days, cells were subjected to AP staining and imaging. Alkaline phosphatase (AP) staining did not reveal significant differences in overall morphology or AP activity among the cell lines (Figure S5E). Growth on feeders supports smaller colonies as compared to the large dense colonies observed in the absence of MEFs after six days of culture on the same surface. In the absence of MEFs and 2i, colonies start to exhibit frayed edges of fibroblast morphology consist with preliminary differentiation.

#### Generation of epitope-tagged WDR76 mESCs

E14 mESC cells modified with an FRT recombination site were acquired from Kristian Helin's lab. To generate tagged overexpression lines for hWDR76 and hWDR76 triple positive charge mutant (termed "3S"), these constructs were cloned into a pcDNA5-derived mammalian expression vector under the control of the promoter EF1a. Cells were co-transfected using Lipofectamine 3000 (Life Technologies) and the pOG44 vector encoding Flp recombinase at a mass ratio of 40:1 pOG44:pcDNA5 at low density (2,500 cells/$cm^2$) in a 15 cm diameter plate. Cells were fed normally for the first 48 hours after transfection, after which cells were treated with 150 µg/mL Hygromycin B. Drug-supplemented media was replaced twice daily initially as many cells were dying, and then once daily as resistant colonies emerged. After 14 days of selection, individual surviving colonies were isolated and expanded under continuing drug selection. Colonies were interrogated by genomic PCR (e.g., Figure S4E) to confirm insertion and were subjected to hygromycin selection before use in experiments, and subjected to blotting to confirm expression levels (Figure S4F).

#### Chromatin immunoprecipitation of hWDR76 mESCs

*ChIP-seq*

The ChIP workflow described by Lee and colleagues (Lee et al., 2005) was used with a number of modifications. For each ChIP-seq experiment, 2.5 x $10^7$ mouse embryonic stem cells of were cultured, crosslinked, and harvested. These cells were grown in parallel at low to moderate colony density over several days prior to crosslinking. Freshly cracked 10% paraformaldehyde was used to crosslink the cells at a final concentration of 1% in 1x PBS supplemented with 1 mM $MgCl_2$ wash buffer for fifteen minutes at room temperature. Crosslinking was quenched with 1M Tris base (for a final concentration of 300 mM) for three minutes, followed by three washes in 1x PBS + 1mM $MgCl_2$. Cells were collected via a plastic spatula and passed through a 28-gauge needle to afford a coarse homogenate, then pelleted by centrifugation at 700g for five minutes, flash frozen and stored at -80˚C until used for ChIP.

Thawed cell pellets were resuspended in 10 mL of 10 mM Tris-HCl, pH 8.0, 200 mM NaCl, 1 mM EDTA, 0.5 mM EGTA, supplemented to 1x Protease Inhibitor Cocktail and incubated with gentle rocking for ten minutes at 4˚C, followed by centrifugation at 1,350g for five minutes at 4˚C. This pellet was gently resuspended in 10 mL of 50 mM HEPES-KOH, pH 7.5, 140 mM NaCl, 1 mM EDTA, 10% glycerol, 0.5% NP-40, 0.25% Triton X-100, supplemented with 200x Protease Inhibitor Cocktail to effect cell lysis. The released crude nuclei were then pelleted as before and then carefully resuspended in 6 mL of 10 mM Tris-HCl, pH 8.0, 100 mM NaCl, 1 mM EDTA,  0.5 mM EGTA, 0.1% Na-deoxycholate, 0.5% N-lauroylsarcosine, supplemented to 1x Protease Inhibitor Cocktail. This nuclei suspension was subjected to sonication to shear chromatin via Covaris S220 in TC12 glass vials (Covaris) with 600 µL of nuclei suspension and 100 mg of glass beads (0.5 mm diameter, Sigma). The Covaris S220 was used at water level 15 with 140W peak incident power (affording an actual average output of 25W) at 20% duty, for 200 cycles per burst. With these settings, the nuclear suspension was sonicated for 20s on, 10 off per cycle for 14 cycles. The batches of sheared chromatin were combined and clarified by addition of 1/10 volume of 10% Triton X-100 and centrifugation at 14,000g for 10 minutes. 100 µL of sheared clarified lysate was saved as input before proceeding to immunoprecipitation. The degree of shearing for 20 µL of clarified lysate assessed after by thermally decrosslinking with proteinase K, recovery with PCR clean column (Qiagen) and 2% agarose gel analysis to confirm uniform shearing between batches and samples with a fragment distribution between 200-700 bp and centered at 500 bp.

Excess 12CA5 was immobilized on 100 µg Protein G Dynabeads (Life Technologies) by overnight end-over-end rotation at 4°C in 1x PBS + 0.05% BSA, then washed in 1 mL of 1x PBS + 0.05% BSA three times prior to use in ChIP. Centrifuge clarified and sterile filtered supernatant collected from 12CA5 hybridomas cultured in low IgG serum at 15 µL of 12CA5 solution per 50 µL of Protein G Dynabeads. IP incubations were set up with 1 mL of clarified nuclear lysate and 100 µL of prepared dynabeads. IP occurred overnight (14-16 hours) with gentle end-over-end rotation at 4˚C in low retention 1.6 mL microcentrifuge tubes. The next day, beads were washed using a neodymium magnetic rack to collect beads and exchange buffers, with accompanying by tube changes. Beads were washed twice in 1 mL of 50 mM HEPES-KOH, 500 mM LiCl, 1 mM EDTA, 1.0% NP-40, 0.7% Na-deoxycholate (pH 7.55) for three minutes of end-over-end rotation at 4°C. This was followed by a final wash in 1 mL 1x TE + 50 mM NaCl. The IP-recovered material was eluted with 100 µL of 1x TE + 1% SDS heated at

65°C for 20 minutes. This was repeated for a final elution volume of 200 µL. Crosslinks in the eluted material were then reversed by incubation at 65°C for 8-12 hours, incubation with RNAse A (0.2 mg/mL final concentration) at 37°C for 2 hours, and incubation with proteinase K (0.2 µg/mL final concentration) at 55°C for 2 hours. The DNA was recovered with Sera-mag Speedbeads (GE Healthcare) and eluted in 1x TE, then used to generate size-selected sequencing libraries using the NEB Next library prep kit reagents. The libraries were sequenced on an Illumina HighSeq4000 at the University of Chicago Functional Genomics Core Facility. Summary statistics from the sequencing experiments can be found in Table S1.

*ChIP-seq bioinformatic analysis*
ChIP of hWDR76 WT in mESCs was aligned to the mm9 genome assembly via Bowtie (Langmead et al., 2009) using default parameters, and the resultant SAM file converted to sorted BAM using Samtools (Li et al., 2009). MACS2 was used for peak calling versus input using "broad" peak parameter and a $p < 10^{-4}$ threshold (Zhang et al., 2008) yielded 617 peaks after filtering for blacklist overlap in Bedtools (Quinlan and Hall, 2010). Alternate analyses using the smaller set (n=334) of default parameter-derived, de-blacklisted peaks from MACS2 reveal similar results. Deeptools (Ramírez et al., 2016) was used to create bedgraph and bigwig coverage of the ChIP data, both normalizing for read depth, and computing ChIP coverage as $\log_2$-fold change of depth-normalized HA-WDR76 IP relative to input.

ChIP coverage from significant de-blacklisted broad MACS2 peaks ($p < 10^{-4}$) was contoured over the top sequence motif found in our WDR76 ChIP peaks ($p < 10^{-216}$ ; binomial test comparing randomly selected background to peaks) with Homer using annotatePeaks (Heinz et al., 2010). Accompanying hmC/C profile from E14 mESCs from TAB-seq in the same E14 cell line (Yu et al., 2012) was generated by profiling their high confidence hmC occupancy dataset (FDR <5%) normalized by the frequency of C over the motif and flanking regions (+/-2kb) using annotatePeaks within the Homer Package (Heinz et al., 2010). Venn analysis comparing high confidence hmC sites from TAB-seq in the same E14 cell line (Yu et al., 2012) and the called ChIP peaks was performed with Bedtools "intersect" and "fisher" and plotted using VennDiagram package in R (Figure 4E, S4G). The density of all hmC sites (% hmC/bp) within ChIP peaks as compared to a series of genomic shuffled positions representing the same spans was performed using shuffleBED (Quinlan and Hall, 2010), and violin plotting (vioplot) and *p*-values computed via unpaired Wilcoxon-Mann-Whitney testing in R (Figure 5C, S4H, S5K; S6A).

#### WDR76 knock-down in K562 cells by CRISPRi

CRISPRi was performed in K562 cells with dCas9-KRAB integrated into the genome, generously provided by Luke Gilbert and Jonathan Weissman(Gilbert et al., 2014). K562 CRISPRi cells were cultured in RPMI-1640 (Gibco) supplemented with 10% fetal bovine serum (Gibco) and 50 units/mL penicillin + 50 µg/mL streptomycin (Gibco) at densities between 2.0 x $10^5$ – 1.0 x $10^6$ cells/mL and subjected to CRISPRi as previously described (Gilbert et al., 2014; Werner et al., 2017). In brief, guide RNAs (sgRNAs) were cloned into a modified pX330 vector (Ran et al., 2013) wherein the spCas9 was supplanted by eGFP and the sgRNA modified to possess the optimized stem loop (Chen et al., 2013). Transfection of 4 µg of each sgRNA vector into the CRISPRi-K562 line in 6-well plates using Lipofectamine 2000 (Life Technologies) followed the manufacturer's protocols. After two days, cells were re-plated on 10 cm plates, and 6 days post-transfection, $GFP^+$ cells were isolated by FACS sorting on a BD Aria II/III, pelleted, and re-suspended in 500 µLTrizol. The aqueous layer from Trizol extraction was used as input for purification with RNA Clean & Concentrator columns (Zymo), and then converted into TruSeq Illumina cDNA libraries as described below. All comparisons were made relative to three replicates transfected with an off-target negative control sgRNA ("nt") (The closest match is 15 nt to a site in protocadherin (PCDH17), that is missing a PAM sequence and is not expressed in K562 cells, referred to as "negative control-1" by its originators (Gilbert et al., 2014). Of the five sgRNA's initially screened (Figure S7A), the most effective three in depleting WDR76 mRNA were carried forward to RNA-seq (Figure S7B). Plots of average WT FPKM of genes within TAD as a function of genes significantly up- or down- regulated by knockout or CRSIPRi depletion (Figure S7C), versus the genes defined by the middle log2 fold change percentile of expressed genes (>0.25 FPKM in both conditions) with (approximately number matched to the up and down gene sets in each case) were prepared in R.

#### RNA-seq

*Preparation of libraries*

cDNA libraries were prepared from the total RNA isolated from triplicate independent cultures of wild type and *Wdr76*$^{-/-}$ (g1-3.1 clone) E14 cells grown in parallel. Briefly, cells were passaged side-by-side at similar confluence over the course of five days. Total RNA from 4 million cells of each genotype was isolated in triplicate by Trizol-chloroform extraction and purification using RNA Clean & Concentrator (Zymo). For each sample, 2 µg of total RNA was ribosome-depleted using Ribo Zero (Illumina). Prior to ribosome-depletion, four in vitro transcribed RNA standards were "spiked-in" to act as an internal calibration ladder (Lovén et al., 2012). As described in our prior work (Werner and Ruthenburg, 2015), the RNA standards are derived from yeast and bacterial genes that do significantly not map to mammalian genomes, but provide a way to calibrate the reads from sequencing of the entire library and estimate relative abundance between samples. The amount of standard added to each sample was adjusted for the relative yield of total RNA for the number of cells used. For each replicate RNA isolation from 4 million cells, the exact yield of total RNA varied between 25-30 µg. Total RNA (2 µg) was processed from each replicate, but this 2 µg represents a slightly different number of cell equivalents. The number of cell equivalents per µg of isolated total RNA was used to adjust the amount of each standard added. Standards were added to each RNA library sample at 40 copies per cell equivalent (*S. cerevisae* RAD51) 200 copies per cell equivalent (*E. coli* RNL2), 1000 copies per cell equivalent (*E. coli* MBP), and 5000 copies per cell equivalent (*S. cerevisae* SUMO).

RNA libraries were prepared from spiked, ribosome-depleted RNA using the Next Ultra Directional RNA Library Kit (NEB) and Next Multiplex index oligonucleotide primers (NEB) for use with Illumina sequencing platforms. The library was prepared according the manufacturer's instructions and amplified using 15 cycles of PCR. Indices for each library sample were chosen to minimize the Hamming distance between index reads in order to ensure proper assignment of reads to samples during sequencing. The libraries were sequenced on an Illumina HighSeq4000 at the University of Chicago Functional Genomics Core Facility. Summary statistics on RNA sequencing experiments can be found in Table S1.

*RNA-seq bioinformatic analysis*

RNA-seq fastq result files were processed using the Tuxedo Suite of software for alignment and differential expression analysis. Bowtie2.2 was used to concatenate fasta files for the RNA standards to the mouse mm9 Fa-Masked genome obtained from the UCSC genome browser. Reads were then aligned to this genome assembly using Tophat 2.1.0 (Kim et al., 2013; Trapnell et al., 2012). Transcriptomes were then assembled de novo for each replicate using Cufflinks2 (Trapnell et al., 2010) with an rRNA, snoRNA, tRNA, and 7SK mask. The three wild type mESC biological replicate transcriptome assemblies and the three *Wdr76*$^{-/-}$ biological replicate transcriptome assemblies were then merged to a single de novo transcriptome for differential expression analysis using Cuffdiff (Trapnell et al., 2013). The Cuffdiff analysis was performed using per condition cross-replicate dispersion estimation and either classic FPKM normalization in order to allow the standards to be uniquely mapped without further normalization, or using geometric mean normalization. The performance of the RNA standards in the sequencing experiment was assessed by counting the number of reads mapping to the standards in the BAM file output of Tophat using by using Samtools (Li et al., 2009). These measurements revealed consistent mapping of the standards during data processing; the number of reads observed for each standard was plotted against the number of molecules added for each standard to yield a calibration curve for each of the samples (Figure S5F). The average slopes of the standard curves for wild type and knockout samples were calculated separately and applied as a scalar to the FPKM values calculated by Cuffdiff (or raw counts per gene), resulting in scaled FPKM values that reflect a back-calculation of actual number of molecules present in the library input. Due to modest changes in RNA quantification between the internal standard-spiked library normalized analyses versus the geometric mean normalized analyses (Figure S5G), we conclude that the effects of *Wdr76* knockout are not global (Lovén et al., 2012) and therefore effectively treated using geometric mean normalization. Accordingly, we performed subsequent mESC and K562 analyses using geometric mean normalization dataset. Despite masking and blacklist filtering (https://sites.google.com/site/anshulkundaje/projects/blacklists), some ultra-high signal artifacts were apparent as large focal peaks, clearly distinct and segmented from neighboring sequence. In an attempt to restrict the differential expression analysis to genes with relatively even distribution of reads mapping across largely exonic structure, we chose to confine our differentially expressed transcript sets to Refseq genes, which largely removes these putative artifacts. From this independent triplicate analysis, we report 487 genes upregulated and 359 genes downregulated in the $Wdr76^{-/-}$ line relative to wt E14 cells using standard CuffDiff parameters (FDR < 0.05; $p < 0.003$, Benjamin-Hochmani multiple hypothesis testing). In the case of K562 CRISPRi, we treated the knockdowns with three distinct sgRNAs as replicates of a single condition as compared to the combined off-target sgRNA replicates for per-condition dispersion estimation using the GRC37 reference genome.

#### Combined bioinformatic analyses

The combined Hind III replicates or Nco I Hi-C datasets representing E14 contact domains (Dixon et al., 2012) or K562 Hi-C contact domains filtered to exclude features larger than 10Mb (Rao et al., 2014) were used for TAD calculations. Unique TADs with significantly up or down regulated Refseq genes were selected using Bedtools intersect. As there are local peaks of WDR76 signal that appear at the edges of TAD boundaries in the meta-TAD analysis (Figure 6A) and the TAD boundaries used are not high resolution, we performed overlap analysis using Bedtools intersect and "Fisher" using 10% feature length cushion appended to the TAD in all subsequent TAD analyses to more faithfully capture these data. Similar analyses without these cushions added (Figure S6C), using the alternate NcoI-TAD dataset (Figure S6D) likewise demonstrate overlap far greater than chance expectation. The converse analysis using TADs bearing significant ChIP-seq MACS2 called peaks versus genes that are differentially expressed also reveals much the same thing (Figure S6B).

De novo motif analysis was performed in Homer using "findMotifsGenome.pl" within all significant deblacklisted MACS2 peaks ($p < 0.0001$), as well as subsets of called peaks in TADs with down- or up- regulated genes in the $Wdr76^{-/-}$ line, or peaks that are not in such TADs (Figure 4F and S4I). Intriguingly, all subsets of peaks with the exception of those in TADs that lacked any significant gene expression changes had this same motif. Motif rank based on *p*-value of top five motifs is presented in bar graph form, and percentage of peaks with the motif versus background frequency are also depicted.

A combination of Cuffdiff (Trapnell et al., 2010), Bedtools 2.25 (Quinlan and Hall, 2010), CEAS (Shin et al., 2009) and shell/Awk scripting were deployed to interrogate the connections between altered gene expression in the *Wdr76* depletion experiments and the previously reported hmC TAB-seq in the same E14 cell line (Yu et al., 2012). Refseq coding genes were assessed for hmC density (computed as the sum of all %hmC within the indicated gene divided by its length) in subsets defined by significant log2 fold changes (FDR < 0.05; $p < 0.003$; using CuffDiff values), as compared to random shuffles of the KO down coordinates in Bedtools restricted to be within the set of all coding genes or defined expression quantiles, *p*-values were computed by unpaired Mann-Whitney-Wilcoxon two-tailed test in R. Bedtools intersect was used to count the number of overlapping genes from those significantly altered in the knockout, or bedtools map was used to compute the TAD hmC density in which the indicated gene sets reside. The two different E14 TAB-seq datasets (Yu et al., 2012) used, either all reported hmC sites set or the higher confidence FDR < 0.05 subset, generally recapitulated the same trends. For each, we computed hmC density as the %hmC for each site summed over a given interval (either genes or TADs), divided by its span then mapped onto the indicated genes using bedtools and awk scripting. For the purposes of applying a logarithmic function, a pseudo count of 1e-6 was added to each FPKM value (Figure S6J). A combination of CEAS (Shin et al., 2009), Homer (Heinz et al., 2010) and custom scripts were used to meta-feature contour ChIP and hmC over genes, TADs and Enhancers (Figure 4F;5D,E; 6A; S5I,L). The density of hmC within the whole interval of each TAD (Dixon et al., 2012) was computed and rank-ordered for division into the indicated quantiles (Figure 6B).

#### L-Ascorbic acid stimulation of TET oxidase activity

L-ascorbic acid has been shown to increase the rate of 5-mC oxidation by TET enzymes *in vitro* and increase global 5-hmC abundance in a TET-dependent manner in numerous cell types, including cultured mESCs (Blaschke et al., 2013). We sought to query the potential effect of such an increase in 5-hmC levels on the expression of genes observed to be differentially regulated in our *Wdr76*$^{-/-}$ mESC cell line. Following multiple passages at maximal doubling rate, *Wdr76*$^{-/-}$ and parent E14 WT mESCs were plated at 1.8 x $10^5$ cells/well in 0.1% gelatin-coated 12-well plates in quadruplicate for both treated and untreated conditions. Cells were plated in +FBS/+LIF/+2i medium comprised of 50% (v/v) respective conditioned medium and were fed by addition of fresh medium approx. 16 hours after plating. Approx. 30 hours after plating, media was completely replaced with 75% fresh / 25% conditioned medium supplemented with either 1mM HEPES•NaOH pH 7.4 (untreated) or 100 µg/mL 2-Phospho-L-ascorbic acid (Sigma 49752) + 1mM HEPES•NaOH pH 7.4 (treated) and grown for an additional 12 hours before harvesting. This duration of treatment was chosen based on the observation of that accumulation of 5-hmC at methylated promoters peaks as early as 12 hours after starting treatment but returns to baseline over the following 60 hours of treatment (Blaschke et al., 2013). Cells were harvested by dissociation with TrypLE Express (Gibco), pelleted (300rcf, 5min, 4°C), washed once with 1X DPBS (with $Mg^{2+}/Ca^{2+}$), pelleted again, homogenized in 0.5 mL TRIzol Reagent, flash frozen in liquid $N_2$, and stored at -80°C.

Global changes in hmC levels were queried from total genomic DNA (gDNA) isolated from *Wdr76*$^{-/-}$ and parent E14 WT mESCs grown similarly as above in triplicate per condition and treated with 100 µg/mL 2-Phospho-L-ascorbic acid for 24 hours, with untreated cells grown in parallel under identical conditions. Following dissociation with TrypLE Express, cells were pelleted and resuspended in 1X PBS (with $Mg^{2+}/Ca^{2+}$), sonicated for 30 minutes in a bath sonicator (Branson), heated to 70˚C for 10 minutes, and finally vortexed at max speed for 1 minute. An equal volume of SPRI beads (GE SpeedBead magnetic carboxylate modified particles in 10mM Tris•HCl pH 8.0, 1mM EDTA, 20% (w/v) PEG 8k, 2M NaCl, 0.055% (v/v) Tween-20) was then added to the lysed suspension. Beads were washed twice with 70% ethanol and gDNA was eluted in 40 µL 0.1 x TBE. gDNA was subjected to blotting with an hmC-specific antibody, as described (Blaschke et al., 2013). Briefly, 1 µg of gDNA was diluted in 90 µL 6x SSC, to which 10 µL of 1M NaOH was added. Samples were heated at 90˚C for 10 minutes, then neutralized with the addition of cold $NH_4OAc$ to a final concentration of 1M. Samples were serially diluted into the wells of a Schleicher and Schuell dotting apparatus with vacuum manifold onto a nitrocellulose membrane equilibrated in 2x SSC such that the highest concentration point represents 100 ng. The wells were washed twice with two sample volumes 2x SSC. The membrane was then briefly dried at 80ºC for 5 minutes, and crosslinked at 254 nm at an energy or 120,000 µJ/$cm^2$ in a Stratalinker. The membrane was blocked in 1% ECL Advance blocking agent (GE Healthcare) in 1x TBS-T for 30 minutes at room temperature. The membrane was then incubated in a 1:5000 dilution of 5-hydroxymethylcytosine polyclonal antibody (Active Motif) overnight at 4˚C. The membrane was then washed three times for ten minutes each in 1x TBS-T, and incubated in a secondary antibody (1:10,000 dilution goat anti-rabbit IgG-HRP, Cell Signaling) for 1 hour at room temperature. The membrane was washed again in 1x TBS-T three times for ten minute each prior to visualization with ECL Advance reagents (GE Healthcare, Lumigen). Representative dot blot titrations for the cell lines examined are presented in Figure S6F.

#### RNA isolation and RT-qPCR

Homogenized mESC samples stored in TRIzol Reagent were thawed, mixed thoroughly with 1/5 volume chloroform, and the organic and aqueous phases were clarified by centrifugation (16,000rcf, 15min, 4ºC). The aqueous layer was cleanly removed, mixed with an equal volume of 100% ethanol, and applied to a Zymo RNA Clean & Concentrator-25 column. Columns were washed with 400 µL RNA Wash Buffer and then treated with 6 units RNase-free DNase I (Thermo Scientific) in 80 µL (10mM Tris-HCl pH 7.5, 2.5mM $MgCl_2$, 0.1mM $CaCl_2$) at room temperature by gently spinning the DNaseI solution into the column (50xg, 10sec) and letting sit for 15 minutes. Flow-through from this step was collected, mixed with two volumes of RNA Binding Buffer and three volumes of 100% ethanol, then loaded back onto its respective column. The remaining column purification steps were performed according to the manufacturer's instructions.

Reverse transcription of total RNA isolated from mESCs was performed in 20 µL reactions using 1 µg total RNA with 100 ng random hexamers (IDT) and 100 U MMLV-HP Reverse Transcriptase (Epicentre) according to manufacturer's instructions. After RT, the RNA was degraded by 100 mM KOH + 13.3 mM Tris base (final concentration) and incubated at 95 °C for 10 min. The pH was then adjusted to ~8.0 using 150 mM HCl, and samples were diluted to 300 µL final volume with 10 mM Tris-HCl pH 8.0. Real time quantitative PCR (RT-qPCR) reactions were performed in technical 10 µL triplicate with Power SYBR Green PCR Master Mix (Applied Biosystems) and 4 µL cDNA per reaction, with 250 nM each primer on a Bio-Rad CFX384 instrument.

The choice of reference gene for RT-qPCR normalization was determined empirically by evaluating the relative stability of 18S rRNA, GAPDH, PGK1, TBP, and ACTB via the geNorm *M* value (Vandesompele et al., 2002) parameter (qbase+ software, Biogazelle). TBP and ACTB were the highest-ranked candidates. All RT-qPCR target gene measurements were performed with TBP and ACTB reactions on the same plate, though we observed no significant difference between relative expression levels ($2^{\Delta Ct}$) calculated with normalization to the arithmetic mean Ct value of only TBP or ACTB, or to the geometric mean Ct value of both TBP and ACTB. For simplicity of uncertainty calculation, we chose arithmetic mean TBP normalization for figure display, as the raw TBP Ct values were closer to those of the target genes of interest being queried than raw ACTB Ct values. For validation of RNA-seq, independent experiments n=3: Grb10, Nanog, Chd1L, Tle1, Ncoa1, Gata6, Zdbf2, Cenpf, and Mki67; n=4: Phlda, Slc11a1, Nup210, Apobec2, Serpinh1, Yipf3, Des, Wdr26, Plagl1, Peg10, Meg3, Peg3, Kit, Tet1, and Tet2.

Fold changes ($2^{\Delta\Delta Ct}$) of gene expression in *Wdr76*$^{-/-}$ mESC were calculated relative to the parent E14 WT line, or for L-ascorbate-treated samples relative to the untreated control, using the arithmetic mean of all independent experiment samples' $2^{\Delta Ct}$ value for each sample type. Throughout figures displaying relative fold changes, the standard error of the mean (s.e.m.) for

*Wdr76*$^{-/-}$ samples includes the propagated uncertainty of the mean WT expression level used for calculating fold changes. Computed s.e.m. only includes variance across independent experiment samples' computed $2^{\Delta Ct}$ values and does not include variance from qPCR technical replicates. Significance testing was performed in R using Welch's two-tailed *t*-test by comparing the distribution of independent experiment samples' computed $2^{\Delta Ct}$ values between the appropriate sample types.

#### Analysis of WDR76 expression in patients using microarray

For analysis of the expression of WDR76 mRNA in AML patient samples, our previously published datasets GSE34184 and GSE30285 were used which contain a total of 100 human AML and nine normal BM samples (including three each of CD34+ hematopoietic stem/progenitor, CD33+ myeloid, and mononuclear cell (MNC) samples) (Huang et al., 2013). The dataset were generated by use of Affymetrix GeneChip Human Exon 1.0 ST arrays (Affymetrix).

#### siRNA transfection

THP-1 cells were maintained in RPMI-1640 medium (Life Technologies) containing 10% FBS, 1% HEPES and 1% penicillin-streptomycin. For MonoMac-6 (MM6) cells, 2 mM L-Glutamine, 1×Non-Essential Amino Acid, 1 mM sodium pyruvate, and 9 µg/ml insulin (Invitrogen) were additionally added to the medium. ON-TARGETplus Human WDR76 siRNA (L-014509-00-0005) was purchased from Dharmacon (GE Healthcare) and transfected into MM6 and THP1 cells at a final concentration of 20 nmol/L or 50 nmol/L using the 4D-Nucleofector™ System (Lonza, Cologne, Germany).

#### Plasmid construction and shRNAs

Mouse Wdr76 coding region was PCR-amplified from mouse cDNA with an N-terminal Flag-tag and subcloned into the MSCV-PIG retroviral vector using XhoI and EcoRI sites. The triple positive charge mutant (3S) was generated using the wildtype Wdr76 vector as template according to the protocol from the Multipoints Mutagenesis Kit (D407, Takara). The three primers used to introduce mutations are as follows: mut-1, GTTTTTTAATTCATCTTTGGAAAGCATAAGAACTGTTCATGTCCACC; mut-2, TTGACTGAGCACTCGAGCAGCATTGCCTCTGCATATTTC; mut-3, TGGACCACCCAAGCCGGGTGGAAGTCTTCCAC, where gray shades show residues being mutated to Serine, and nucleotides in red indicate the mutation sites.

For shRNAs targeting mouse Wdr76, GIPZ clones (shWdr76#1: V3LMM_456053; shWdr76#2: V3LMM_456057) were purchased from GE Healthcare.

#### Cell growth and apoptosis assays

Cell growth was assessed by MTT (G4000, Promega, Madison, WI) following the manufacturer's instructions. Briefly, 48 hours post transfection, cells were seeded on 96-well plate in triplicates at the density of 5000-10000 cells/100 µL. Dye solution was added at indicated time points and incubated at 37°C for 4 hours before stopping the reaction. The absorbance at 570nm (with reference at 630nm) was read on a microplate reader the next day.

For apoptosis assays, FITC Annexin V apoptosis Detection Kit 1 (BD Biosciences, San Diego, CA) was used for staining cells before they were analyzed in a BD LSRFortessa analyzer (BD Biosciences).

#### Virus infection and colony-forming assays (CFA)

These assays were conducted as described previously (Huang et al., 2013) with some modifications. Briefly, retroviruses or lentiviruses (shRNAs targeting Wdr76) were produced in

293T cells by co-transfection of individual expression construct with the pCL-Eco packaging vector (IMGENEX, San Diego, CA) or the pMD2.G:pMDLg/pRRE:pRSV-Rev packaging mix (individually purchased from Addgene), respectively. The virus particles were harvested at 48 and 72 hours after transfection.

Bone marrow (BM) cells were collected from 5- to 7-week-old B6.SJL (CD45.1) wild-type mice, and BM progenitor (i.e., lineage negative, Lin-) cells were enriched with the Mouse Lineage Cell Depletion Kit (Miltenyi Biotec). BM progenitor cells were then co-transduced with freshly collected MLL-AF9-MSCVneo (MLL-AF9) along with Wdr76 retroviruses (MSCV-PIG (vector), Wdr76-MSCV-PIG (Wdr76), or Wdr76-mut-MCSV-PIG (3S mut Wdr76)) or shWdr76 lentiviruses as indicated through two rounds of "spinoculation". Cells were then plated into ColonyGEL methylcellulose medium (ReachBio, Seattle, WA) supplied with 10 ng/ml of murine recombinant IL-3, IL-6, GM-CSF and 30 ng/ml of murine recombinant SCF, along with 1.0 mg/ml of G418 (Gibco BRL, Gaithersburg, MD) and 2.5 μg/ml of puromycin (Sigma-Aldrich). Cultures were incubated at 37°C in a humidified atmosphere of 5% $CO_2$ for 6 to 7 days. Serial replating was performed by collecting colony cells and replated them in methylcellulose medium every 6-7 days. Colony numbers were counted for each passage.

##### Mouse bone marrow transplantation (BMT)

These assays were conducted as described previously (Huang et al., 2013) with some modifications. Briefly, colony cells were collected from the colony-forming assays and transplanted via tail vein injection into lethally (900 cGy, 96 cGy/min, γ-rays) irradiated 8- to 9-week-old C57BL/6 (CD45.2) recipient mice. For each recipient mouse, $0.15\times 10^6$ donor cells from CFA assays and a radioprotective dose of whole bone marrow cells ($1\times 10^6$) freshly harvested from a C57BL/6 (CD45.2) mouse were transplanted. Leukemic mice were euthanized by $CO_2$ inhalation when they showed signs of systemic illness. Spleen and thymus from leukemic mice were collected at the time of sacrifice and weighted (Figure S7H-I). BM cells were isolated from both tibia and femur, and 50,000 cells were resuspended in 200 μL of cold MACS Buffer (1xPBS supplemented with 2 mmol/L EDTA and 0.5% BSA) and loaded for cytospin preparation. BM cytospin and blood smear slides were stained with Wright-Giemsa (Polysciences) (Figure S7G).

##### RNA extraction and qPCR

Total RNA was isolated using the miRNeasy mini kit (Qiagen) and quantified by UV spectrophotometry. For analysis of mRNA expression, 200-500 ng of RNA was reverse-transcribed into cDNA in a total reaction volume of 10 μL with the QuantiTect Reverse Transcription Kit (Qiagen). Quantitative real-time PCR analysis was then performed with 0.5 μL diluted cDNA (with 2.5-fold dilution) using Maxima SYBR green qPCR master mix (Thermo Fisher Scientific) on the QuantStudio 7 Flex PCR system (Thermo Fisher Scientific). GAPDH or ACTB was used as endogenous control (Figure S7D-F).

##### Western blots of leukemia lines

Cells were washed twice with ice-cold PBS and lysed using 1x SDS buffer (100 μL for $1\times10^6$ cells). Equal volumes of lysates were loaded and separated by 10% SDS-PAGE and transferred to polyvinylidene fluoride membranes. Membranes were blocked with 5% non-fat milk (Bio-Rad), incubated sequentially with primary and secondary antibodies and detected by immunoblotting with the Pierce ECL Western Blotting Substrate (Thermo Fisher Scientific). Antibodies used for Western blotting were as follows: WDR76 (ab108149, Abcam), β-Actin (3700, Cell Signaling Technology). β-Actin was used as a loading control (Figure S7E).

#### QUANTIFICATION AND STATISTICAL ANALYSIS

To align reads, a reference genome was first created, consisting of the human genome (GRCh38/hg38) or the mouse genome (mm9) appended respectively by the sequences of each of the nucleosome standard barcodes. Reads were then mapped to the appropriate reference genome using Bowtie2 using the sensitive pre-set and end-to-end alignment options (Langmead and Salzberg, 2012). Using SAMTools (Li et al., 2009), any reads with a map quality < 20 were discarded to prevent low-quality reads from impacting downstream analyses.

Normalized BigWigs of depth normalized coverage and BEDGraphs of genome coverage versus input were then generated using Deeptools (Ramírez et al., 2016). Peak calling was conducted for HA-WDR76 IP using Macs2 (Zhang et al., 2008) versus sequenced input.Profiles of ChIP and hmC distributions about features including gene bodies, enhancers, and TADs were generated using HOMER annotatePeaks (Heinz et al., 2010).

Bedtools Fisher or “phyper” in R were used to compute two-tailed Fisher Exact testing, as indicated in captions (Quinlan and Hall, 2010).

Statistical details of experiments can be found in the relevant figure legends. Linear correlations with R were forced through origin for more appropriate slope comparison.

#### DATA AND SOFTWARE AVAILABILITY

The ChIP- and RNA-seq reported in this paper have been deposited to the Gene Expression Omnibus (GEO) with the accession GSE108832.

#### SUPPLEMENTAL EXCEL TABLE AND VIDEO LEGENDS

**Table S1, Sequencing and Alignment Statistics.** Provided as accompanying Excel file.

**Table S2, Primer Table.** Primers used for qPCR. Provided as accompanying Excel file.
