## Supplementary Table 1 for "WDR76 promotes MLL-rearranged leukemia via selective recognition of 5-hydroxymethylcytosine in DNA"

### mESC RNA-seq statistics

| Sample | index primer set | Sample ID | index | index number | PF Clusters | % of the lane | % perfect barcode | % one mismatch barcode | Yield (Mbases) | % PF clusters | % >= Q30 bases | mean quality score | raw reads | reads mapped | % mapped | multiple alignments | % multi mapped | happy spliced reads |
|---|---|---|---|---|---|---|---|---|---|---|---|---|---|---|---|---|---|---|
| WT-1 | NEB NEXT (E7335) | AR-KM1-1 | TGACCA | 4 | 65,013,956 | 17.1 | 99.2 | 0.8 | 3,251 | 100 | 97.3 | 39.5 | 65013956 | 56704073 | 87.20% | 9338540 | 16.50% | 129919 |
| WT-2 | NEB NEXT (E7335) | AR-KM1-2 | GCCAAT | 6 | 66,469,714 | 17.5 | 99.1 | 0.9 | 3,323 | 100 | 97.3 | 39.5 | 66469714 | 59766737 | 89.9 | 9721540 | 16.3 | 126382 |
| WT-3 | NEB NEXT (E7335) | AR-KM1-3 | CTTGTA | 12 | 58,043,257 | 15.3 | 99.2 | 0.9 | 2,902 | 100 | 97.3 | 39.5 | 58043257 | 51173757 | 88.2 | 7867246 | 15.4 | 120314 |
| 3.1 KO-1 | NEB NEXT (E7335) | AR-KM1-4 | CGATGT | 2 | 76,990,270 | 20.3 | 99.2 | 0.8 | 3,850 | 100 | 97.3 | 39.5 | 76990270 | 66956626 | 87 | 9634477 | 14.4 | 134279 |
| 3.1 KO-2 | NEB NEXT (E7335) | AR-KM1-5 | CAGATC | 7 | 59,857,944 | 15.8 | 99.2 | 0.8 | 2,993 | 100 | 97.4 | 39.5 | 59857944 | 52269381 | 87.3 | 5848439 | 11.2 | 127865 |
| 3.1 KO-3 | NEB NEXT (E7335) | AR-KM1-6 | GATCAG | 9 | 50,296,223 | 13.3 | 99.3 | 0.7 | 2,515 | 100 | 97.4 | 39.5 | 50296223 | 43974337 | 87.4 | 5768014 | 13.1 | 126100 |

### K562 RNA-seq statistics

| Sample | index primer set | Sample | index | index number | PF Clusters | % of the lane | % perfect barcode | % one mismatch barcode | Yield (Mbases) | % PF clusters | % >= Q30 bases | mean quality score | raw reads | reads mapped | % mapped | multiple alignments | % multi mapped | happy spliced reads |
|---|---|---|---|---|---|---|---|---|---|---|---|---|---|---|---|---|---|---|
| NS-1 | NEB NEXT (E7335) | KM2-1 | TGACCA | 4 | 48,083,324 | 12.3 | 97.7 | 2.3 | 2,404 | 100 | 97.9 | 39.7 | 48083324 | 47088938 | 97.9 | 8420105 | 17.9 | 129623 |
| NS-2 | NEB NEXT (E7335) | KM2-2 | ACAGTG | 5 | 81,555,966 | 20.9 | 97.5 | 2.5 | 4,078 | 100 | 97.7 | 39.6 | 81555966 | 79258838 | 97.2 | 11297102 | 14.3 | 138450 |
| NS-3 | NEB NEXT (E7335) | KM2-3 | TAGCTT | 10 | 72,342,091 | 18.6 | 97.4 | 2.6 | 3,617 | 100 | 97.6 | 39.6 | 72342091 | 70315979 | 97.2 | 10486121 | 14.9 | 127420 |
| KDgRNA1 | NEB NEXT (E7335) | KM2-4 | GCCAAT | 6 | 67,830,362 | 17.4 | 97.6 | 2.4 | 3,392 | 100 | 97.5 | 39.5 | 67830362 | 65187603 | 96.1 | 8867231 | 13.6 | 129930 |
| KDgRNA2 | NEB NEXT (E7335) | KM2-5 | CAGATC | 7 | 56,763,525 | 14.6 | 97.5 | 2.5 | 2,838 | 100 | 97.8 | 39.6 | 56763525 | 55747109 | 98.2 | 7630615 | 13.7 | 128601 |
| KDgRNA5 | NEB NEXT (E7335) | KM2-6 | CTTGTA | 12 | 58,418,316 | 15.0 | 97.7 | 2.3 | 2,921 | 100 | 97.6 | 39.6 | 58418316 | 56484092 | 96.7 | 7205769 | 12.8 | 124773 |

### ChIP-seq statistics

| Sample | index primer set | Sample | index | index number | PF Clusters | % of the lane | % perfect barcode | % one mismatch barcode | Yield (Mbases) | % PF clusters | % >= Q30 bases | mean quality score | raw reads | reads mapped | % mapped | multiple alignments | % multi mapped | RPM scalar |
|---|---|---|---|---|---|---|---|---|---|---|---|---|---|---|---|---|---|---|
| WT IP | NEB NEXT (E7335) | AR-KM-1 | TGACCA | 4 | 71244682 | 19.48 | 98.42 | 1.58 | 3,562 | 100 | 96.8 | 39.3 | 71244682 | 65197021 | 91.51% | 19699177 | 27.65% | 0.015338 |
| WT input | NEB NEXT (E7335) | AR-KM-4 | CGATGT | 2 | 91580660 | 25.04 | 98.23 | 1.77 | 4,579 | 100 | 97.0 | 39.4 | 91580660 | 82807969 | 90.42% | 25533862 | 27.88% | 0.012076 |
