## Supplementary Table 2 for "WDR76 promotes MLL-rearranged leukemia via selective recognition of 5-hydroxymethylcytosine in DNA"

**Table S2**
qPCR Primer sets used in Malecek et al.

| primer number | description | sequence |
|---|---|---|
| 1 | Grb10 ex3F | ctcaagaagtggtgcagtatc |
| 2 | Grb10 ex3R | atattaactcgtccgtggaaag |
| 3 | Phlda ex1F | gttctggaagtcgatcagc |
| 4 | Phlda ex1R | caccatcgtcaccaactatta |
| 5 | Slc22a18 ex18F | gatggtcatcactgatctgac |
| 6 | Slc22a18 ex18R | agtagggagccgaagataat |
| 7 | Chdl1 ex18F | ctagatcttcataagccaggtc |
| 8 | Chdl1 ex18R | acggttaccagtccttct |
| 9 | Kit ex2F | ccatccagcacaatcagagtta |
| 10 | Kit ex2R | ggtcttgaaagtccatctgaca |
| 11 | WDR26 ex13F | ctaacagtgccatcgtctgag |
| 12 | WDR26 ex13R | cacacgcgcacagtaaattg |
| 13 | Slc11a ex2F | tgagcaggcccagttat |
| 14 | Slc11a ex2R | gatcttctcactcaggtaggt |
| 15 | 18S rRNA F | gtaacccgttgaaccccatt |
| 16 | 18S rRNA R | ccatccaatcggtagtagcg |
| 17 | mGAPDH sen | tgacgtgccgcctggagaaa |
| 18 | mGAPDH asen | agtgtagcccaagatgcccttcag |
| 19 | mPGK1 sen | ctgactttggacaagctggacg |
| 20 | mPGK1 asen | gcagccttgatcctttggttg |
| 21 | cMyc F | acacaaactcgaacagcttc |
| 22 | cMyc R | accgttctccttactctcac |
| 23 | Dnmt1 F | tgaccgcttctacttcctc |
| 24 | Dnmt1 R | ccttccctttccctttgttc |
| 25 | Dnmt3a F | ggtgcttttgtgtcgagtg |
| 26 | Dnmt3a R | cttatgcccgcacatgtag |
| 27 | Dnmt3b F | tggagttcagtaggacagc |
| 28 | Dnmt3b R | ccttgccattcatgactacag |
| 29 | Klf4 F | gctgtggcaaaacctatacc |
| 30 | Klf4 R | cacagtggtaaggtttctcg |
| 31 | Nanog F | tctcctccattctgaacctg |
| 32 | Nanog R | ccattgctagtcttcaaccac |
| 33 | Oct3/4 Pou5f1 II F | cactcacatcgccaatcag |
| 34 | Oct3/4 Pou5f1 II R | acttgatcttttgcccttctg |
| 35 | Sox2 F | ccattaacggcacactgc |
| 36 | Sox2 R | cctcccaattcccttgtatc |
| 37 | Tet1 F | ggctttctggtgtactttcag |
| 38 | Tet1 R | tgcaccctacagtcaaattg |
| 39 | Tet2 F | ccagaagcaagaaaccaagg |
| 40 | Tet2 R | gagcaatgacagtagccag |
| 94 | Tle1 ex6F | ccgatgatggcattcaactct |

| | | |
|---|---|---|
| 95 | Tle1 ex6R | ttcaaagcatcaacaacaggtg |
| 96 | mNcoa1 F | GACTGATGATGACGTGCAGAAG |
| 97 | mNcoa1 R | ATCCATCCAAAGCCTCCAGAAG |
| 98 | mCenpf F1 | TGGAACAGGAACTTAAGCGGT |
| 99 | mCenpf R1 | CTTCGTATGTGGAACTCGGTG |
| 100 | mMki67 F1 | CCATCATTGACCGCTCCTTT |
| 101 | mMki67 R1 | TATCTTGACCTTCCCCATCAGG |
| 102 | mZdbf2 F | ACTAACCTGGAACAGCACTTGT |
| 103 | mZdbf2 R | ATATGGGTGGTGTCGCAGAA |
| 104 | mGata6 F | ACCGGTCATTACCTGTGCAATG |
| 105 | mGata6 R | GTGATGAAGGCACGCGCT |
| 108 | mSerpinh1 F | CTTTAAGCCACACTGGGATGAG |
| 109 | mSerpinh1 R | TCCGGTGCATCATCGTAACA |
| 110 | mApobec2 F | GAGAAGCTGAAAGAGCTGATCG |
| 111 | mApobec2 R | CTGTACTTCGACCACATAGCAGA |
| 112 | mNup210 F | TCAGGAGGCTGTCTATAAGAATGT |
| 113 | mNup210 R | TTCTGTAATCTTGCCCTGCCT |
| 114 | mYipf3 F | CCAGGTCCGAAGCAGGCTC |
| 115 | mYipf3 R | CAGCATGAGCGGTCCGTAGA |
| 116 | mTmem238 F | CGGGGACCTGCTCATCTACT |
| 117 | mTmem238 R | GTAGGCCGTAGTCACGCT |
| 118 | mPeg3 F | GCACCAGCCGAGGTCTCAAA |
| 119 | mPeg3 R | GTTGCGAGCCACATCCTTGA |
| 120 | mRps16 F | GCCCAAATTTATGCCATCCGA |
| 121 | mRps16 R | TGATCTCCTTCTTGGAGGCTT |
| 122 | mMeg3 F | AGGACTTCACGCACAACAC |
| 123 | mMeg3 R | TGCATTTTCTCCTCAGCCCTT |
| 124 | mPeg10 F | CTGCCTTGAAGACCTTCCTGA |
| 125 | mPeg10 R | GTCACGAAGCACACACGGAT |
| 126 | mPlagl1 F | TCCAAGTATAAGCTGATGAGACACA |
| 127 | mPlagl1 R | ACTTCTTGCCGCAATCGT |
| 128 | mCenpf F2 | GACTTGGAAAGTGTTGTTCGGTG |
| 129 | mCenpf R2 | GTCAGTGTCCTGCTTTGGTGT |
| 130 | mMki67 F1 | TCAGTGAAGATCTGTCAGGACTAACTGA |
| 131 | mMki67 R1 | GGTGATGGGCTCAGGTATGT |
| 132 | mDes F2 | CCTCAAGGGCACCAACGACT |
| 133 | mDes R2 | TCCTGGTACTCGCGCAGATG |
| 134 | gooda_grbF | gcaaaatcctgtgcatttgggc |
| 135 | gooda_grbR | caaagggccatgtgcattcc |
| 136 | phladapus_F | cggggcatggggaaagtgaa |
| 137 | phladapus_R | ggtcccgtgcgtttctgtag |
| 138 | OK_boss_plagl1F | cgccacctcacaccaattcc |
| 139 | OK_boss_plagl1R | gtaccatgcagggcgtttgg |
| 140 | mWDR76 exon11 qFORW | cgttcatgtttacgatgccaggttc |
| 141 | mWDR76 exon11 qREV | cagcacatgtagtcaccactctgttac |

| | | |
|---|---|---|
| 142 | mGRB10 intr11 qFORW | AAAGCCAAAGTGTGCCTGGT |
| 143 | mGRB10 intr11 qREV | ACATGGCAAGAGAACGGGAA |
| 144 | mGRB10 intr4 qFORW | TTGTCACAGAGGCTCCAAGA |
| 145 | mGRB10 intr4 qREV | TTAGCCCATCCAATTCATCCTG |
| 146 | mSLC11A1 intr4 qFORW | GGGTCAGGGCAAAATGTGTC |
| 147 | mSLC11A1 intr4 qREV | TGACCCTAACTCCAGTCCCT |
| 148 | mSLC11A1 intr9 qFORW | CCTAAGAGCCCCCAGGGTAA |
| 149 | mSLC11A1 intr9 qREV | GGGTGGGTTGACTTCCTGTC |
| 150 | mPEG10 intr1 qFORW | AGGCCTAACTGATGCCTTGAA |
| 151 | mPEG10 intr1 qREV | TCCAGGAAGTCCAATGGTGT |
| 152 | mActb Forw4 | CCACCATGTACCCAGGCATT |
| 153 | mActb Rev4 | AGGGTGTAAAACGCAGCTCA |
